# Evidence for conserved gene expression and biological processes operative in human podocytes and adult brain

**DOI:** 10.1101/2025.03.17.643629

**Authors:** Wasco Wruck, Chantelle Thimm, James Adjaye

## Abstract

**Background:** Podocytes are essential for the proper functioning of the glomerular filtration barrier and are characterized by their intricate structure which includes well-organized primary and secondary foot processes. The brain expresses podocyte-associated proteins such as Nephrin, and Synaptopodin which are well documented. Processing of information and inter-cellular communication between podocytes and neurons employ similar molecular mechanisms which include actin-based projections, adhesion molecules, and signaling pathways.

**Methods:** In this study, we analysed brain-associated biological processes, transcription factors and pathways significantly regulated in UdPodocytes, i.e. podocytes differentiated from SIX2-positive UdRPCs (urine-derived renal progenitor cells). We compared gene expression in iPSC-derived brain and kidney organoids with UdPodocytes and mapped the overlapping 344 genes to brain regions via the GTEX (Genotype-Tissue Expression) database and investigated their regulation induced by the mediator of the renin-angiotensin system-angiotensin II (ANG II)

**Results:** The protein interaction network of UdPodocytes genes associated with brain in GTEX contains modules for pre-synapse, post-synapse, endocrine processes and neural crest differentiation. We also found that the genes overlapping between brain and podocytes are also expressed in iPSC-derived kidney and brain organoids and could map the involved genes to all regions of the brain with the frontal cortex as the most enriched. The genes SUZ*12, NFKB1* and *PAX2* were the most significantly over-represented transcription factors. We independently confirmed the conserved expression of PAX6, KCNQ3, TUJ1,MAP2, TAU and ACTN4 in Udpodocytes as shown by Western blotting, Immunofluorescence and RT-PCR. We further investigated if the conserved genes are also regulated by ANGII. This unveiled genes down-regulated upon ANGII-stimulation of Udpodocytes to be associated with axon guidance, Calcium, Hippo and cGMP-PKG signaling as over-represented pathways.

**Conclusion:** In conclusion, we have identified 344 genes with overlapping expression in brain and podocytes. These are mainly associated with synaptic signaling and cell projections. This implies that human urine-derived SIX2-renal progenitor cells differentiated into podocytes can serve as a platform for dissecting and understanding the relevance of conserved biological processes in podocytes which are currently annotated as neuron projection, axons, neurogenesis and synaptic signaling.

## Introduction

The central nervous system (CNS) and the kidneys are very closely connected, for example, the CNS provides afferent impulses that regulate and influence renal blood flow, the glomerular filtration rate (GFR) and the sodium balance [1] and Cao et al. report a kidney-brain neural circuit driving kidney damage and heart failure [2]. With respect to early development, podocytes and neurons are derived from distinct lineages, neurons arise from the ectodermal germ layer and podocytes from metanephric mesenchyme [3] developing from the intermediate part of the mesoderm germ layer [4].

The similarity between podocytes and neurons is largely determined by common features of primary processes operative in both cell types as well as the need to exchange signals across membranes in synaptic or synaptic-like constellations [5]. Processing of formation in podocytes and neurons use the same molecular machinery which include actin-based projections such as podocyte foot processes and dendritic spines [6]. These are organized in microtubules (MT) and intermediate filaments for thick processes and actin filaments for thin projections [6]. Further similarity resides in the cell-cell contacts in synapses and the slit diaphragm where adhesion molecules in neurons and podocytes belong to the same families such as Cadherins/Protocadherins, Ig-like proteins, Neurexin and Podocalyxin [7]. Synaptopodin (SYNPO) is an Actin-associated protein that regulates the cytoskeleton in podocytes. It is not only expressed in mature podocytes but also in parts of the cytoskeleton of postsynaptic densities (PSD) and their dendritic spines. Studies have shown that the expression of SYNPO is restricted to a sub-population of cells within the telencephalic synapse [8], [9].

Furthermore, communication between podocytes and other glomerular cells is mediated via signaling mechanisms involving neurotransmitters which are also found in synapses [7] such as the glutamate receptors NMDAR and GRM1. Within podocytes, Nephrin connects these glutamate receptors through scaffolding molecules, such as PSD95, to the actin cytoskeleton of the foot processes [7].

Numerous kidney-associated diseases can result in neuronal damage. A decrease in the glomerular filtration rate (GFR) due to age or disease leads to higher concentration of the filtered waste products in the blood and can cause uremia. This accumulation of uremic substances in the blood can exert toxic effects on neurons such as in uremic encephalopathy, uremic neuropathy and uremic myopathy [10]. Besides the uremic toxins, oxidative stress, inflammation, and impaired blood circulation and blood brain barrier are factors mediating neurological disorders in kidney diseases, particularly in chronic kidney disease (CKD). On the other hand, leakage in the glomerular filtration barrier which leads to low concentrations of molecules such as Albumin in the blood and high concentrations in the urine are another source of kidney damage such as in diabetic nephropathy.

The similarity between podocytes and neurons manifests directly in diseases associated with calcium signaling. Deregulated or non-functional proteins from the TRPC family are associated with focal segmental glomerulosclerosis (FSGS-TRPC6) and TRCP5 induces pathologically motile podocytes while in brain the loss of TRCP5 can result in deficits in gait and motor coordination [7]. Another disease affecting neurons and podocytes, Charcot-Marie-Tooth neuropathy, often co-occurring with FSGS, is associated with mutations in *INF2* which is involved in regulation of the actin and microtubule cytoskeleton [11]. Interestingly, apart from podocytes, brain dysfunction in tubular and tubulointerstitial diseases is more often seen in disorders of water handling such as Bartter and Gitelman syndromes [12].

In this study, we further investigated similarities between a set of genes expressed in brain and podocytes with respect to neuronal-associated gene ontologies operative in urine-derived podocytes [13]. We also confirmed expression in iPSC-derived brain and kidney organoid cultures and revealed their responsiveness to ANGII-stimulation. Signalling pathways and genes regulatory networks suggest common functionality in brain and podoytes.

## Methods

### Cell culture conditions

Urine-derived renal progenitor cells (UdRPCs) from distinct individual males with ages 48 and 51 years (UM48, UM51) and 21 and 27year old females (UF21, UF27), were isolated following the protocol detailed by Rahman et al. [14]. These cell cultures were maintained on 6- or 12-well plates uncoated at 37 °C in a hypoxic environment. Cells were grown in Proliferation Medium (PM), consisting of a 1:1 mixture of DMEM high-glucose (Gibco) and keratinocyte growth basal medium (Lonza, Basel, Switzerland), supplemented with 5% fetal bovine serum (Gibco), 0.5% non-essential amino acids (Gibco), 0.25% Glutamax (Gibco), and 0.5% penicillin-streptomycin (Gibco).

For differentiation into podocytes, cells were seeded at a low density (50,000 cells per 6-well plate) on a Corning® Collagen I (Merck, Darmstadt, Deutschland) coated six or 12-Well plate and cultured for 24 hours in PM. The following day, the medium was replaced with Advanced RPMI 1640 (Gibco) supplemented with 0.5% fetal bovine serum, 1% penicillin-streptomycin, and 30 μM retinoic acid (Sigma-Aldrich Chemistry, Steinheim, Germany). After seven days, cells exhibited the characteristic morphology of podocytes. Furthermore a human SV40-temperature sensitive immortalized podocyte cell line (AB 8/13) [15] was used as an established reference. The cells were first cultured at 33 °C in RPMI 1640 (Gibco). To induce podocyte differentiation and maturation the, cells were seeded on Corning® Collagen I (Merck, Darmstadt, Deutschland) coated dishes and cultured at 37 °C in Advanced RPMI 1640 (Gibco) supplemented with 0.5% fetal bovine serum, 1% penicillin-streptomycin, and 30 μM retinoic acid (Sigma-Aldrich Chemistry, Steinheim, Germany).

### Immunofluorescence-based detection of protein expression

Cells were fixed with 4% paraformaldehyde for 15 minutes at room temperature (RT) and subsequently washed three times with PBS. Afterwards the cells were blocked using 3% BSA in PBS. Primary antibodies (detailed in Table S5) were applied, and the cells were incubated overnight at 4 °C. The next day, the cells underwent a single washing step with Triton X-100 in PBS, followed by two washes with PBS. Secondary antibodies (detailed in Table S5) together with Hoechst as a nuclear stain, were applied for 2 hours at RT. Fluorescence imaging was performed using an LSM700 microscope (Carl Zeiss).

### Western Blotting

Podocytes were lysed using RIPA buffer (Sigma-Aldrich Chemistry) supplemented with 5 M NaCl, 1% NP-40, 0.5% DOC, 0.1% SDS, 1 mM EDTA, and 50 mM Tris (pH 8.0), along with freshly added protease and phosphatase inhibitors (10 μL/mL, Sigma-Aldrich). A total of 20 μg of protein from each lysate was resolved on a 10% SDS-PAGE gel and transferred onto an Immobilon-P membrane (Merck Millipore, Burlington, VT, USA). Membranes were incubated overnight at 4 °C with the primary antibody, followed by three washes with 0.1% Tween-20 in Tris-buffered saline. Secondary antibodies were then applied for 1 hour at room temperature. Protein visualization was performed using the Pierce™ ECL Western Blotting Substrate (Thermo Fisher Scientific, Massachusetts, USA), with both solutions mixed at a 1:1 ratio. Quantification of protein bands was conducted using ImageJ software. Further details regarding the antibodies used are available in Supplemental Table S5.

### RNA isolation

For RNA isolation from the various cell lines, the ZYMO Research Kit Direct-zol™ RNA Miniprep R20 was utilized following the manufacturer’s protocol. Cells were detached by adding Tryple E and incubated for 5 min. Afterwards, a centrifuging step was added 4 mins 1000 g. Cell pellet was resuspend in 350 μl volume of 95–100% ethanol (EtOH) and thoroughly mixed by pipetting. The resulting solution was transferred to a Zymo-Spin™ column and centrifuged at 12,000 g for 30 seconds at 4 °C. To degrade genomic DNA, the column was washed with RNA Wash Buffer and centrifuged again at 12,000 g for 30 seconds at 4 °C. Subsequently, the column was treated with DNA digestion mix (1:16 dilution of DNase I in DNA Digestion Buffer) for 15 minutes. Following digestion, the column was rinsed twice with RNA Prewash Buffer, each followed by centrifugation at 12,000 g for 30 seconds at 4 °C. A final wash with RNA Wash Buffer was performed, including a 2-minute centrifugation step.

After drying the column by centrifuging for 1 minute, it was transferred to a fresh RNase-free Eppendorf tube. RNA was eluted by adding 35 μL of DNase/RNase-free water to the column and centrifuging for 1 minute at 16,000 g. The eluted RNA samples were immediately placed on ice, and concentrations measured with a Nanodrop device.

### Semi-quantitative RT-PCR

1000 ng of RNA sample was used for reverse transcription using the TaqMan Kit following the manufacturer’s instructions. PCR was performed using the GoTaq® G2 Hot Start Polymerase Kit from Promega. For each sample, 1 μL cDNA (5 ng/μL) was mixed with 24 μL master mix. As a negative control, 1 μL RNAse-free water was used. As references, commercial RNA samples of Human kidney total RNA (Takara, #636529) and fetal brain (BioChain®) were used. PCR was performed using a thermal cycler under the following conditions, polymerase activation: 95 °C for 2 min; 35 cycles amplification: 95 °C for 30 s, 60 °C for 30 s, 72 °C for 30 s; final elongation: 72 °C for 5 min, and finally held at 4 °C. The PCR product was resolved in a 2% agarose gel by gel electrophoresis. *RPL0* was used as the housekeeping gene.The imaging device Fusion FX was used for visualization.

### Acquisition of brain and podocyte transcriptome data

Microarray datasets related to experiments conducted with brain cells and podocytes were downloaded from the public repository at the NCBI GEO (National Center for Biotechnology Information, Gene Expression Omnibus) (Table 1). In order to minimize technical variability only datasets from the same technical platform were employed. Kidney-related datasets are associated with two previous studies from our lab by Erichsen et al. (GSE171240) pertaining to Angiotensin-II-treated podocytes [13] and by Nguyen et al. (GSE186823) containing kidney organoids treated with the nephrotoxin Puromycin Aminonucleoside (PAN) [16]. Brain-related datasets are associated with a previous study from our lab pertaining to brain organoids [17] and with datasets associated with genes expressed in brain regions based on RNAseq data from the Genotype-Tissue Expression (GTEx) project [18].

**Table 1:** Brain and kidney transcriptome datasets from NCBI GEO and other sources used in this study.

| dataset | contents | brain/kidney region | platform | PubMedId |
| --- | --- | --- | --- | --- |
| GSE279611 | control | brain organoids | Affymetrix<br>Human Clariom-S | 35269426 |
| GTEx | tissue gene<br>expression | brain regions | RNA-Seq | 23715323 |
| GSE171240 | ANGII treated<br>podocytes | podocytes | Affymetrix<br>Human Clariom-S | 35406662 |
| GSE186823 | PAN treated kidney<br>organoids | kidney organoids | Affymetrix<br>Human Clariom-S | 35203286 |
|  |  | brain |  |  |
|  |  | kidney |  |  |

### Identification of brain-related biological processes in urine-derived podocytes

This study was initiated by the unexpected observation of a plethora of over-represented brain-associated GOs in transcriptomes of urine derived SIX2-positive renal progenitor cells -UdRPCs differentiated into podocytes and stimulated with ANGII [13]. In the gene expression data from our previous study by Rahman et al. [19] genes expressed in UdRPCs and UdPodocytes [13] were compared in a Venn diagram using the VennDiagram package [20] from the R/Bioconductor environment [21]. Gene expression was determined by a detection-p-value < 0.05 as described in our previous publication [22]. Genes exclusively expressed in the UdPodocytes but not in the UdRPCs were subjected to over-representation analysis via the GOstats package [23].

### Association of genes with whole brain and brain regions

Genes associated with tissues were downloaded from the GTEX portal (version GTEx_Analysis_v6_RNA-seq_RNA-SeQCv1.1.8_gene_median_rpkm.gct) [18] as median RPKM data from RNAseq experiments. From these genes with a median RPKM value five times greater than the mean of all tissues were considered specific for that tissue. Particularly for this study genes associated with brain regions were extracted from the dataset.

### Comparison of brain-related genes in urine-derived podocytes to kidney and brain organoids

Genes expressed in kidney as well as brain organoids were extracted from our previous studies based on iPSC-derived kidney [16] and brain organoids [17]. The gene-set expressed in common was compared to the genes we found above exclusively expressed in UdPodocytes in a Venn diagram via the R package VennDiagram [20]. The intersection set from the venn diagram was further refined in a pie-chart for brain regions associated with the contained genes in the GTEX dataset. The pie-chart of brain regions was drawn with the function “pie3D” from the R package “plotrix”. Analysis of transcription factors over-represented in the promoter regions of the intersection gene set was preformed via the EnrichR web tool [24]. Association of genes to GO terms were made via the package “org.Hs.eg.db” from the R/Bioconductor [21]. For determining transcription factors (TFs), the GO terms of category “Molecular Function” with the 20 most associated genes were extracted via ratios dividing the number of associated genes by the total gene number. TFs were taken from the GO terms nucleic acid binding, DNA binding, transcription regulator activity, sequence-specific DNA binding and DNA-binding transcription factor activity, RNA polymerase II-specific. The protein interaction network was constructed using these TFs and Biogrid interactions [25] as described in our previous publication [17].

### Comparison of neuron-related genes in urine-derived podocytes to brain-associated genes

Genes associated with whole brain, i.e. all brain regions, in the GTEX dataset were compared to the genes exclusively expressed in UdPodocytes in a Venn diagram via the R package VennDiagram [20]. The intersection gene set from the venn diagram was submitted to the STRING-DB web tool [26] to construct a protein interaction network which in turn was used as input for Cytoscape [27]. In Cytoscape a clustering for subnetworks was performed via the plugin MCODE [28]. The sub-networks were highlighted in the whole network and analyzed for enrichment via cytoscape-builtin functions. Most significant terms from the enrichment analysis were plotted as barcharts of negative log10 p-values indicating ratios of involved gene numbers to total gene numbers on a color scale via the R package ggplot2 [29].

## Results

### Urine-derived Podocytes develop projections with synapses

Urine-derived renal progenitor cells (UdRPCs) were derived from urine and differentiated into podocytes (UdPodocytes) according to the scheme illustrated in Figure 1a [13], [30]. Nephrin (NPHS1) plays a major role in podocyte development and is localised at the slit diaphragm (Figure 1b: NPHS1 - red staining, phalloidin - green staining in UdPodocytes). NPHS1 is expressed in developing brain [31] and we found it up-regulated in kidney organoids [16]. The green phalloidin staining shows the actin filament structure of the podocytes. UdPodocytes can sporadically develop axon-like projections with synapses connecting to other cells (Figure 1c, white arrows pointing to synapses).

**Figure 1:**
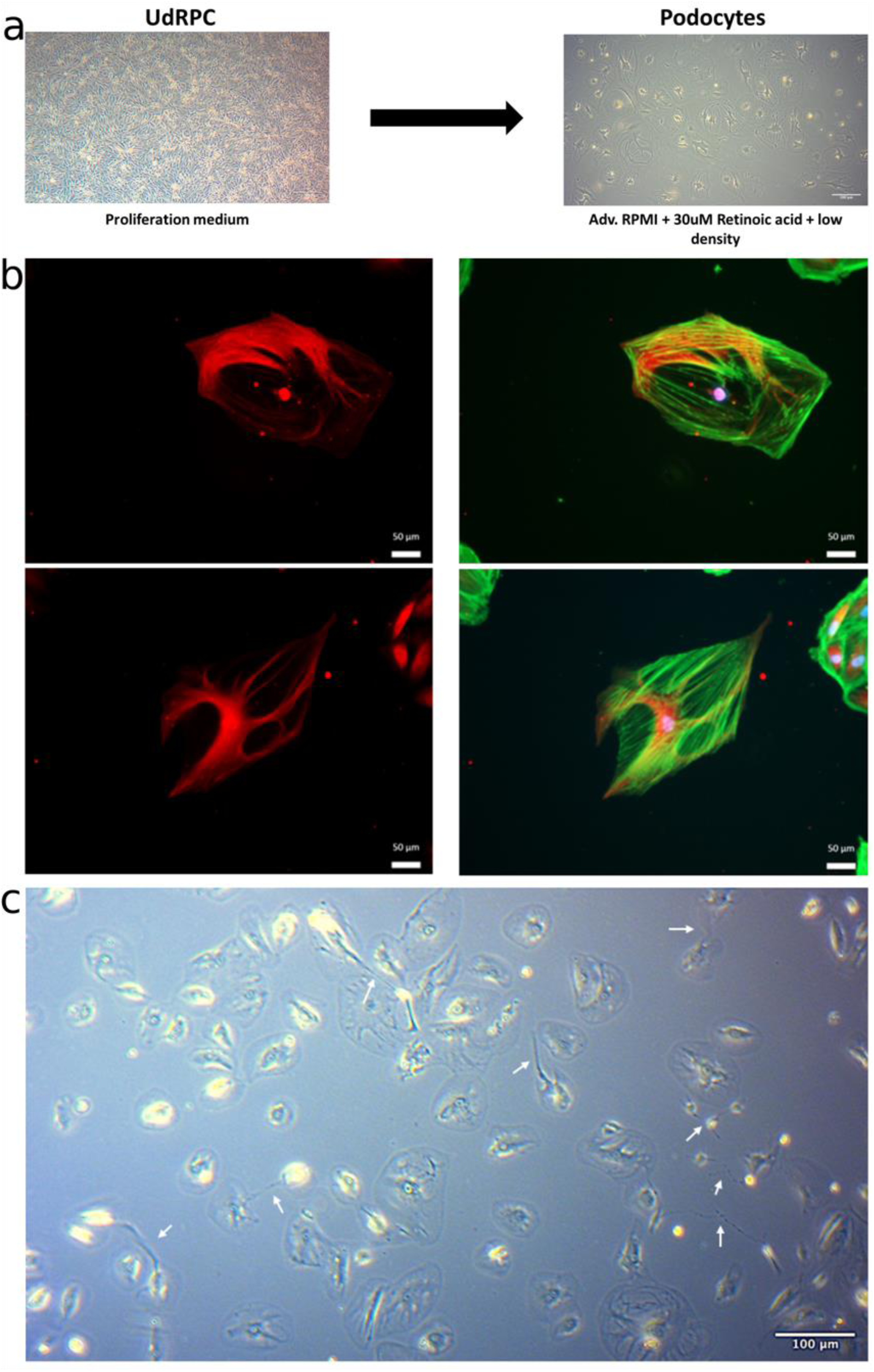
Podocytes develop projections with synapses. (a) Urine-derived renal progenitor (UdRPCs) were differentiated into UdPodocytes according to this scheme. (b) Nephrin (NPHS1) plays a major role in podocyte development and is found at the slit diaphragm. The staining here shows NPHS1 in red and phalloidin in green in UdPodocytes. (c) Podocytes derived from UdRPCs can develop axon-like projections with synapses inter-connecting other cells (arrows pointing at examples).

### Identification of brain-associated biological processes in urine-derived podocytes

This study was initiated by the unexpected observation of a plethora of over-represented brain related GOs in transcriptomes of UdRPCs differentiated into podocytes [13], [30]. A probable explanation for this result is that there is obviously more transcriptome data available of the brain than in the kidney and as such much more GO annotations exist for brain. Nevertheless, these results also demonstrate the similarity in cell-cell communication-associated molecular mechanisms between podocytes and brain. Figure 2 shows neuron-related GO terms extracted from a list of GOs significantly over-represented in urine-derived podocytes (UdPodocytes, 600 genes) extracted from a Venn diagram comparison of expressed genes between urine-derived renal progenitor cells (UdRPCs) and UdPodocytes (Figure 2a, Figure S1). The most significant Cellular Component (CC) is *excitatory synapse* (Figure 2b) the most significant Molecular Function (MF) is *neurotransmitter:sodium symporter activity*. The three most significant Biological Processes in the list of GO terms are *axon guidance, cell morphogenesis involved in neuron differentiation* and *neuron projection morphogenesis* (Figure 2d). A more detailed analyses of the genes associated with *axon guidance* revealed genes encoding Ephrins, Slits, and Semaphorins (Figure S2). These ligands and receptors are responsible for the regulation of the actin cytoskeleton on the one hand and on the other hand for attraction and repulsion. In summary, the GO terms can be categorized into groups related to neurons including amongst others *neurogenesis* and *neuron differentiation*, neuron projections including *neuron projection morphogenesis*, axons including *axon guidance* and *axon development* and synapses including *positive regulation of synapse assembly*.

**Figure 2:**
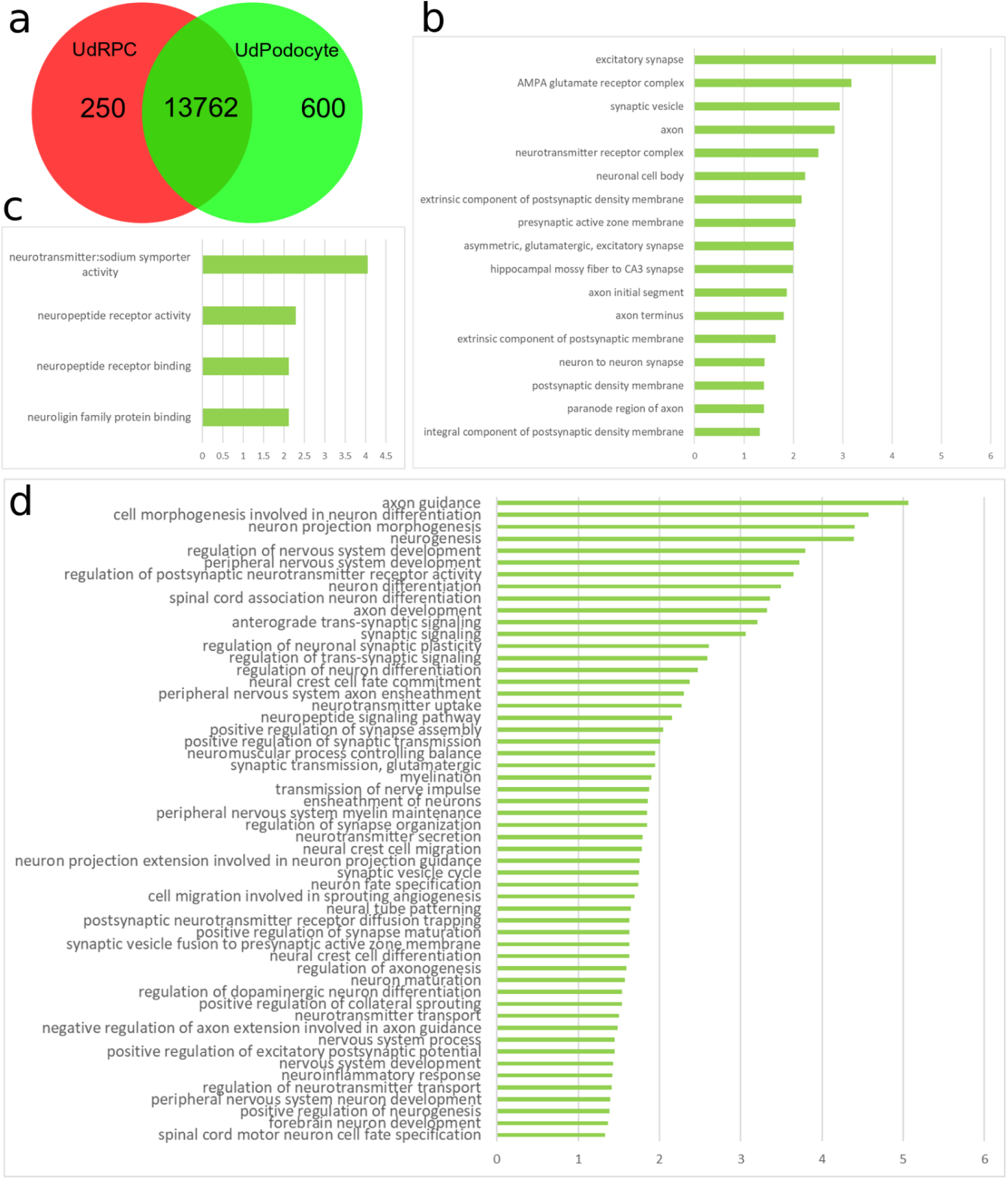
Brain-associated biological processes are enriched in Urine-derived Podocytes (UdPodocytes). Genes expressed exclusively in UdPodocytes in a Venn diagram comparison (a) with urine-derived SIX2-positive renal progenitor cells (UdRPCs) were subjected to gene ontology (GO) over-representation analysis and brain-related terms were extracted from them. From the resulting list of GOs (b) the cellular components, (c) the molecular functions and (d) the biological processes with a p-value less than 0.05 are plotted here with bars indicating their negative logarithmic (base 10) p-value.

### Comparison of brain-associated gene expression in urine-derived podocytes and in iPSC-differentiated brain and kidney organoids

We further compared the genes associated with neuron-related biological processes in urine-derived podocytes detected above with transcriptome data from brain and kidney organoids. Figure 3a shows a venn diagram comparing the genes expressed in kidney and brain organoids with the 600 genes identified to be exclusively expressed in UdPodocytes. The resulting intersection set of 344 genes were mapped to brain regions via associations retrieved from the GTEX database (Figure 3b) showing distribution over multiple brain regions with most hits and most significant over-representation in the frontal cortex (p=0.0006, hypergeometric test). GO analysis via the R package GOStats [23] was applied to detect over-represented GO terms in the 344 genes (Figure 3c). This revealed categories such as *cellular morphogenesis, neuron differentiation, synaptic signaling* and axo*n guidance*. In a follow-up REVIGO analysis the “Biological Process”-associated GO terms “*cell projection morphogenesis*” and “*regulation of postsynaptic neurotransmitter receptor activity*” were central hubs (Figure S3). Moreover, Figure 3d and Table S2g depict results of a Metascape analysis based on the 20 GO terms of the category “Molecular Function” with most matches of the 344 genes. (approximately 70% of the 344 genes are associated with binding activity while 25% are associated with catalytic, 13% with hydrolase and 10% with transporter activity) yielding similar results as Figure 3c. Over-representation analysis of the 344 genes with the EnrichR tool [24] revealed SUZ12, NFKB1 and PAX2 as most significantly over-represented transcription factors (Table S2c-e) and also transcription factors including PAX6, PAX7, SOX2 and GATA4 associated with TF-associated GO Molecular Functions *DNA-binding, transcription regulator activity, etc.* (Table S2f) which were partially validated via kidney expression in the Protein Atlas [32] (Table S2h). These validated proteins could be connected in a protein interaction network of Biogrid interactions using EGFR and XPO1 as major hubs (Figure S4).

**Figure 3:**
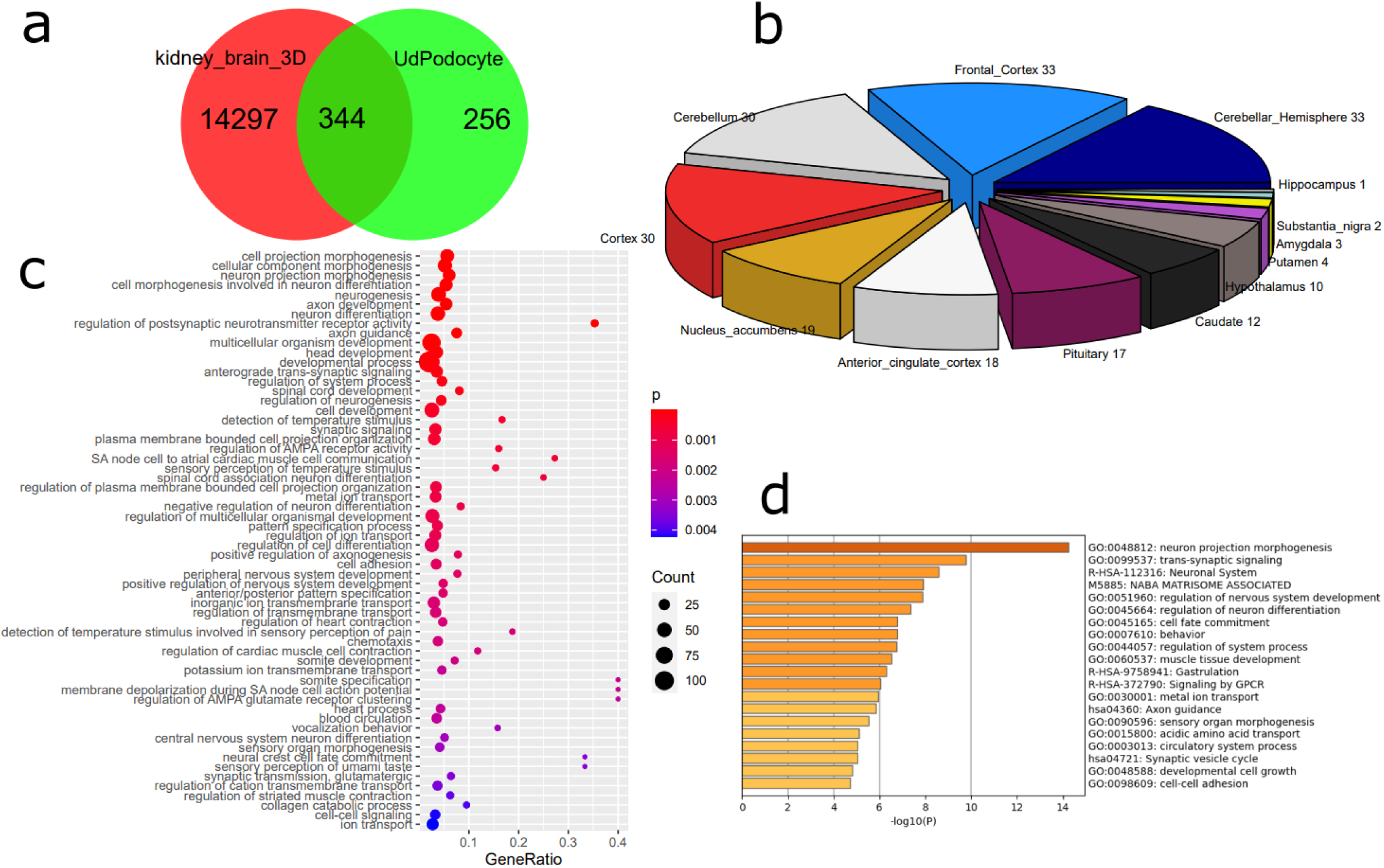
The overlap between urine-derived Podocytes (UdPodocytes) with brain and kidney organoids shows over-representation of the frontal cortex brain region and biological categories such as cellular morphogenesis, neuron differentiation, synaptic signaling and axon guidance. (a) Genes expressed in kidney and brain organoids were compared with the 600 genes identified as exclusively expressed in UdPodocytes resulted in an intersection of 344 genes. (b) The 344 genes map to multiple brain regions with most hits in frontal cortex. (c) Results of the GO analysis using the 344 genes expressed in common revealed biological terms from categories such as *cellular morphogenesis, neuron differentiation, synaptic signaling* and *axon guidance*. (d) Mapping genes to the GO: Molecular Function (GO-MF) top category shows that most of the 344 genes are associated with binding and transcription factor activity and submitting the 294 genes associated with the top 20 GO-MFs to Metascape analysis besides similar biological processes as in (c) reveals involvement of the extra-cellular matrisome.

### The protein interaction network of UdPodocytes genes associated with brain

Figure 4a shows the protein interaction network of the 130 genes from the intersection set of the venn diagram comparison of genes exclusively expressed in UdPodocytes and genes associated with brain in the GTEX database (Figure 4b). In a REVIGO analysis of the “Biological Process”-associated GO terms found over-represented in the 130 gene-set, “*neurogenesis*” and “*regulation of trans-synaptic signaling*” were central hubs (Figure S5). Furthermore, we submitted the 130 genes to the STRING online database, added no further interacting proteins but removed unconnected proteins. The STRING network was then loaded into Cytoscape and analysed with the MCODE plugin to identify clusters/subnetworks. In Figure 4a the most significant five subnetworks MCODE1-5 are highlighted. Bar charts in Figure 4c-g show biological terms found enriched in the subnetworks MCODE1-5 with the corresponding color. Subnetwork MCODE1 (Figure 4c, red) contains the proteins NTRK2, SNAP25, SYT1, BDNF, NRXN1 and NRXN3 and is most significantly enriched with the GO *presynapse*. Subnetwork MCODE2 (Figure 4d, blue) contains the proteins SYNDIG1, CNIH2, CACNG4, SHISA7, SHISA6 and CAMK2B and is most significantly enriched with the GO *postsynaptic density membrane*. Subnetwork MCODE3 (Figure 4e, green) contains the proteins CRH, GAL and KISS1 and is most significantly enriched with the GO *regulation of endocrine process*. Subnetwork MCODE4 (Figure 4f, violet) contains the proteins MYO7A, OTOG and STRC and is most significantly enriched with the GO *Sensory processing of sound by outer hair cells of the cochlea*. Subnetwork MCODE5 (Figure 4g, grey) contains the proteins GBX2, PAX7 and WNT1 and is most significantly enriched with the GO *Neural crest differentiation*. mRNA expression of *KCNQ3, PAX6* and *TUJ1* which are amongst the 130 selected genes were confirmed in four distinct podocyte cultures and the immortalized podocyte cell line (Figure S6, primers used Table S6) and protein expression was confirmed in the Human Protein Atlas (proteinatlas.org) [32] (Figure S7). The expression of RPL0 was measured as a housekeeping gene.

**Figure 4:**
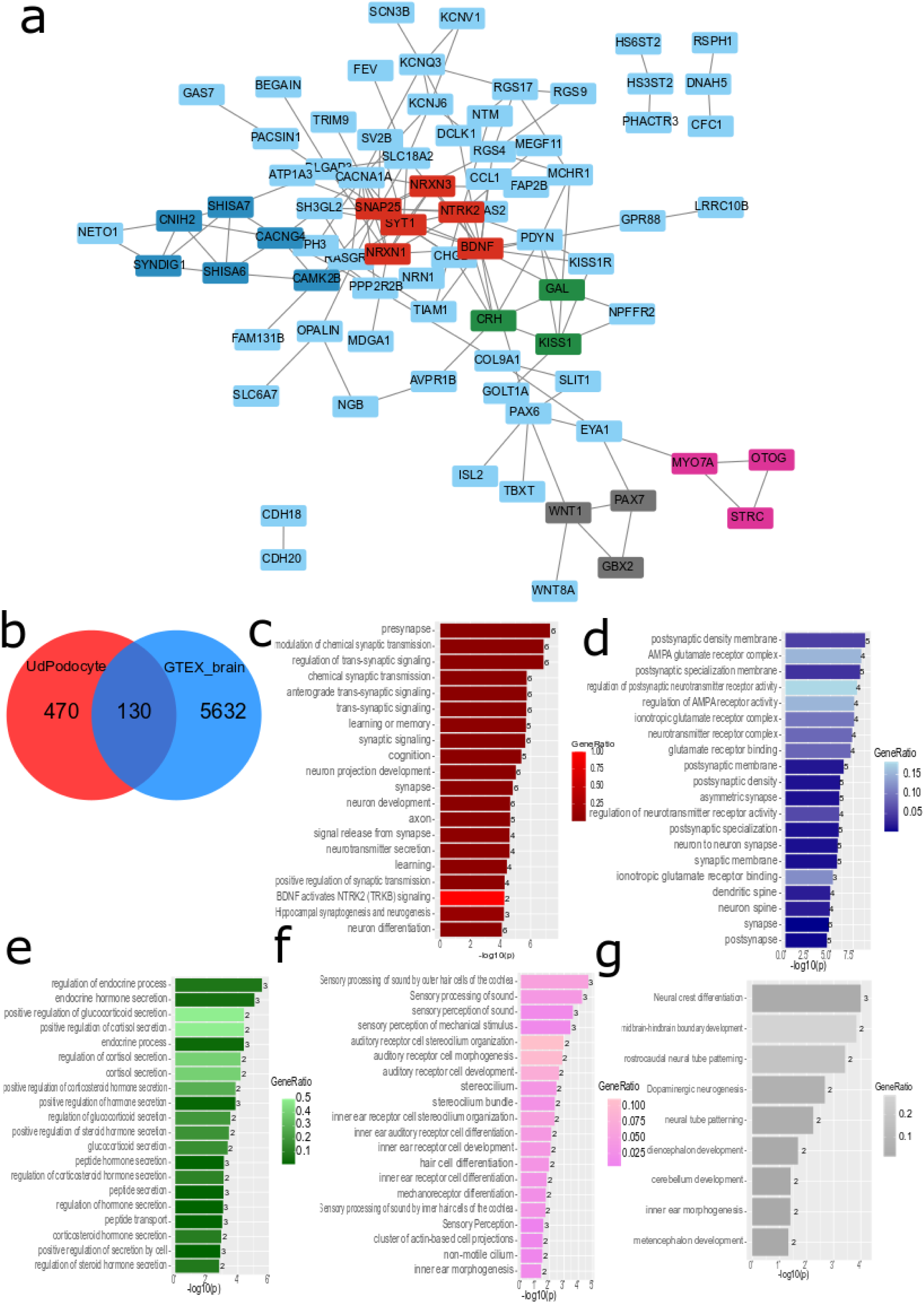
The protein interaction network of UdPodocytes genes associated with brain in GTEX contains modules for pre-synapse, post-synapse, endocrine processes, cochlea and neural crest differentiation. (a) STRING protein interaction network of 130 genes resulting from the venn diagram comparison of genes exclusively expressed in UdPodocytes and genes associated with brain in the GTEX database (b). Analysis of the STRING network in Cytoscape and MCODE yielded several subnetworks of which the most significant five MCODE1-5 are highlighted. Bar charts in panels (c-g) show biological terms found enriched in the subnetworks MCODE1-5 with the corresponding color. (c) In subnetwork MCODE1 *presynapse* is the most significantly enriched biological term. (d) In subnetwork MCODE2 *postsynaptic density membrane* is the most significantly enriched biological term. (e) In subnetwork MCODE3 *regulation of endocrine process* is the most significantly enriched biological term. (d) In subnetwork MCODE4 *Sensory processing of sound by outer hair cells of the cochlea* is the most significantly enriched biological term. (e) In subnetwork MCODE5 *Neural crest differentiation* is the most significantly enriched biological term.

### Expression of common brain associated proteins

In our previous manuscripts, we showed the capacity of urine-derived renal progenitors (UdRPCs) to differentiate into podocytes [13], [30], [19],[33]. Figure 5 depicts an immunofluorescence staining confirming the differentiation capacity of all five UdRPC preparations. Two podocyte markers were used, the cytoskeletal protein Alpha Actinin 4 (ACTN4) and the adhesion molecule Nephrin (NPHS1) (Figure 5 A+B). For investigation of common brain-associated proteins, β-Tubulin (TUBB) staining was also performed (Figure 5C). Furthermore, the expression of the podocyte cytoskeletal protein ACTN4 served as reference, TUBB, microtubule-associated TAU (MAPT) and Microtubule Associated Protein 2 (MAP2) were analyzed by Western blotting (Figure 6A-C).

**Figure 5:**
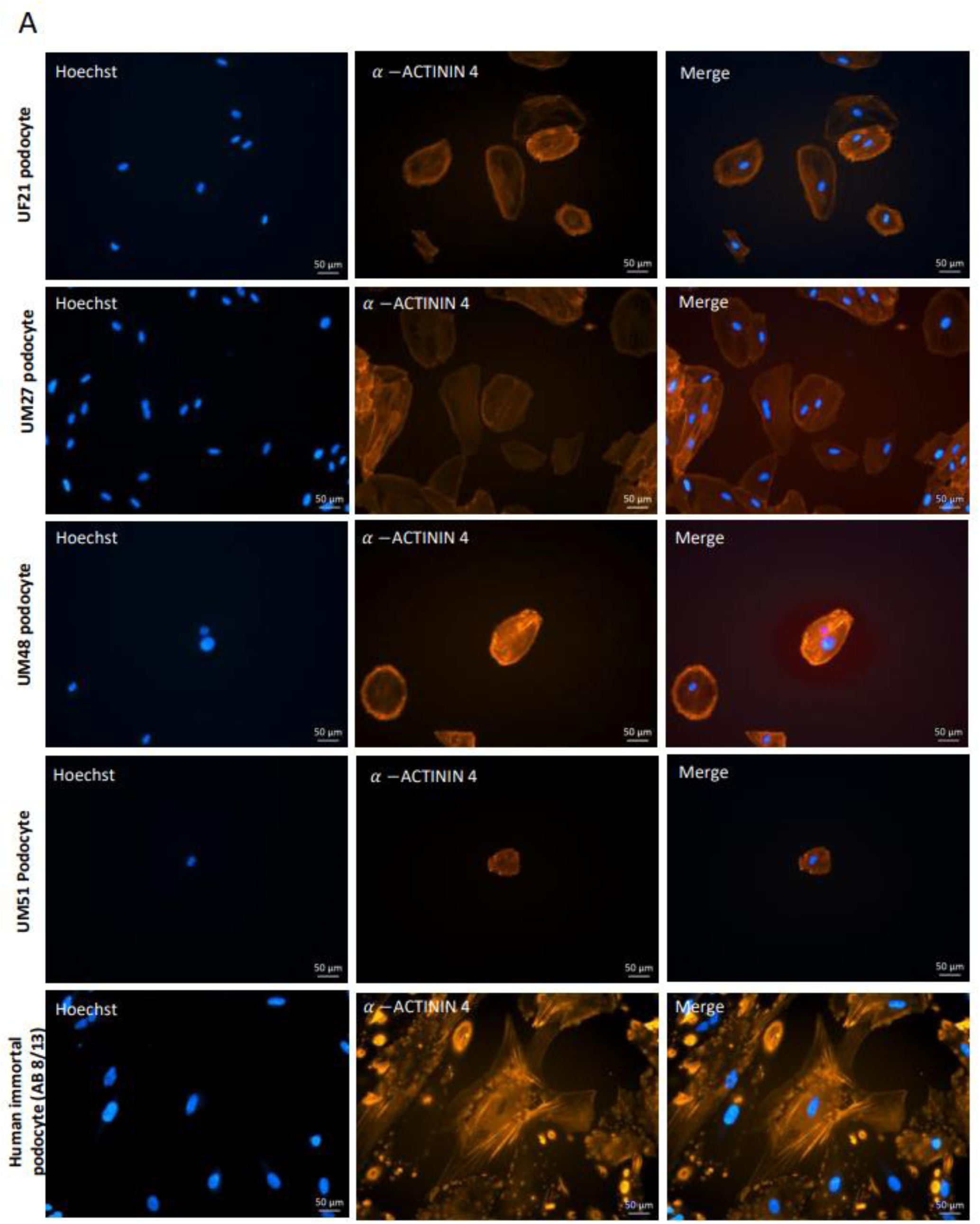
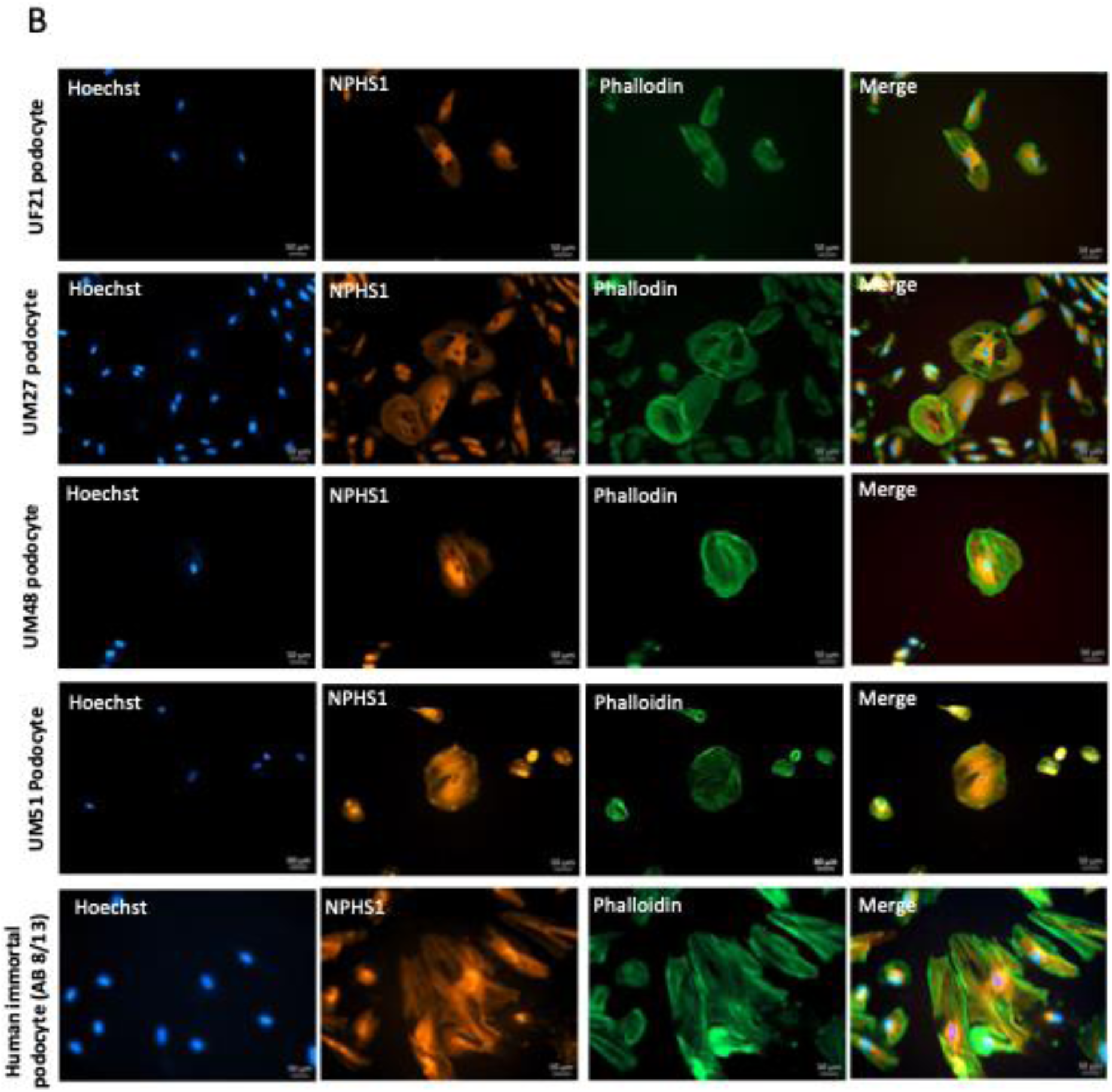
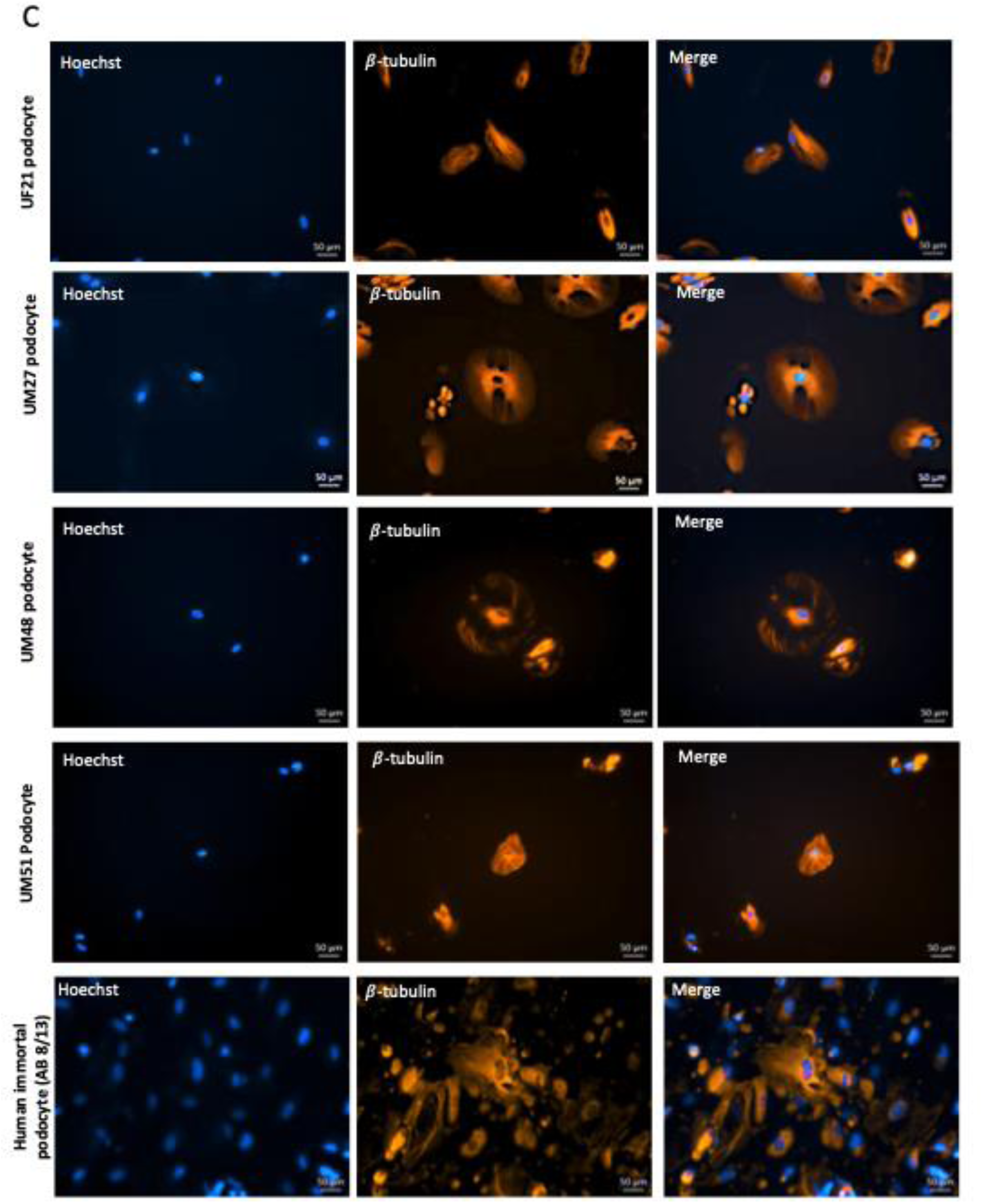
Expression of Alpha-actinin-4 (*ACTN4*), Nephrin (*NPHS1*) and β-TUBULIN (*TUBB*) in human podocytes. Urine-derived nephron progenitor cells (UdRPC) and the immortalized podocyte cell line (AB 8/13) were differentiated into podocytes by culturing at 60–70% density in Advanced RPMI supplemented with 30 μM retinoic acid (RA). The typical Podocyte cytoskeleton was visualized by immunofluorescence-based detection of (**A**) Alpha-actinin-4 (ACTN4), and (**B**) Nephrin (NPHS1) combined with the cytoskeleton marker Phalloidin. As a common brain-associated protein (**C**) β-TUBULIN (TUBB) was stained in all five podocyte preparations.

**Figure 6:**
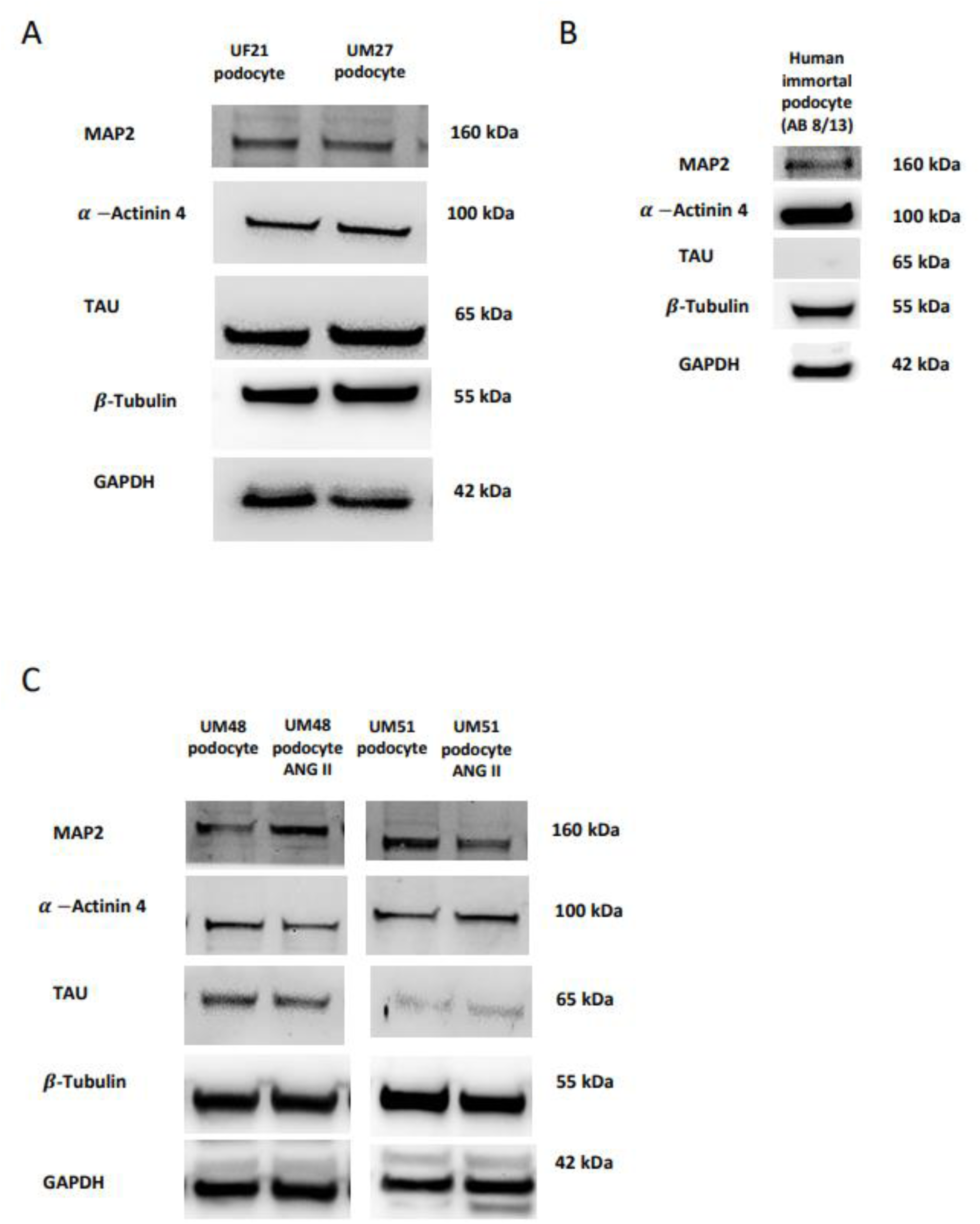
Western Blot analyses of brain-associated proteins expressed in human podocytes. Expression of MAP2, TUBB and TAU were analyzed by Western blotting and the expression of the podocyte markers-ACTN4 was confirmed in all cell cultures. For normalization, expression glyceraldehyde-3-phosphate dehydrogenase (GAPDH) was used. Full uncropped Western blot images are presented in Supplementary Figure S8.

### Responsiveness of genes expressed in podocytes and brain to the renin-angiotensin-system mediator-ANG II

Figure 7a shows a cluster analysis and heatmap of the 344 genes overlapping between UdPodocytes and kidney and brain organoids in UdPodocytes untreated (ctrl, red color bar), treated 24h with ANGII (blue color bar) and other treatments including the Angiotensin-II-type I receptor (AT1R) antagonist Losartan (no color bar). Following our previous publications showing down-regulation of key molecules of the actin cytoskeleton by ANG II [13] and beneficial effects of the ATR1 blocker - Losartan [30] we were interested in down-regulated gene clusters upon 24h of ANGII treatment. We then used these genes for a follow-up cluster analysis and heatmap (Figure 7b) in UdPodocytes from two individuals untreated (ctrl, red color bar), treated 6h with ANGII (effect not yet apparent, also red color bar) and treated 24h with ANGII (blue color bar). Furthermore, genes from the cluster down-regulated by ANG II were subjected to GO analysis (Figure 7c, Suppl. Table S4) and revealed terms related to *projection morphogenesis, axon development* and *AMPA receptors* among the most significantly over-represented Biological Processes. Over-representation analysis results of KEGG pathways (Figure 7d) included significant (p < 0.05) *axon guidance, Hippo signaling, Calcium signaling* and *cGMP-PKG signaling*.

**Figure 7:**
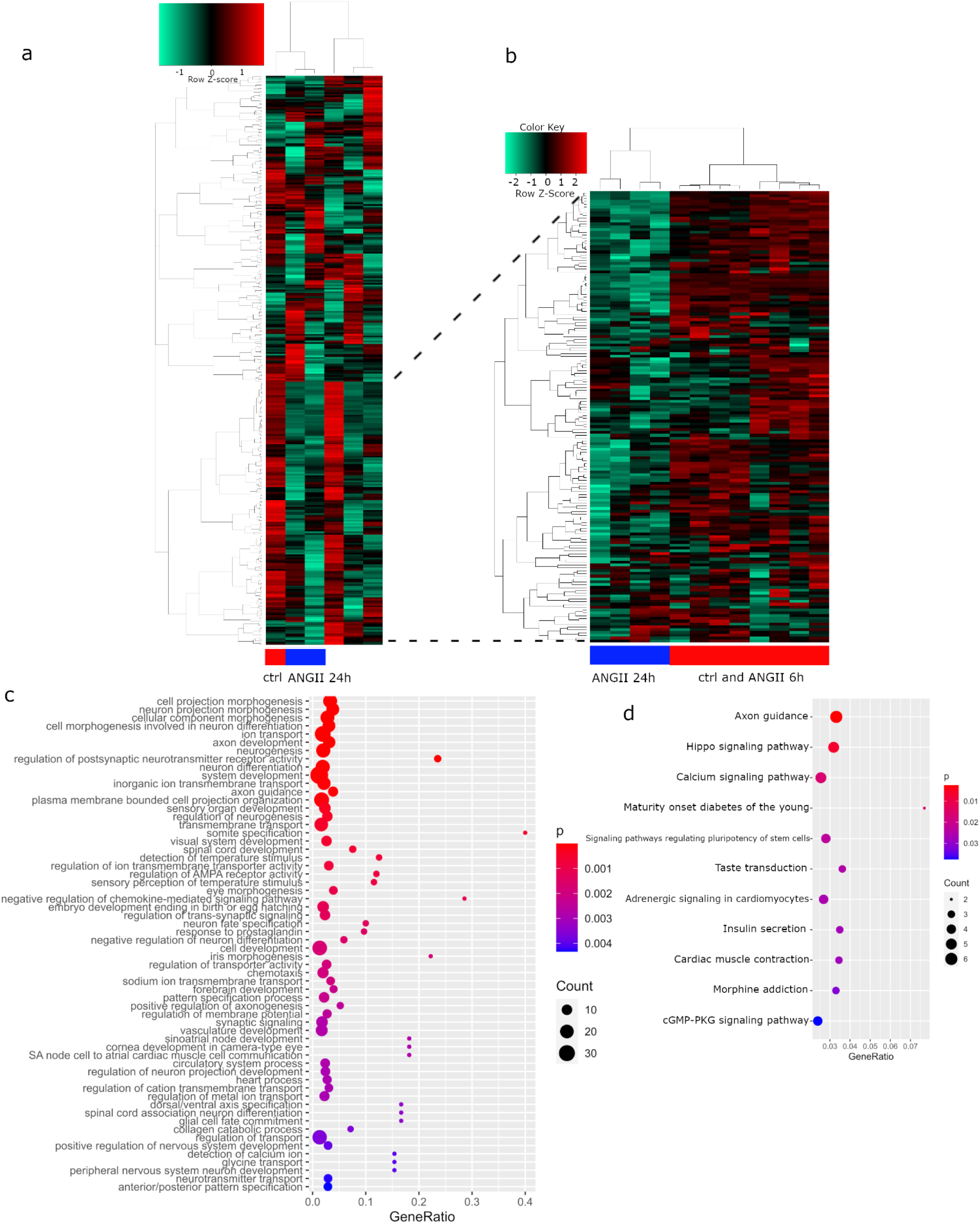
The 344 genes overlapping between UdPodocytes and kidney and brain organoids are responsive to ANGII stimulation. (a) cluster analysis and heatmap of the 344 genes overlapping between UdPodocytes and kidney and brain organoids in UdPodocytes untreated (ctrl, red color bar), treated 24h with ANGII (blue color bar) and other treatments including the AT1R antagonist Losartan (no color bar). This cluster analysis dissected one cluster which was down-regulated upon 24h of ANGII treatment and on which we focused in the follow-up analysis. (b) cluster analysis and heatmap in UdPodocytes from two individuals using the genes from the ANGII-down-regulated cluster in UdPodocytes untreated (ctrl, red color bar), treated 6h with ANGII (effect not yet apparent, also red color bar) and treated 24h with ANGII (blue color bar). (c) The 60 most significantly over-represented gene ontologies of type Biological Process include terms related to *projection morphogenesis, axon development* and *AMPA receptors*. (d) KEGG pathways significantly over-represented (p < 0.05) include *axon guidance, Hippo signaling, Calcium signaling and cGMP-PKG signaling*.

### Comparison of synaptic pathways between brain and podocytes

Figure 8 is an illustration of the 344 genes expressed in common between UdPodocytes, kidney and brain organoids which were found enriched (p=3.89 10^-4^, adjusted_p=0.037) in the KEGG pathway hsa04721 *Synaptic vesicle cycle* via EnrichR enrichment analysis. The involved genes were mapped to the pathway chart and marked with red shading. The pathway chart in Figure 8 shows that most genes of the synaptic vesicle cycle are also expressed in podocytes therefore implying overlapping mechanisms in neurotransmitter uptake, Calcium signalling and vesicle processing.

**Figure 8:**
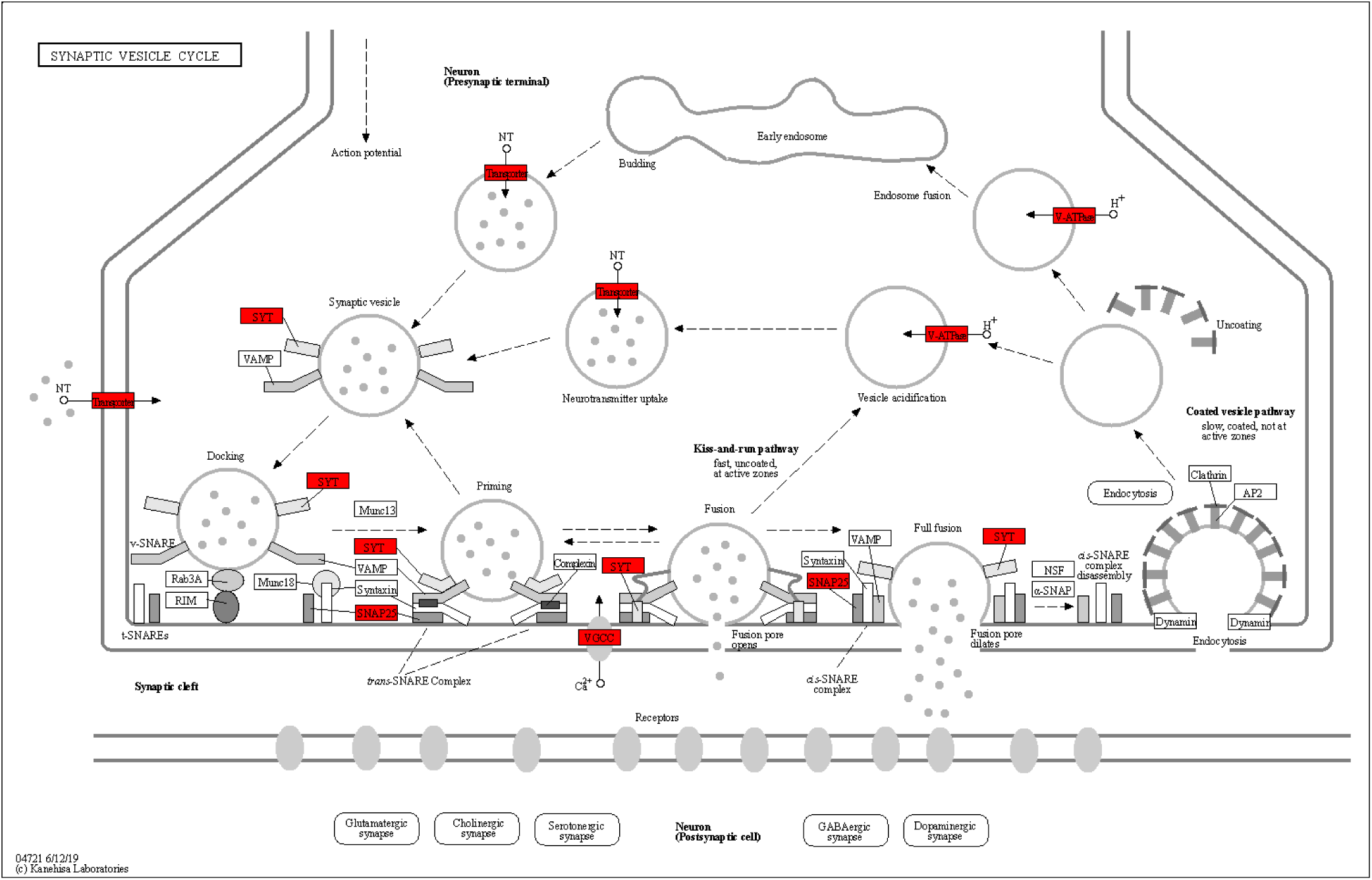
Most genes of the synaptic vesicle cycle are expressed in podocytes and in brain. The 344 genes expressed in UdPodocytes and in kidney as well as in brain organoids were subjected to EnrichR analysis and the KEGG pathway hsa04721 *Synaptic vesicle cycle* was found as one of the most enriched pathways (p=3.89 10^-4^, adjusted_p=0.037). Genes in crucial locations of the pathway are involved as illustrated by the boxes with red shading.

In Figure 9a, genes from the KEGG pathway hsa04728 *Dopaminergic synapse* expressed in common in UdPodocytes, kidney and brain organoids are shown. The EnrichR enrichment analysis identified this pathway as the most significant specific synapse-related pathway (p=0.027, adjusted_p=0.485). The pathway map shows additionally numerous genes of post-synaptic cells are expressed in podocytes and kidney organoids. CACNA1C (indicated in the map as Cav 2.1/2.2) involved in the calcium channels has been reported to be a major player in polycystic kidney disease [34]. In general, ion channels mediate the homeostasis of electrolytes and mutations in the genes coding them may lead to renal disease [35], [12]. Furthermore, the KEGG pathway-hsa04724 *Glutamatergic synapse* (p=0.047, adjusted_p=0.485) was found over-represented (Figure 9b) and thus may imply similar mechanisms in podocytes and diverse types of synapses.

**Figure 9:**
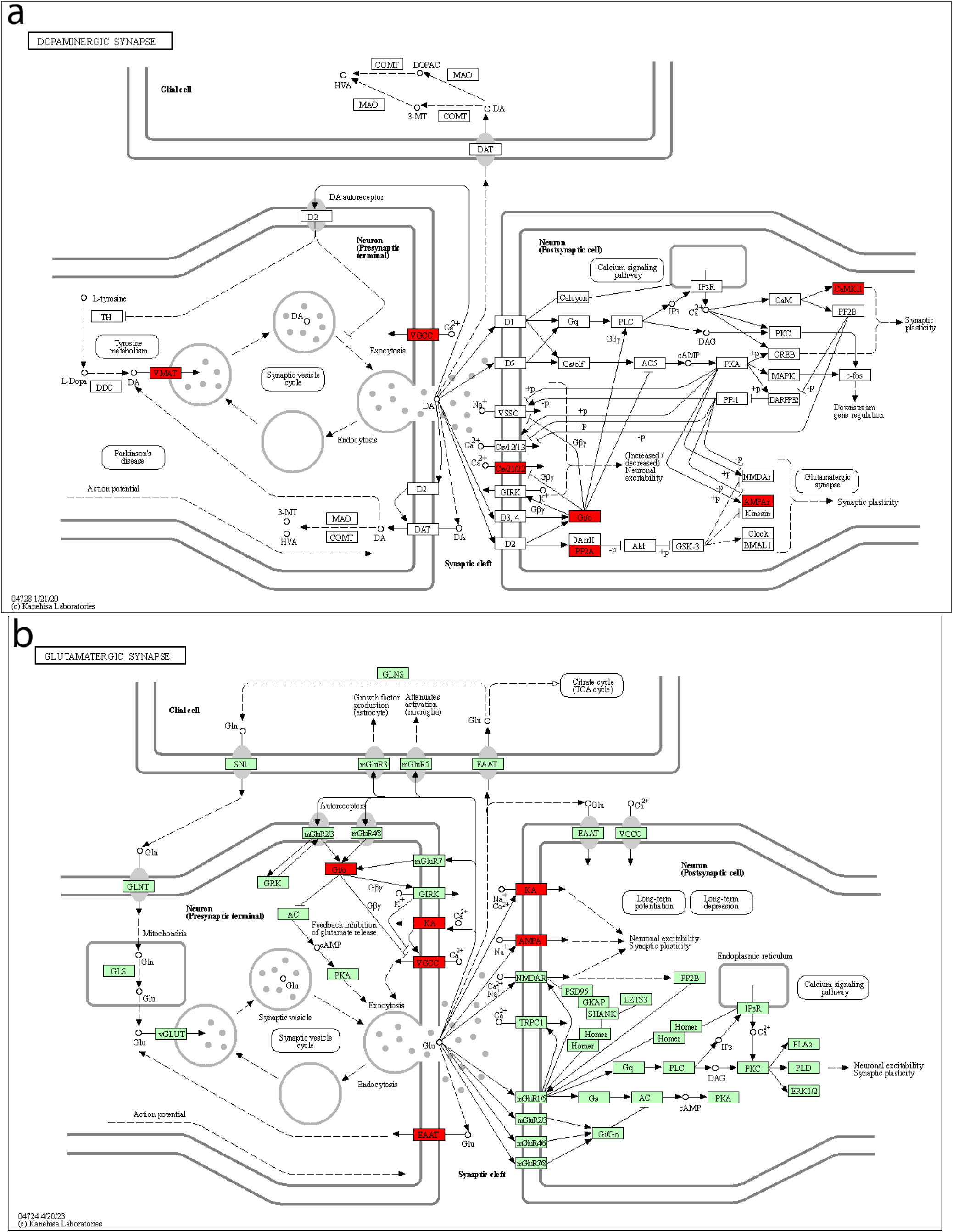
Genes of the pre- and post-synaptic parts of the Dopaminergic and Glutamatergic synapse are expressed in podocytes and brain. The 344 genes expressed in UdPodocytes and in kidney as well as in brain organoids were subjected to EnrichR analysis and the KEGG pathways (a) hsa04728 *Dopaminergic synapse* (p=0.027, adjusted_p=0.485) and (b) hsa04724 *Glutamatergic synapse* (p=0.047, adjusted_p=0.485) were found amongst the most enriched pathways. Genes in crucial locations of the pathways, e.g. Dopamine uptake into vesicles, synaptic plasticity and calcium signaling, are involved in the Dopaminergic synapse (a) and glutamate receptors of AMPA and kainate types triggering neuronal excitability and synaptic plasticity in the Glutamatergic synapse (b), as illustrated by the boxes with red shading.

## Discussion

In this study we compared gene expression in podocytes and brain and unveiled considerable overlap. Starting with genes expressed exclusively in UdPodocytes [13], [30] in comparison to SIX2-posive UdRPCs [19] we identified a plethora of genes over-represented in brain-associated GOs such as *synapses, synaptic signaling, neuron projections/axons* and *myelination*. Our in vitro model revealed expression of Nephrin (NPHS1) and axon-like projections from cells with synapses connecting to other cells upon differentiation of UdRPCs into UdPodocytes (Figure 1). The major function of NPHS1 as a cell adhesion molecule that functions in the glomerular filtration barrier in the kidney has been widely described [36]. Moreover, expression of NPHS1 has been identified in brain [31] where Putaala et al. conclude that it has a developmental role and is mainly expressed in radial glia cells [31]. Li et al. report Nephrin expression close to synaptic proteins in primary neuronal cells and co-immunoprecipitation with synaptic molecules including glutamate receptors and PSD95 and hypothesize that podocytes and brain cells use similar organization and signaling mechanisms [37]. One crucial player in the organization of foot processes and synapses is ACTN4, which bundles F-actin in the cytoskeleton and cross-links it to developing processes, in neurons and podocytes acting together in a network with glutamate receptors [38], [37]. These control the plasticity of dendritic spines in neurons in response to signaling events [38]. A corresponding mechanism of podocyte glutamatergic signaling participating in the integrity of the glomerular filtration barrier was reported by Giardino et al. who identified foot process effacement in a knock-out mouse model of Rab3a which regulates exocytosis of glutamate [5]. In our human UdPodocyte model, we found ACTN4 expression connecting the cytoskeleton of podocytes.

Another Actin-associated protein SYNPO, which is evident in the present transcriptome data, regulates the cytoskeleton in mature podocytes, where it is essential for the functioning of podocyte foot processes. Previous studies have shown that SYNPO also plays a role in the formation of spine apparatuses in spines of telencephalic neurons, which is involved in synaptic plasticity [8], [9]. Dendritic spines are small thorn-like protrusions found on the dendrites of most of the excitatory neurons in the brain. They are the point of contact between neurons, forming the post-synaptic side of the excitatory synapses, and orchestrate the molecular machinery that allows the transmission of signals from the afferent neurons [39], [40], [41], [42]. The stability of the spines is associated with SYNPO [43], [44], [45], [46]. Studies have shown that deletion of SYNPO in mice results in deficits in spatial learning [47], [48]. Furthermore, podocytes lacking SYNPO acquire cytoskeletal alterations, which results in damage to the podocytes and disturbance of the foot processes and the slit diaphragm [49], [8].

ACTN4, microtubule-associated proteins such as TUBB1 and MAPT (TAU) are cytoskeletal-associated proteins expressed in podocytes and down-regulated upon ANGII stimulation [13] TAU links actin and microtubule networks [50], [51].

With respect to these partially similar signaling mechanisms in podocytes processes and neuronal synapses, Weide and Huber ask what is pre- and post-synaptic [52] refering to the publication of Giardino et al. who report Myosin IIA at post-synapses and in foot processes [5]. Moreover, Mundel and colleagues report Synaptopodin (SYNPO) at podocyte foot processes and dendritic spines [9] and Kobayashi and colleagues [6] find the typical postsynaptic density protein Densin localised in the cytoplasm beneath the slit membrane.

UdPodocytes constitute a promising model for studying cytoskeletal dynamics in healthy and diseased podocyte.

Genes such as *SERPINI1*, *RSPH1*, *KCNQ3* and *PAX*6 are also expressed in both brain and podocytes (Table S2). PAX6 is a crucial transcription factor associated with brain development and function. It has been shown to regulate genes associated with neural stem and progenitor cells. Studies have shown molecular interactions, such as the regulation of cell-cell adhesion or ion transport [53], [54]. PAX2 and PAX8 are crucial regulators of multiple steps of renal system development [55]. However, to date studies associating PAX6 function in nephrogenesis is lacking. SERPINI1 is a Serine protease inhibitor that inhibits plasminogen activators and plasmin but not thrombin [56], [57], [58], [59], [60]. It is involved in the formation or re-organization of synaptic connections as well as for synaptic plasticity in the adult nervous system and protects neurons from cell damage by tissue-type plasminogen activator and probably plays the same role in kidney.

The gene *KCNQ3,* encodes a potassium channel protein that functions in the regulation of neuronal excitability and is considered to be a key characteristic of activity-dependent synaptic plasticity of neurons [61]. One of the fundamental functions of renal K channels is their role in generating the cell-negative potential that provides one of the main driving forces for the passive movement of charged solutes and carrier molecules across the apical and basolateral membranes of tubule cells [62].

In-depth analyses of the 600 brain-annotated genes expressed in UdPodocytes revealed GOs such as *excitatory synapse* and *AMPA glutamate receptor complex*. Amongst this list, GRIA3 has been reported to be epigenetically down-regulated in podocytes under diabetic conditions [63]. Additionally, we found AMPA receptor over-represented among genes expressed in brain and podocytes and down-regulated upon ANGII stimulation of Upodocytes. AMPA belongs to the group of ionotropic glutamate receptors [64] and the same holds for NMDA receptors which are more intensively studied with regards to their role in regulating ion channels in podocytes [65], [66]. In comparison to neurons, NMDA receptors have overlapping and distinct features in podocytes, e.g. unlike in neurons they are resistant to activation by L-glutamate and L-aspartate and show minimal inactivation upon continuous NMDA supply, but similar to neurons they can be activated by D-aspartate and L-homocysteic acid [65]. Staruschenko et al. propose that NMDA receptors in podocytes are more adapted to slowly flowing metabolites in contrast to fast synaptic signaling processes. Our finding of AMPA receptor expression in podocytes together with Li et al’s report of down-regulated AMPA-associated GRIA3 in podocytes under diabetic conditions [63] suggests that further studies investigating the role of AMPA receptors in podocyte physiology will be promising.

A body of publications describe ANGII signaling in the brain and beyond that a brain renin-angiotensin system which despite profound evidence [67], [68], [69], [70] is also a subject of debate [70]. Persistent activation of ANGII signaling can lead to neurodegeneration and Angiotensin receptor blockers (ARBs) possess therapeutic potential [70]. Besides its role in regulating blood pressure, ANGII has also been reported to act as a neurotransmitter [71] which can interact with glutamate, possibly acting on glutamate receptors and also mediating increase of blood pressure, upon glutamate administration into the paraventricular nucleus (PVN).

We identified Calcium signaling as associated with the ANGII -down-regulated geneset. Greka and Mundel demonstrated that ANGII triggers - Calcium (Ca2+) flow into podocytes via transient receptor potential canonical 6 (TRPC6) and TRPC5 channels and modulate actin cytoskeleton [72], by interacting with Ca2+-activated phosphatase, Calcineurin, and Rho GTPases which are crucial determinants of podocyte function. Impairments in these channels e.g. by mutations in TRPC6 can lead to Focal Segmental Glomerusclerosis (FSGS). Besides Calcium signaling, Hippo Signaling, cGMP-PKG signaling and axon guidance pathways were also over-represented. cGMP-bound ion channels regulate sodium uptake and excretion [73]. Chen et al. [74] described YAP and TAZ, as key players in Hippo signaling which are important for the integrity of podocytes and TAZ together with the transcription factor TEAD activate crucial podocyte-associated genes such as *SYNPO*, *ZO-1* and *ZO-2* (synaptopodin, zonula occludens-1, and zonula occludens-2). Interestingly, proteins from the axon guidance pathway were found to be associated with risk of end-stage kidney disease in diabetes [75]. The genes in our ANGII-down-regulated signature are all upstream of the actin-cytoskeleton-regulation complex within the axon guidance pathway thus pointing at down-regulation of actin cytoskeleton by ANGII treatment, in line with our previous publication showing ANGII-dependent downregulation of *NPHS1* and *SYNPO* and subsequent disruption of the podocyte cytoskeletal architecture [13]. Moreover, Shao et al. [76] revealed that HNF1B regulates axon guidance-associated genes in the developing mouse kidney and suggest that axon guidance ligand receptor pairs of neighboring cells, some also from the ureteric bud, may orchestrate kidney development via auto- and paracrine signaling. This may be underlined by Combes et al. 2019 [77] who found axon-guidance associated GDNF-RET ligand receptor crosstalk in single-cell analysis of the developing mouse kidney. Analysing the 600 UdPodocyte-specific genes in the axon guidance pathway revealed numerous ligand receptor pairs which regulate both actin cytoskeleton and are responsible for attraction and repulsion of projections. Tapia et al. [78] described the axon-guidance-associated protein-Semaphorin 3A (SEMA3A) as a repulsive gene leading to foot process effacement and proteinuria upon exogenous induction of SEMA3A. Additionally, Lu and Zhu [79] describe involvement of several Semaphorins in diabetic nephropathy and partially aberrant foot processes in Sema3G-deficient mice. Hashimoto et al. [80] described co-localization of the axon guidance receptor Ephrin-B1 with Nephrin at the slit diaphragm between neighboring foot processes during development and is down-regulated in nephropathy. Furthermore, Wu et al. [81] describe a major role of the axon guidance factors Slit2 and Robo2 in podocyte attachment to the glomerular basement membrane and the structural integrity of foot processes. Taken together, it is tempting to speculate that foot processes partly use similar signaling mechanisms as axon guidance for their interdigitation and impairment of these signaling processes and lead to pathologically altered glomerular permeability.

In the follow-up analysis we identified abundant overlap of 344 genes expressed in common in UdPodocytes and in iPSC-derived brain and kidney organoids. The 344 genes expressed in common map to multiple brain regions with most hits in frontal cortex. As transcription factors enriched in the promoter regions of the 344 genes we found SUZ12 most significant (p=8.42×10-22, EnrichR ENCODE and CHEA datasets), NFKB1 (p=2.29×10-3, EnrichR TRANSFAC and JASPAR) and PAX2 (p=1.59×10-3, EnrichR TTRUST). Mapping genes to the GO Molecular Function top category showed that about 70% of the 344 genes are associated with binding activity while about 25% are associated with catalytic activity.

Of note, we identified Brain-derived neurotrophic factor (BDNF) as central protein in one module associated with “pre-synapse” within our protein interaction network of brain-associated UdPodocyte genes (Figure 4a,c). Besides the brain-associated roles of BDNF in synaptic plasticity, learning, memory, behavior, depression and neurodegenerative disorders such as Alzheimer’s disease [82], Endlich and colleagues proposed BDNF as marker for CKD. They found a strong correlation of BDNF expression with the podocyte marker Nephrin, the kidney injury marker KIM-1 and the urinary albumin-to-creatinine ratio in CKD patients [83]. Putatively connected to BDNF’s role in podocyte development, the authors showed de-differentiation of podocytes upon BDNF inhibition. Furthermore, Saito and colleagues reported another protein from this module, Neurexin-1 (NRXN1) known as presynaptic adhesion molecule associated with synaptic differentiation, expressed at the podocyte slit diaphragm, co-localized with Nephrin and CD2AP, and proposed it as marker of podocyte injury [84]. SNAP25, a further protein of this module, as well as BDNF were shown to be correlated with the Parkinson’s Disease (PD) -associated protein Parkin (PARK2), putatively via ubiquination of p21 and JNK, and inhibition of PARK2 led to down-regulation of BDNF and SNAP25 and impaired neural stem cell differentiation [85]. A study revealing an association between PD and impaired glomerular filtration [86] may be seen in this context.

We conclude that our study has revealed similarities between the transcriptomes of brain and podocytes which can be condensed into gene networks regulating (i) projections such as foot processes, axons and denritic spines implicating modulation of actin cytoskeleton and (ii) signaling pathways such as Calcium-, Hippo- and cGMP-signaling happening at synaptic (e.g. glutamatergic, dopaminergic) or resembling constellations such as the slit diaphragm between interdigitating foot processes.

Interestingly, axon guidance was found as most significant pathway in response to ANG II stimulation of podocytes which might point at similar mechanisms guiding axons and foot processes in kidney development and disease. This was in fact underlined by reports on ligand-receptor crosstalk of axon guidance molecules (GDNF-RET) for the developing kidney and association of axon guidance molecules with risk of end-stage kidney disease in diabetes [79], [87].

Interestingly, many of the brain-associated biological processes are currently not annotated for kidney. This points to the comparative lack of advancement in the annotation of kidney associated processes. We therefore propose our urine-derived podocyte platform as a non-invasive and easily accessible in vitro model enabling the investigation of the functions of these brain-associated genes and biological processes in healthy and diseased kidneys.

## Author contributions

WW and CT analysed the data and wrote the manuscript. JA supervised the work, co-wrote the manuscript and gave the final approval. We thank Leon Szepanowski and Abida Islam Pranty for designing the primers used in this study.

## Acknowledgments

James Adjaye acknowledges financial support from the Medical faculty of the Heinrich-Heine University, Duesseldorf. We thank Prof. Moin Saleem for providing the human kidney-biopsy derived immortalized podocyte cell line used in this study

## Data availability

No datasets were generated in the current study. The datasets used for the meta-analysis are available at the National Center for Biotechnology Information (NCBI) Gene expression Omnibus (GEO) accessions referred to in Table 1.

## Supplementary Material

**Supplementary Table 1 (tableS1.xlsx): significantly over-represented GOs in the 600 genes subset only expressed in UdPodocytes from the Venn diagram of UdPodocytes vs. UdRPCs filtered for neuronal terms**

**Supplementary Table 2 (tableS2.xlsx): (a) subsets from the Venn diagram comparing the 600 genes exclusively expressed in UdPodocytes with genes expressed in kidney and brain organoids. (b) Significantly over-represented GOs in the 344 genes subset expressed in common in UdPodocytes and brain and kidney organoids. (c) enriched transcription factors (TFs) found via the EnrichR analysis of the 344 genes using the ENCODE_and_ChEA dataset. (d) enriched TFs found via the EnrichR analysis of the 344 genes using the TRANSFAC_and_JASPAR dataset. (e) enriched TFs found via the EnrichR analysis of the 344 genes using the TRRUST dataset. (f) 20 most abundant GO Molecular Function (MF) terms amongst the 344 genes. (g) Metascape enrichment analysis results of genes in (f). (h) TFs from (f) validated for expression in kidney via the Proteinatlas.**

**Supplementary Table 3 (tableS3.xlsx): (a) subsets from the Venn diagram comparing the 600 genes exclusively expressed in UdPodocytes with genes expressed in brain in the GTEX database. (b) Significantly over-represented GOs in the 130 genes subset expressed in common in UdPodocytes and GTEX brain**

**Supplementary Table 4 (tableS4.xlsx): (a) Significantly over-represented GOs in the genes down-regulated upon ANGII treatment amongst the 344 genes expressed in common in UdPodocytes and brain and kidney organoids. (b) GO terms in (a) filtered by neuronal and brain-related key words listed at the bottom of the sheet.**

**Supplementary Table 5 (tableS5.xlsx): Antibodies used in this study.**

**Supplementary Table 6 (tableS6.docx): Primer sequences used in this study.**

## Supplementary Figures

**Figure S1:**
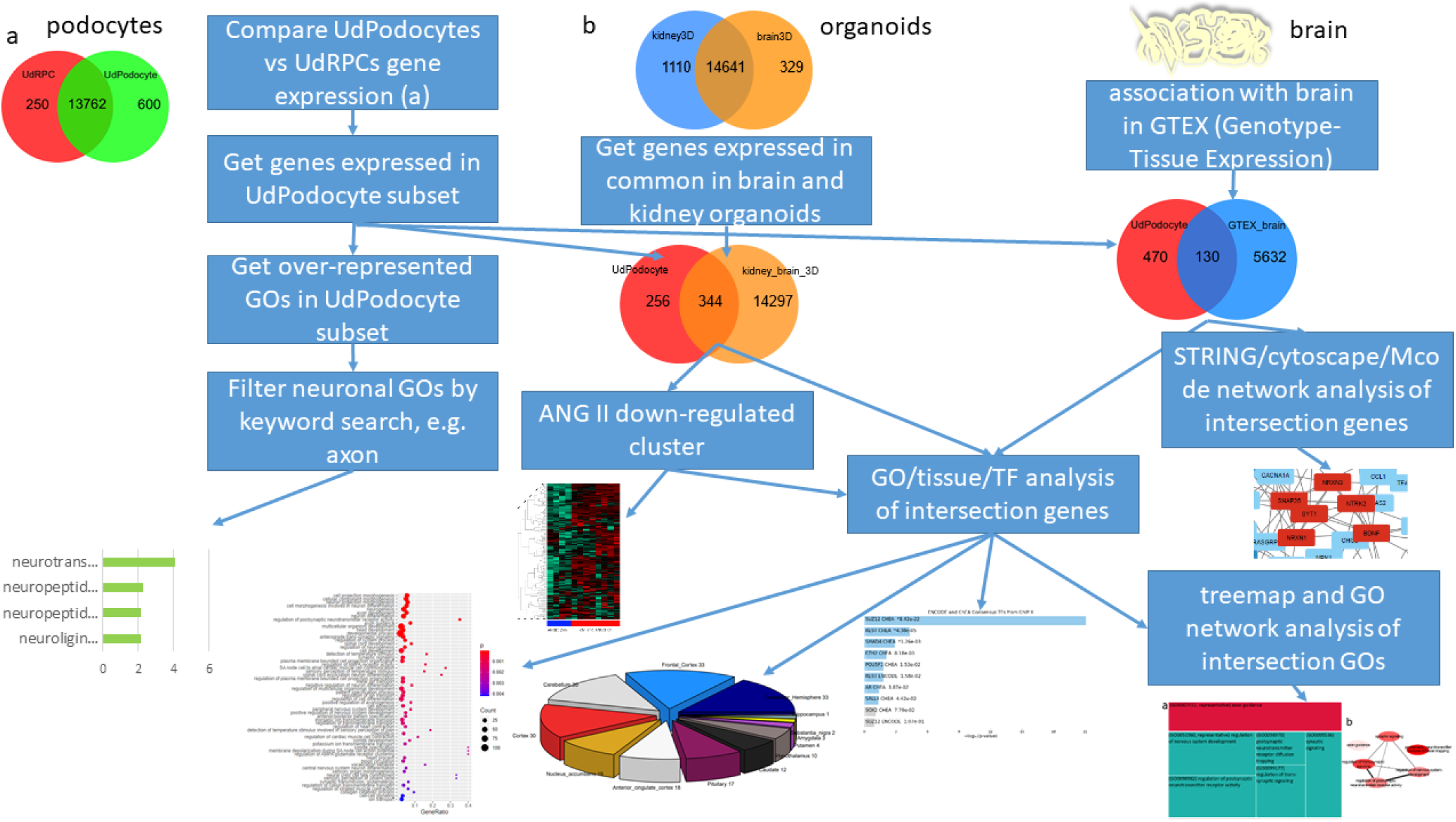
Flow chart of the comparison between brain and podocytes.

**Figure S2:**
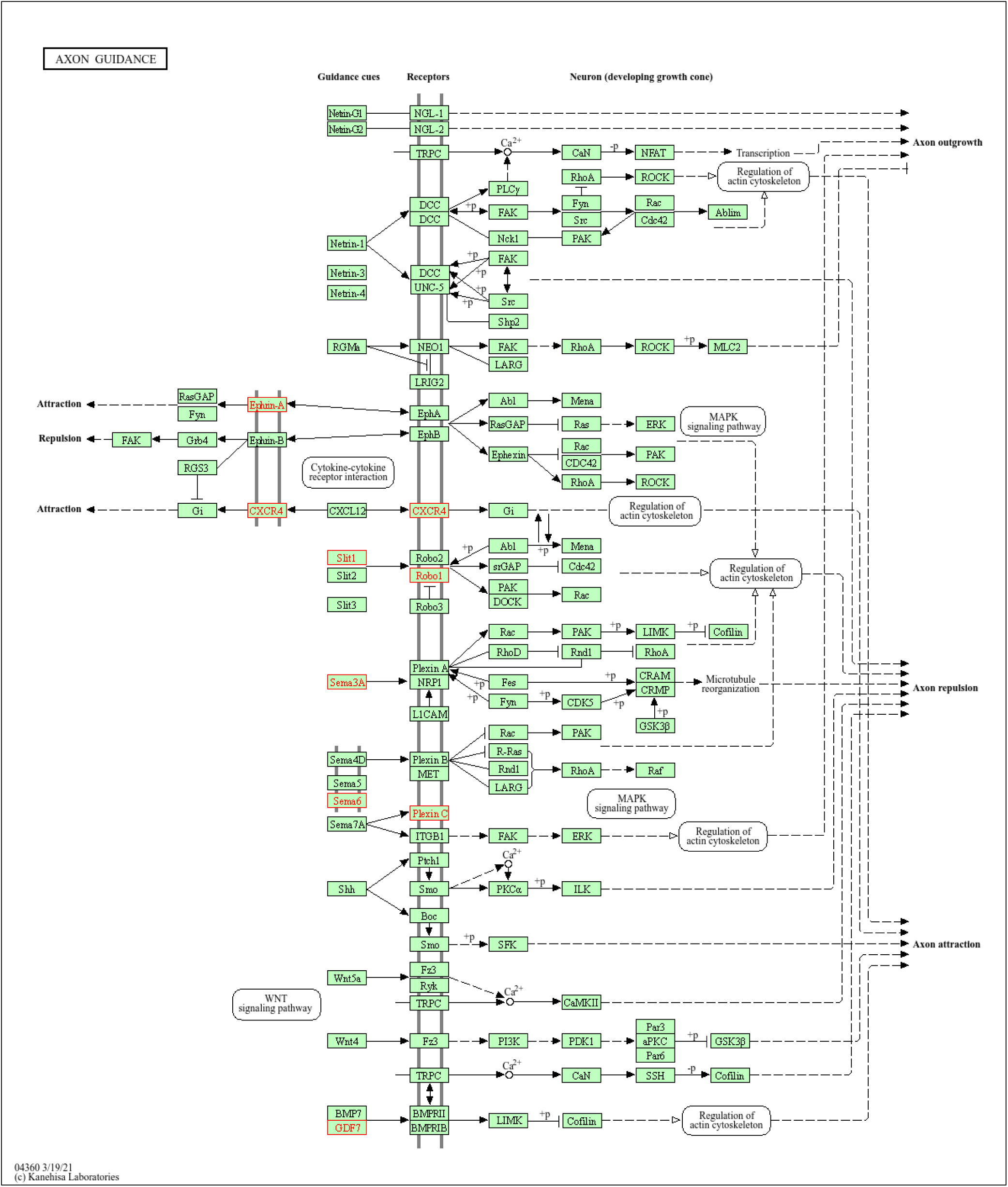
Genes related to the GO term “axon guidance”, the most significantly over-represented biological process in the 600 genes exclusively expressed in urine-derived Podocytes (UdPodocytes), marked within the KEGG pathway chart of axon guidance (hsa04360). Several guidance factors, such as ephrins, Slits, and semaphorins are expressed in the UdPodocytes. These ligands and receptors are responsible for the regulation of the actin cytoskeleton on the one hand and on the other hand for attraction of the projection while genes responsible for repulsion are missing. Genes exclusively expressed in the UdPodocytes are marked in red.

**Figure S3:**
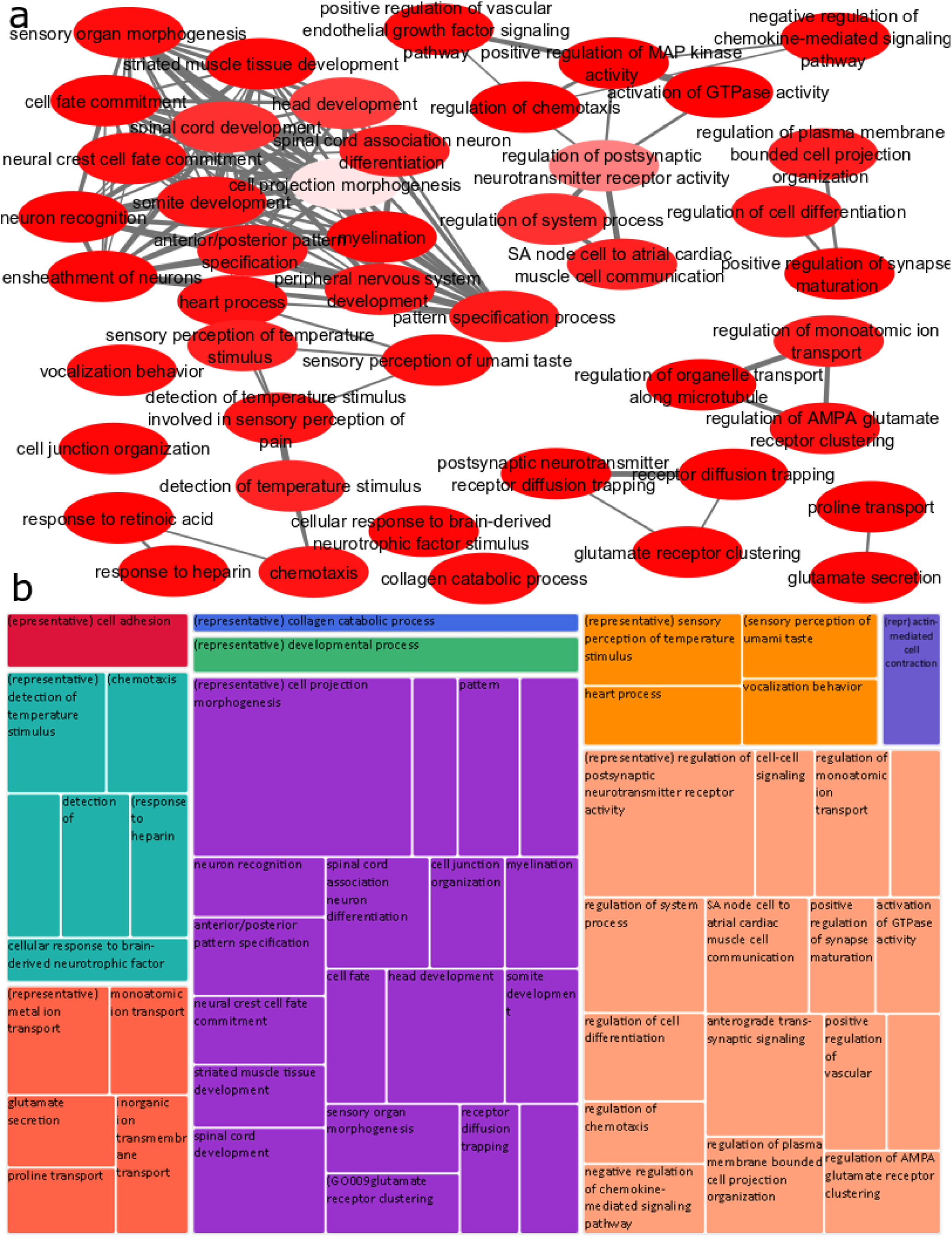
Overlap between biological processes in urine-derived Podocytes (UdPodocytes) with brain and kidney organoids (344 genes) relates to cell projection morphogenesis and regulation of postsynaptic neurotransmitter receptor activity. (a) The gene ontology network was generated with the tools REVIGO and Cytoscape and summarizes GO-BP (Gene ontologies - Biological Process) terms found over-represented with a p-value < 0.01 in the 344 genes overlapping between UdPodocytes and brain and kidney organoids. Cell projection morphogenesis and regulation of postsynaptic neurotransmitter receptor activity -related terms emerged as representative for their clusters. GOs are represented by the network nodes with light red associated with the highest significance of over-representation of a GO term. The edges refer to similarities between the GO terms. (b) Treemap of the REVIGO tool corresponding to (a) summarizing biological process overlapping between UdPodocytes and brain and kidney organoids.

**Figure S4:**
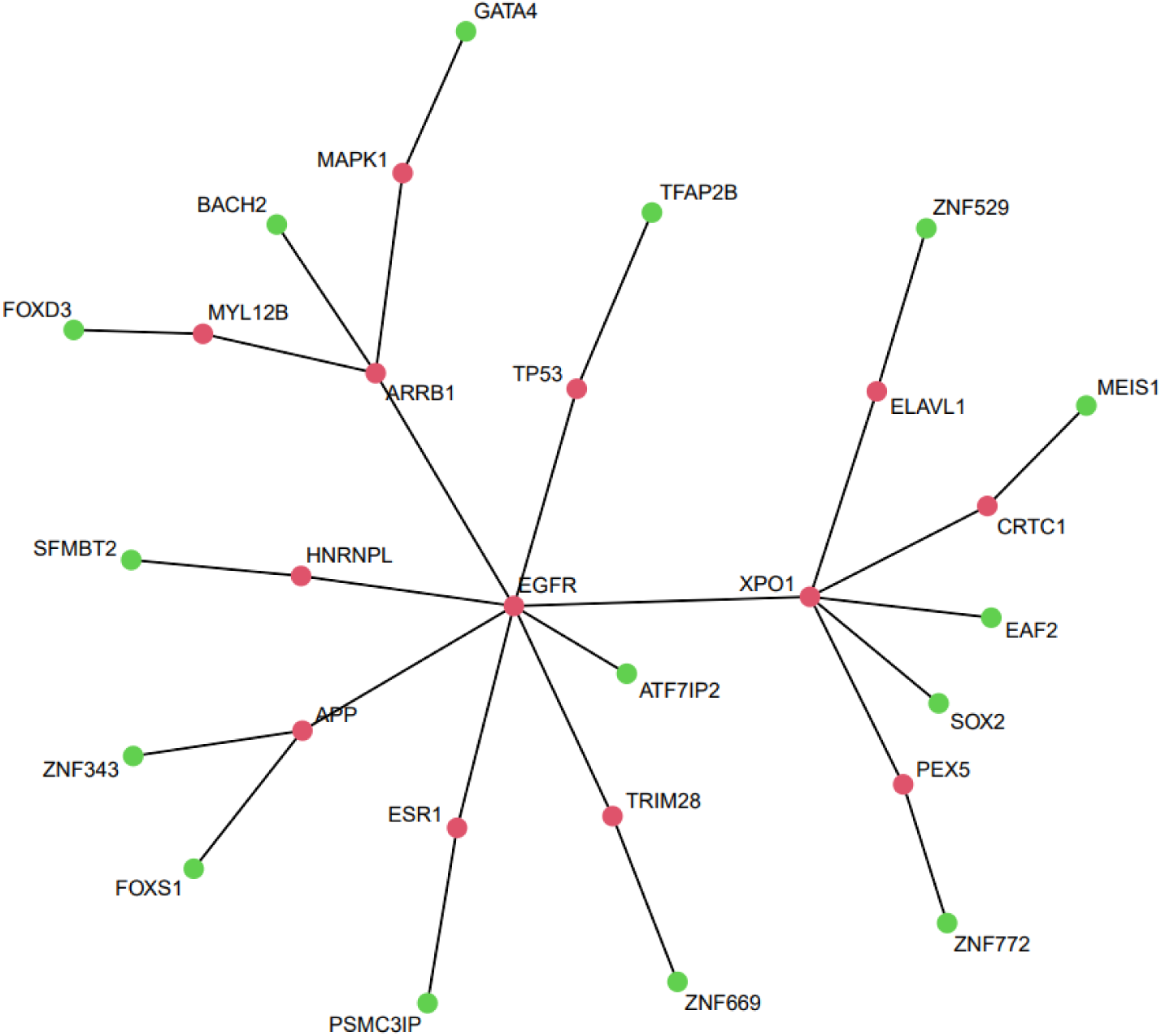
Protein interaction network of Proteinatlas-validated Transcription Factors (TFs) amongst the 344 genes subset expressed in common in UdPodocytes and brain and kidney organoids. The TFs validated for expression in the kidney via the Proteinatlas could be connected in a protein interaction network of Biogrid interactions using EGFR and XPO1 as major hubs. Green nodes are the original TFs and red nodes were added via Biogrid interactions.

**Figure S5:**
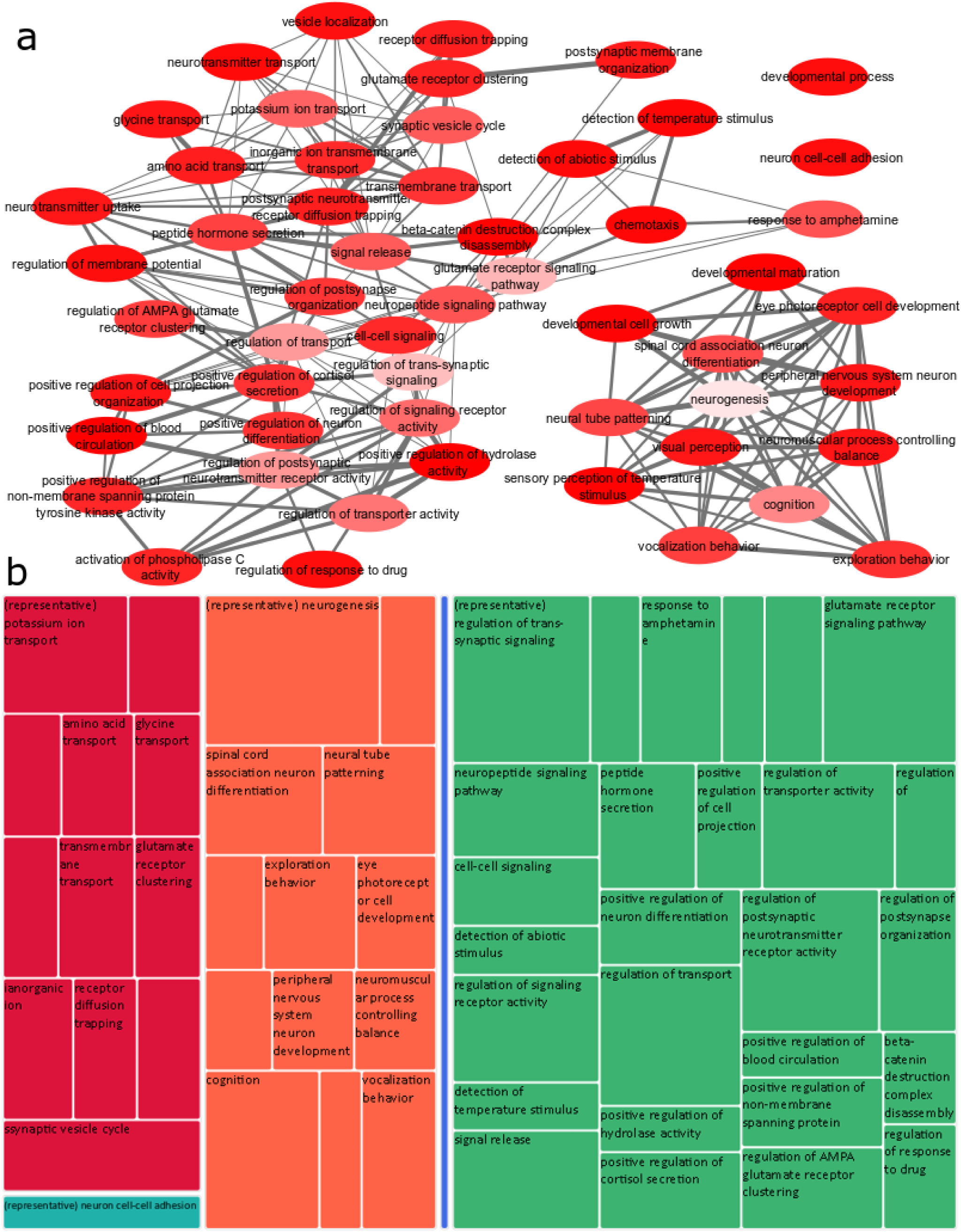
Overlap between biological processes in urine-derived Podocytes (UdPodocytes) with genes associated with brain in Genotype-Tissue Expression GTEX (130 genes) relates to neurogenesis and regulation of trans-synaptic signaling. (a) The gene ontology network was generated with the tools REVIGO and Cytoscape and summarizes GO-BP (Gene ontologies - Biological Process) terms found over-represented with a p-value < 0.01 in the 130 genes overlapping between UdPodocytes and genes associated with brain in GTEX. Neurogenesis and regulation of trans-synaptic signaling -related terms emerged as representative for their clusters. GOs are represented by the network nodes with light red associated with the highest significance of over-representation of a GO term. The edges refer to similarities between the GO terms. (b) Treemap of the REVIGO tool corresponding to (a) summarizing biological process overlapping between UdPodocytes and brain and kidney organoids.

**Figure S6:**
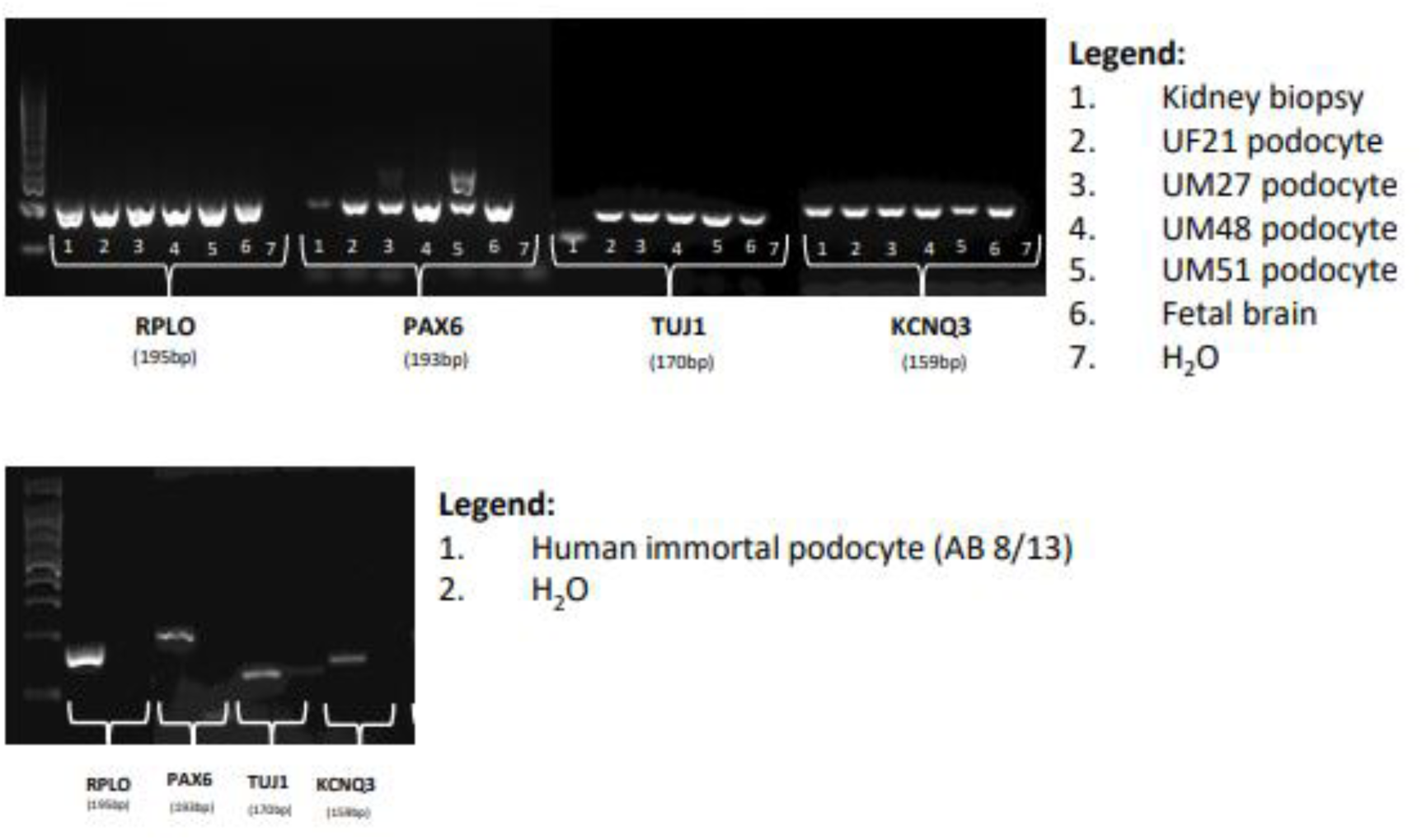
RT-PCR measurements of genes expressed in brain and podocytes. (**A**) RT-PCR measurements of genes *RPLO, PAX6, TUJ1* and *KCNQ3* expressed in brain and podocytes, 1: kidney biopsy, 2: UF21 podocyte, 3: UM27 podocyte, 4: UM48 podocyte, 5: UM51 podocyte, 6: fetal brain, 7: H_2_O. Ribosomal protein lateral stalk subunit P0 (RPL0) was used as a housekeeping gene. (**B**) PCR measurements of the genes *RPLO, PAX6, TUJ1* and *KCNQ3* expressed in the human immortal podocyte line (AB 8/13). The loading scheme was as follows: 1. human immortal podocyte line (AB 8/13) and 2. H_2_O. For normalization, RPL0 was used.

**Figure S7:** Protein expression from the Protein Atlas for the relevant proteins (figS7_Proteinatlas_podobrain.pdf).

**Figure S8:** Complete images of Western Blot analyses of brain-associated proteins expressed in human podocytes (figS8_WB_uncropped.pdf).

