## Supplementary Table 6 for "Evidence for conserved gene expression and biological processes operative in human podocytes and adult brain"

**Supplementary Table 5: Primer sequences used in this study.**

| gene | forward | reverse | bp |
| --- | --- | --- | --- |
| RPL0 | TCGACAATGGCAGCATCTAC | ATCCGTCTCCACAGACAAGG | 195 |
| Pax6 | CAGAGAAGACAGGCCAGCAA | CCATGGTGAAGCTGGGCATA | 193 |
| hTuj1 | ATGAACACCTTCAGCGTCGT | CATCCGTGTTTTCCACCAGC | 160 |
| SYNPO | CCCCAACCTCTCCTCTAACC | ATGACACAGGAGGCAGAAGAAT | 116 |
| KCNQ3 | GACCCCGCAGGGCATC | ACAATCAGGAACACCAACGC | 159 |
| RSPH1 | CAACGGGGACACCTACGAA | TCATTTGCCCACTCTCCTTCAT | 174 |
| Serpini1 | CTGCCACTTATCTGGCCCTC | GGAGCCATCACTAAATTCCCCATA | 176 |
