## Supplementary Figure 7 for "Evidence for conserved gene expression and biological processes operative in human podocytes and adult brain"

### Proteinatlas

Brain related genes

Synapsin1

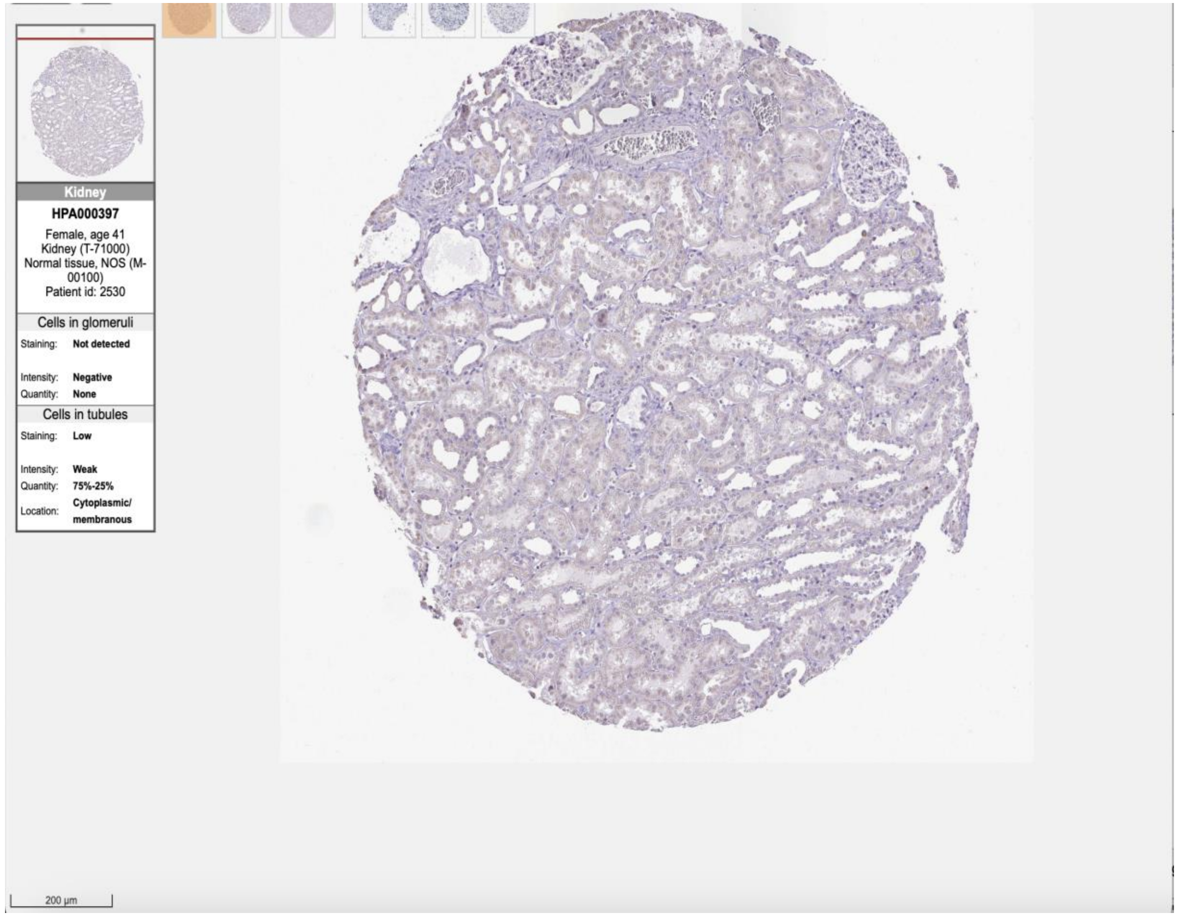
Kidney
HPA000397
Female, age 41
Kidney (T-71000)
Normal tissue, NOS (M-00100)
Patient id: 2530
Cells in glomeruli
Staining: Not detected
Intensity: Negative
Quantity: None
Cells in tubules
Staining: Low
Intensity: Weak
Quantity: 75%-25%
Location: Cytoplasmic/ membranous
200 µm

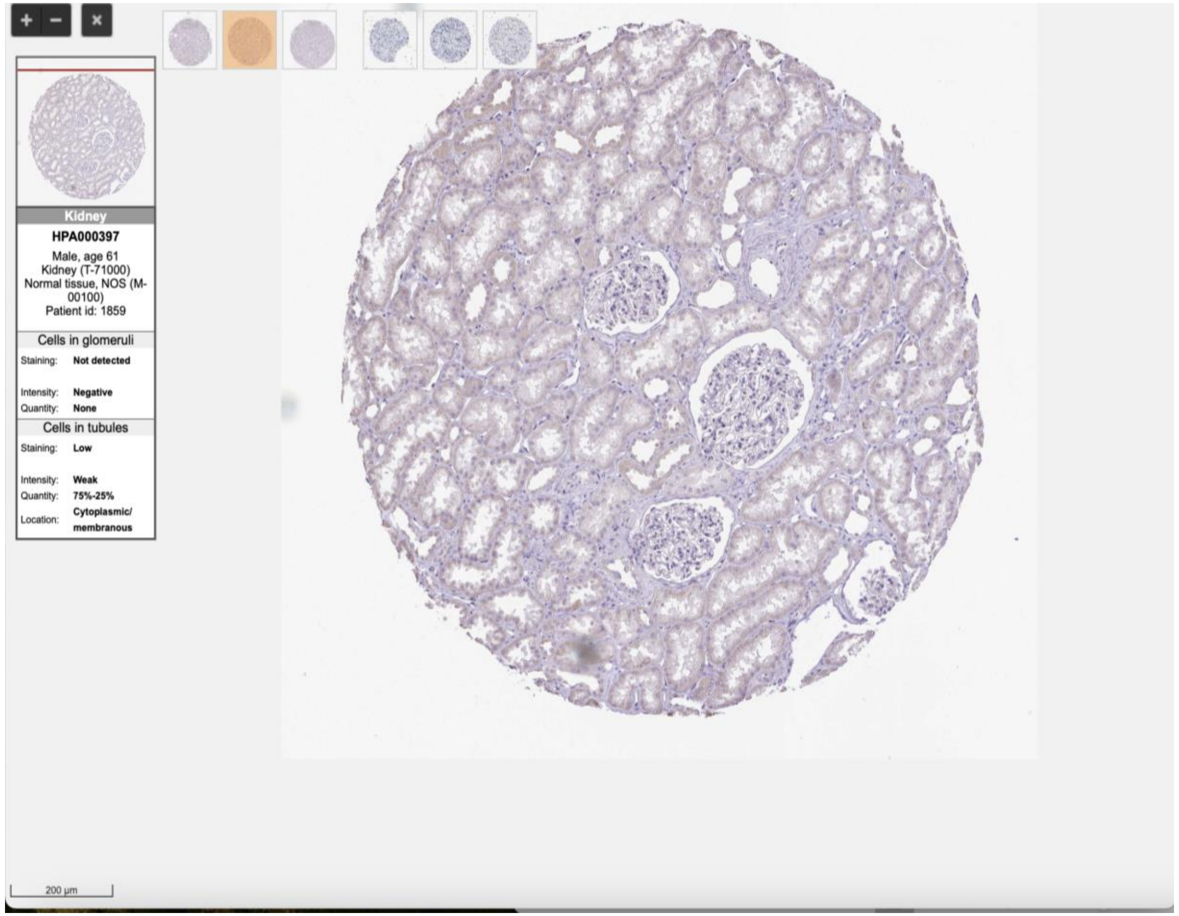
Kidney
HPA000397
Male, age 61
Kidney (T-71000)
Normal tissue, NOS (M-00100)
Patient id: 1859
Cells in glomeruli
Staining: Not detected
Intensity: Negative
Quantity: None
Cells in tubules
Staining: Low
Intensity: Weak
Quantity: 75%-25%
Location: Cytoplasmic/ membranous
200 µm

MAPT

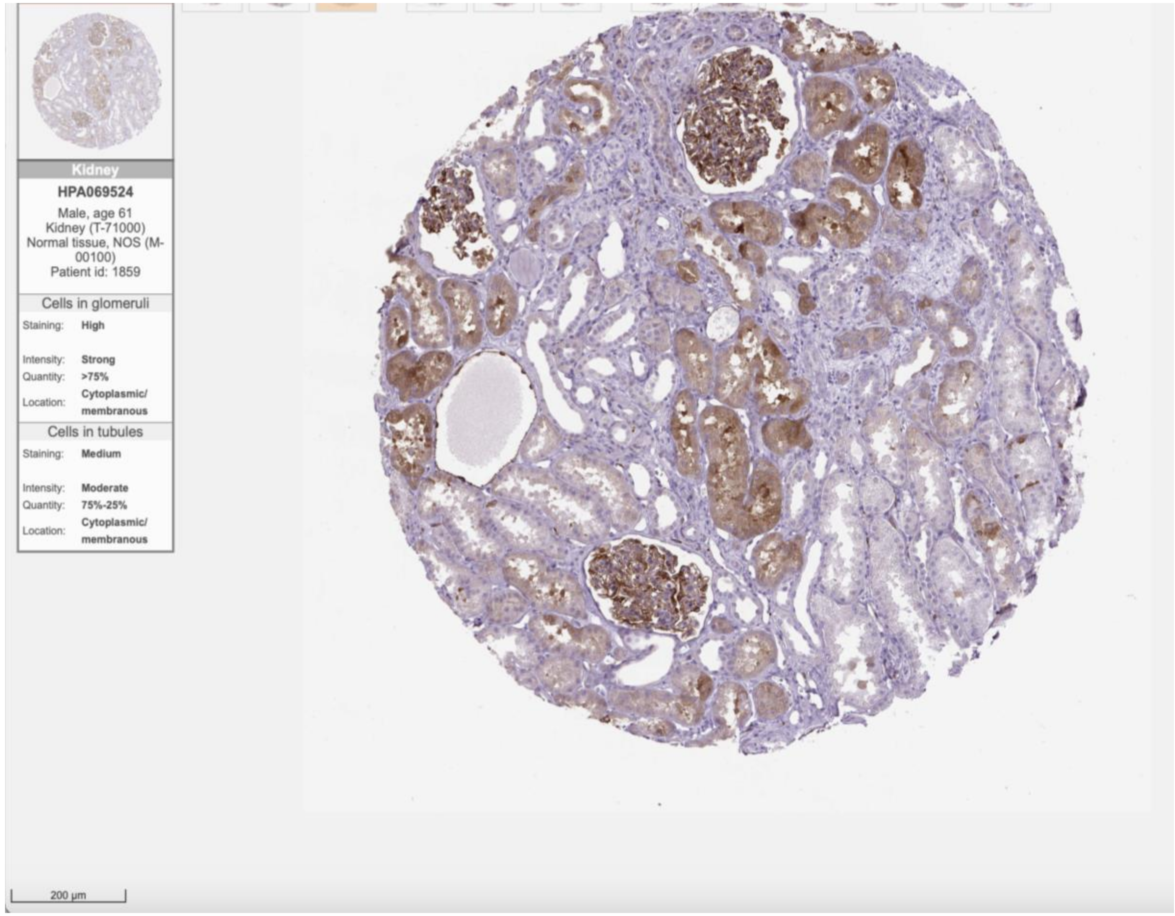
Kidney
HPA069524
Male, age 61
Kidney (T-71000)
Normal tissue, NOS (M-00100)
Patient id: 1859
Cells in glomeruli
Staining: High
Intensity: Strong
Quantity: >75%
Location: Cytoplasmic/ membranous
Cells in tubules
Staining: Medium
Intensity: Moderate
Quantity: 75%-25%
Location: Cytoplasmic/ membranous
200 µm

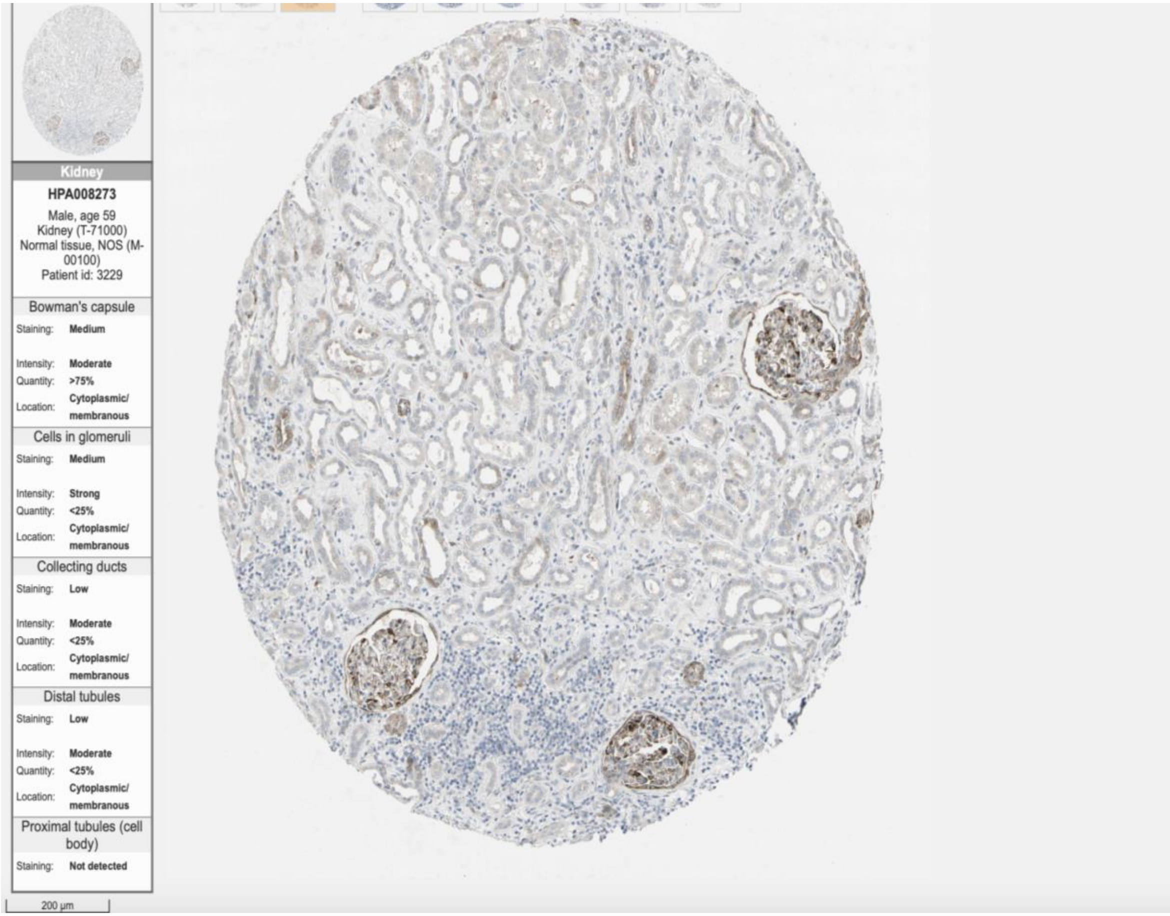
Kidney
HPA008273
Male, age 59
Kidney (T-71000)
Normal tissue, NOS (M-00100)
Patient id: 3229
Bowman's capsule
Staining: Medium
Intensity: Moderate
Quantity: >75%
Location: Cytoplasmic/ membranous
Cells in glomeruli
Staining: Medium
Intensity: Strong
Quantity: <25%
Location: Cytoplasmic/ membranous
Collecting ducts
Staining: Low
Intensity: Moderate
Quantity: <25%
Location: Cytoplasmic/ membranous
Distal tubules
Staining: Low
Intensity: Moderate
Quantity: <25%
Location: Cytoplasmic/ membranous
Proximal tubules (cell body)
Staining: Not detected
200 µm

MAP2

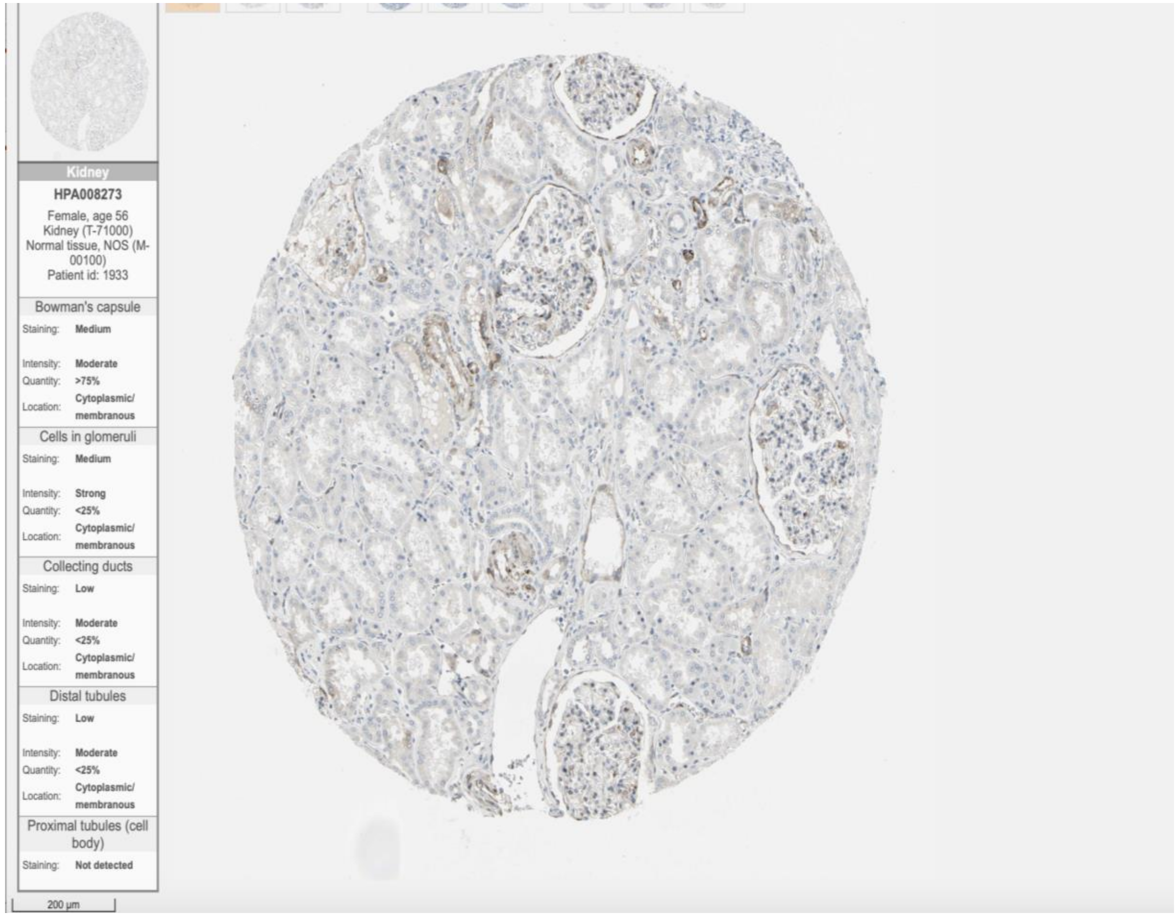

Kidney
HPA008273
Female, age 56
Kidney (T-71000)
Normal tissue, NOS (M-00100)
Patient id: 1933
Bowman's capsule
Staining: Medium
Intensity: Moderate
Quantity: >75%
Location: Cytoplasmic/ membranous
Cells in glomeruli
Staining: Medium
Intensity: Strong
Quantity: <25%
Location: Cytoplasmic/ membranous
Collecting ducts
Staining: Low
Intensity: Moderate
Quantity: <25%
Location: Cytoplasmic/ membranous
Distal tubules
Staining: Low
Intensity: Moderate
Quantity: <25%
Location: Cytoplasmic/ membranous
Proximal tubules (cell body)
Staining: Not detected
200 µm


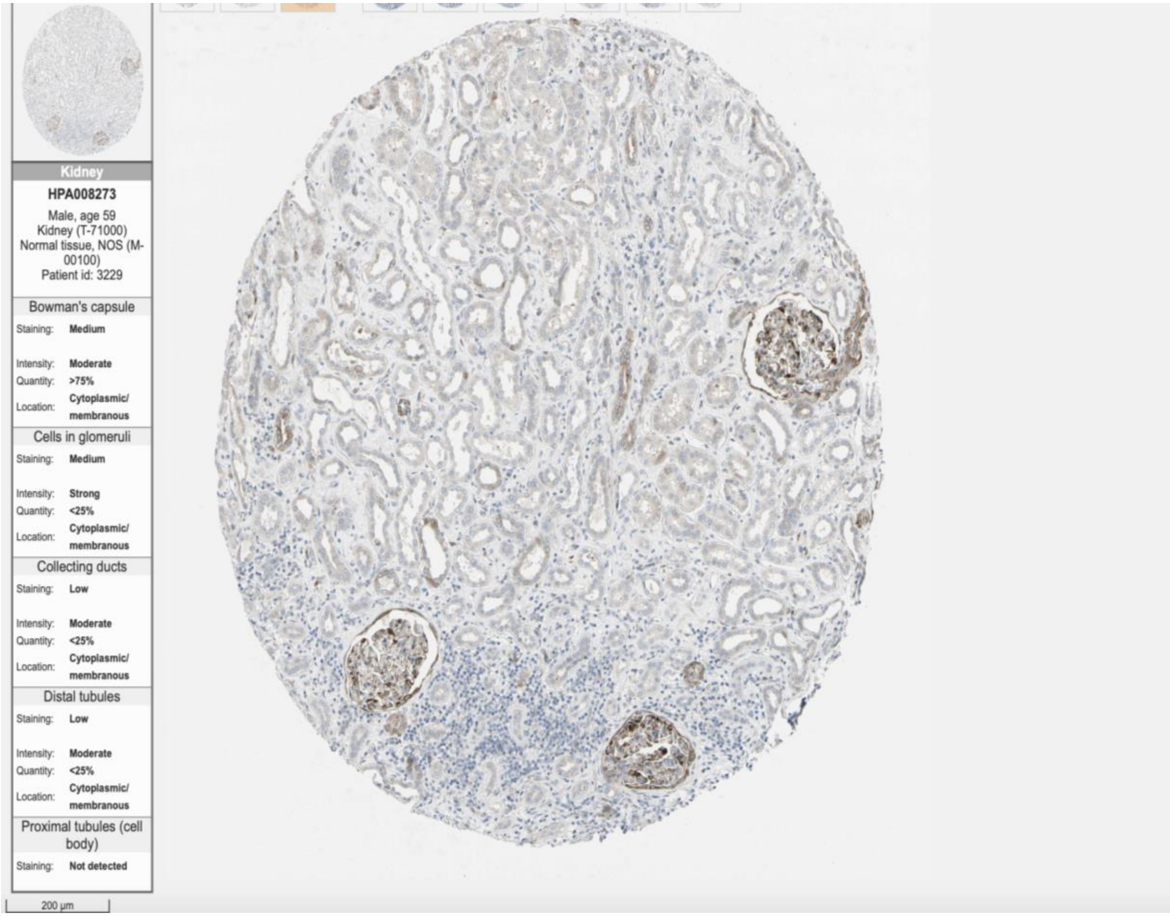

Kidney
HPA008273
Male, age 59
Kidney (T-71000)
Normal tissue, NOS (M-00100)
Patient id: 3229
Bowman's capsule
Staining: Medium
Intensity: Moderate
Quantity: >75%
Location: Cytoplasmic/ membranous
Cells in glomeruli
Staining: Medium
Intensity: Strong
Quantity: <25%
Location: Cytoplasmic/ membranous
Collecting ducts
Staining: Low
Intensity: Moderate
Quantity: <25%
Location: Cytoplasmic/ membranous
Distal tubules
Staining: Low
Intensity: Moderate
Quantity: <25%
Location: Cytoplasmic/ membranous
Proximal tubules (cell body)
Staining: Not detected
200 µm

BDNF

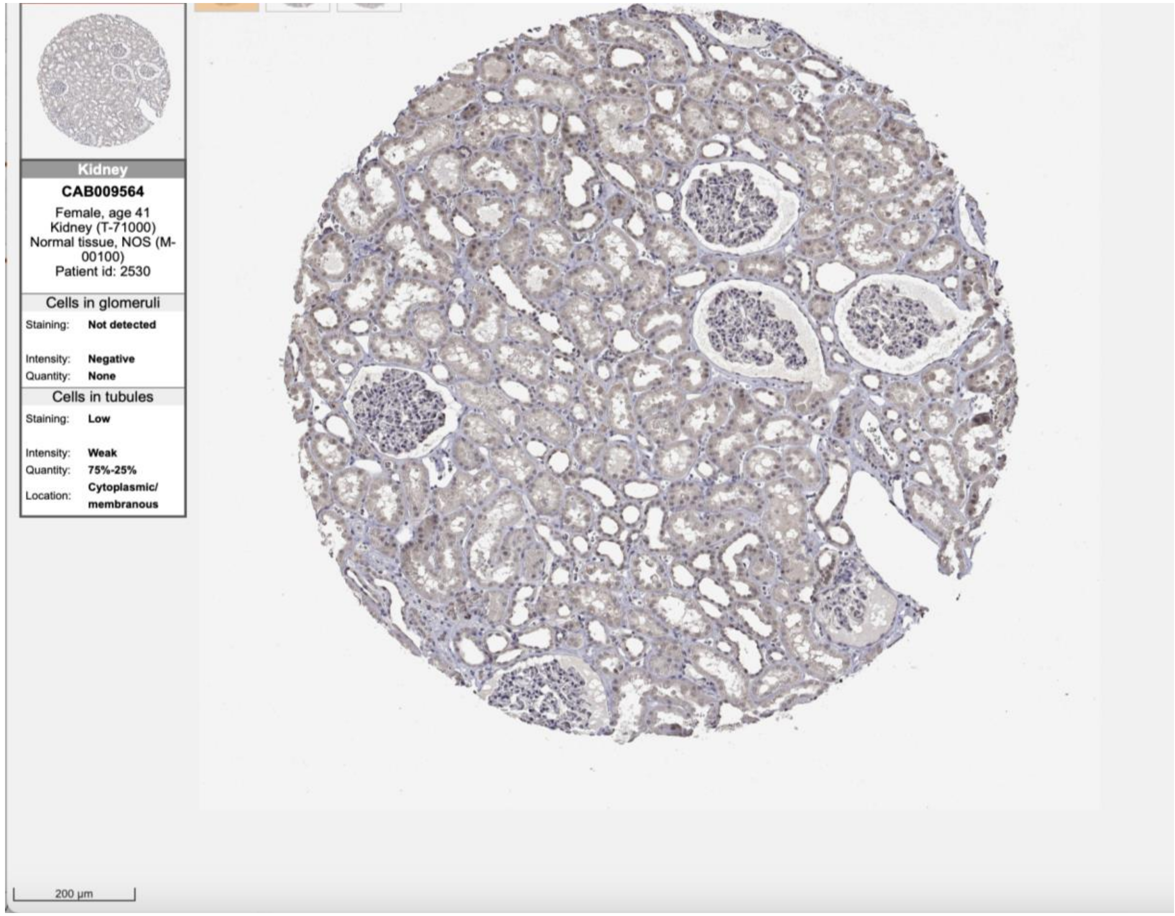
Kidney
CAB009564
Female, age 41
Kidney (T-71000)
Normal tissue, NOS (M-00100)
Patient id: 2530
Cells in glomeruli
Staining: Not detected
Intensity: Negative
Quantity: None
Cells in tubules
Staining: Low
Intensity: Weak
Quantity: 75%-25%
Location: Cytoplasmic/ membranous
200 µm

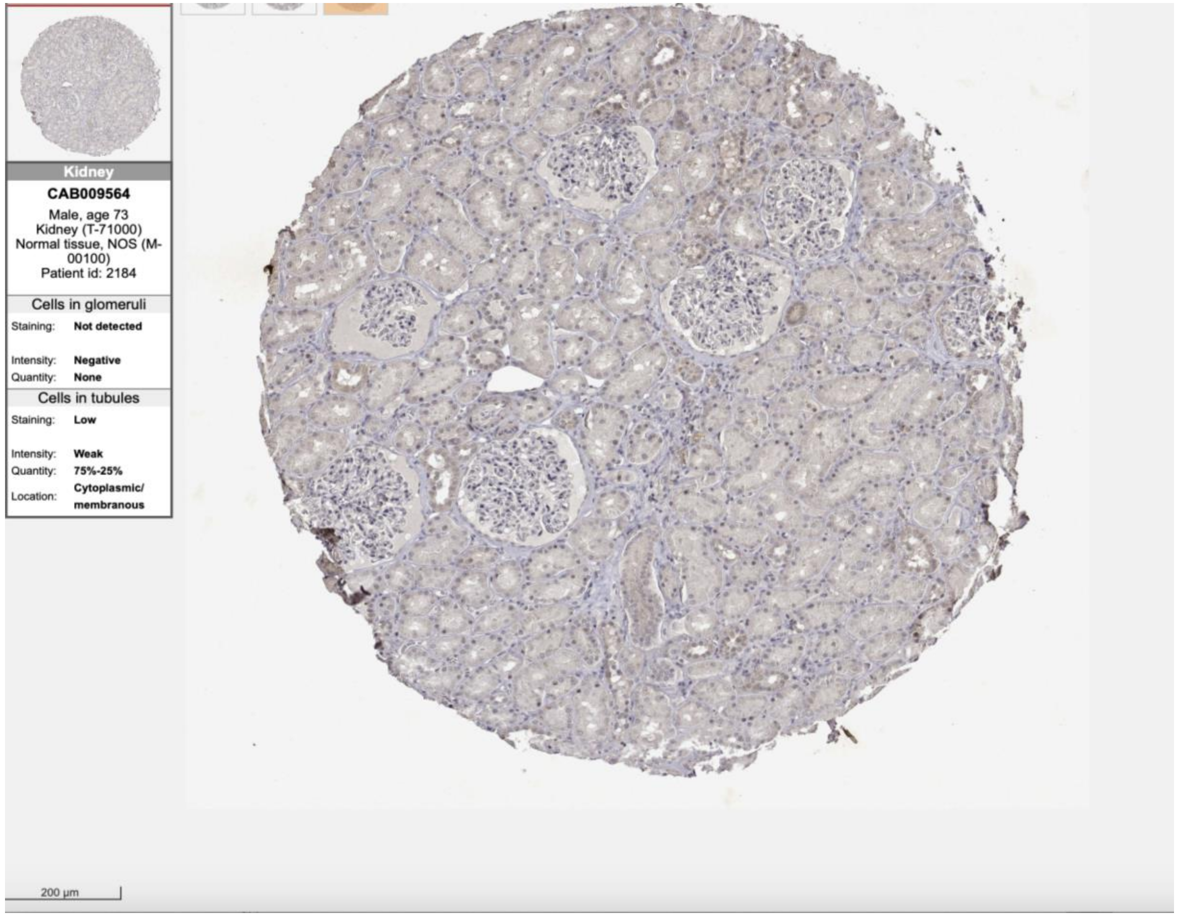
Kidney
CAB009564
Male, age 73
Kidney (T-71000)
Normal tissue, NOS (M-00100)
Patient id: 2184
Cells in glomeruli
Staining: Not detected
Intensity: Negative
Quantity: None
Cells in tubules
Staining: Low
Intensity: Weak
Quantity: 75%-25%
Location: Cytoplasmic/ membranous
200 µm

**TUBB**

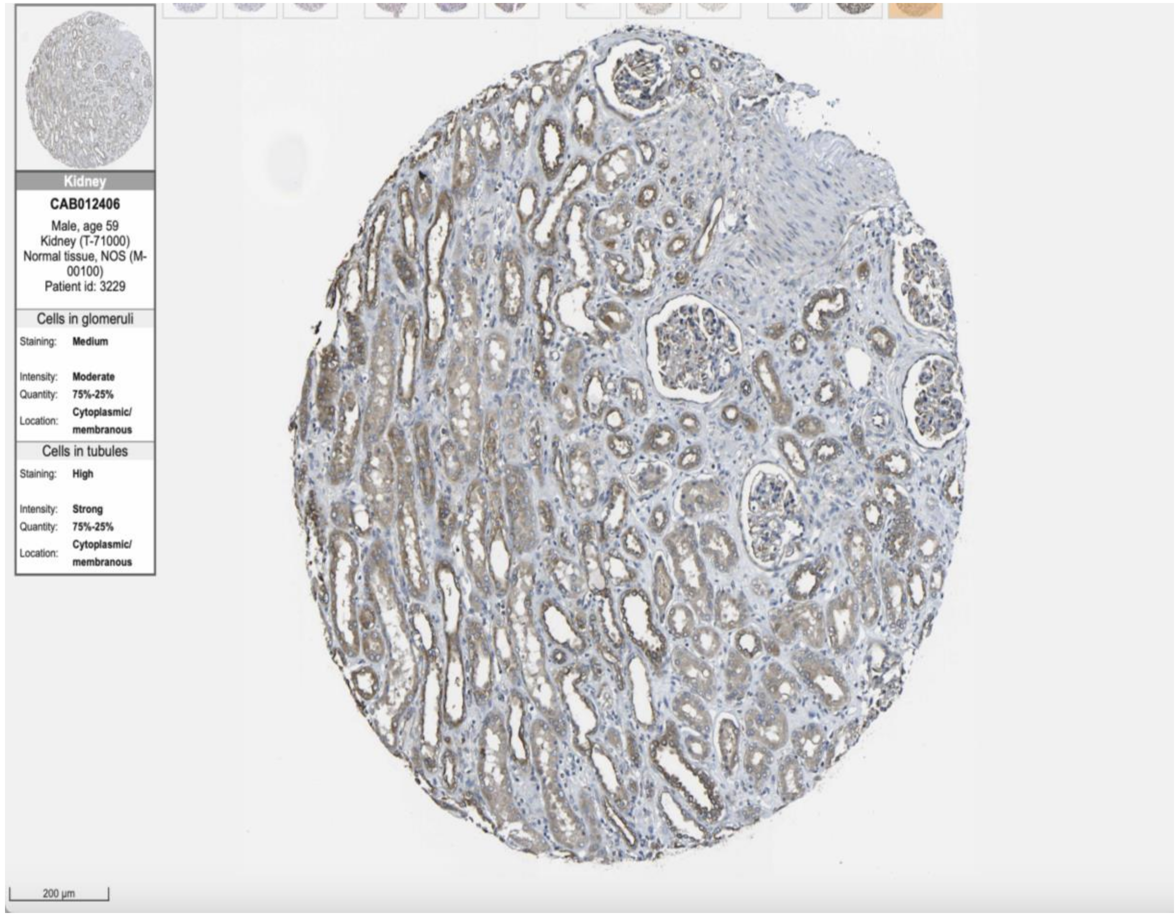

Kidney
CAB012406
Male, age 59
Kidney (T-71000)
Normal tissue, NOS (M-00100)
Patient id: 3229
Cells in glomeruli
Staining: Medium
Intensity: Moderate
Quantity: 75%-25%
Location: Cytoplasmic/ membranous
Cells in tubules
Staining: High
Intensity: Strong
Quantity: 75%-25%
Location: Cytoplasmic/ membranous
200 µm


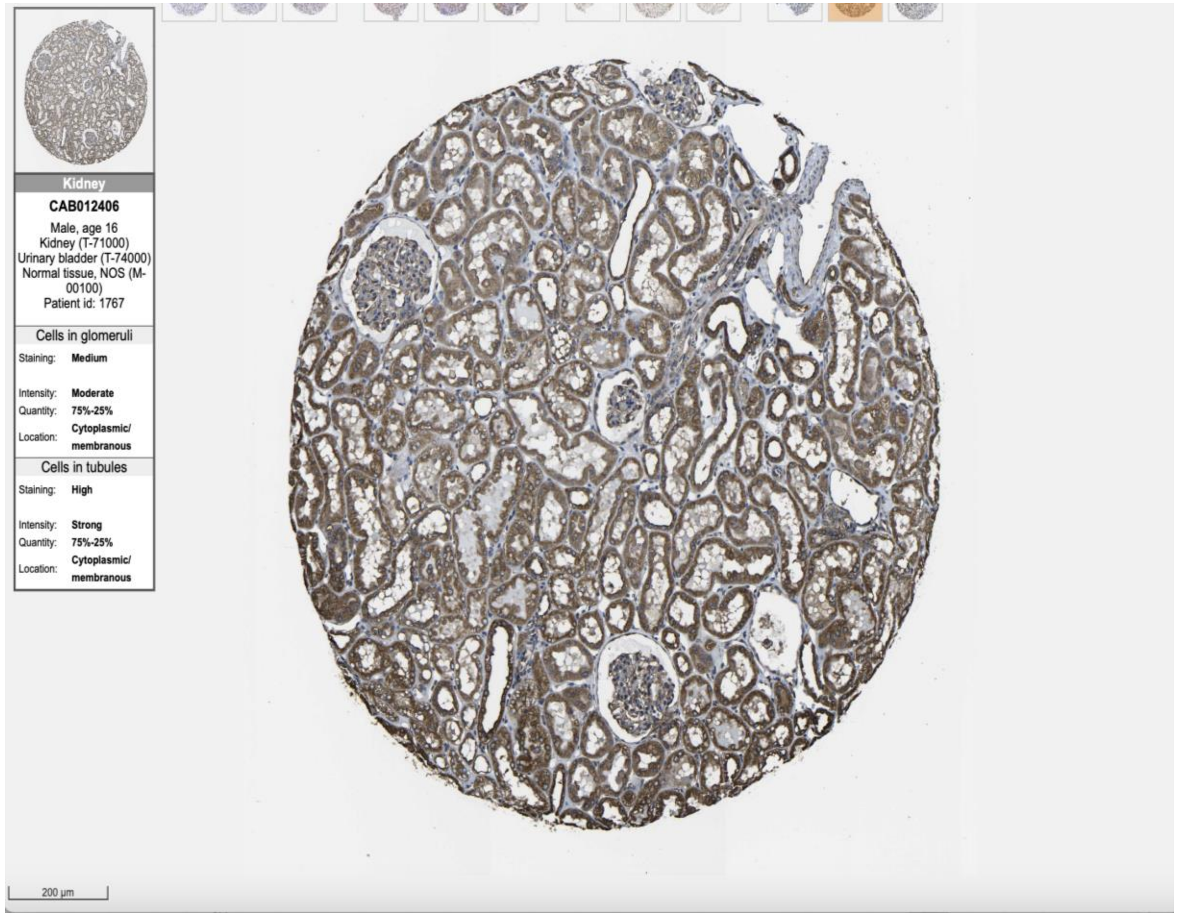

Kidney
CAB012406
Male, age 16
Kidney (T-71000)
Urinary bladder (T-74000)
Normal tissue, NOS (M-00100)
Patient id: 1767
Cells in glomeruli
Staining: Medium
Intensity: Moderate
Quantity: 75%-25%
Location: Cytoplasmic/ membranous
Cells in tubules
Staining: High
Intensity: Strong
Quantity: 75%-25%
Location: Cytoplasmic/ membranous
200 µm

RSPH1

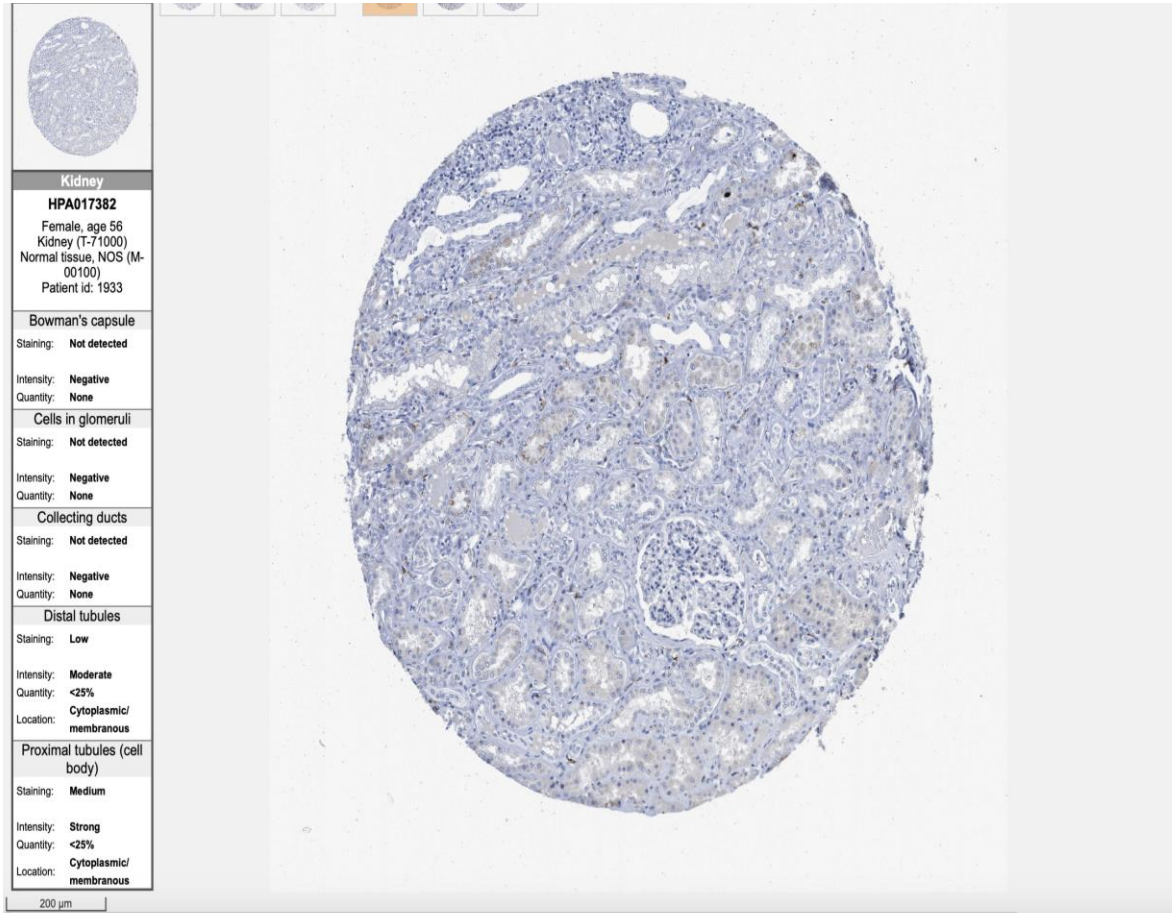
Kidney
HPA017382
Female, age 56
Kidney (T-71000)
Normal tissue, NOS (M-00100)
Patient id: 1933
Bowman's capsule
Staining: Not detected
Intensity: Negative
Quantity: None
Cells in glomeruli
Staining: Not detected
Intensity: Negative
Quantity: None
Collecting ducts
Staining: Not detected
Intensity: Negative
Quantity: None
Distal tubules
Staining: Low
Intensity: Moderate
Quantity: <25%
Location: Cytoplasmic/ membranous
Proximal tubules (cell body)
Staining: Medium
Intensity: Strong
Quantity: <25%
Location: Cytoplasmic/ membranous
200 µm

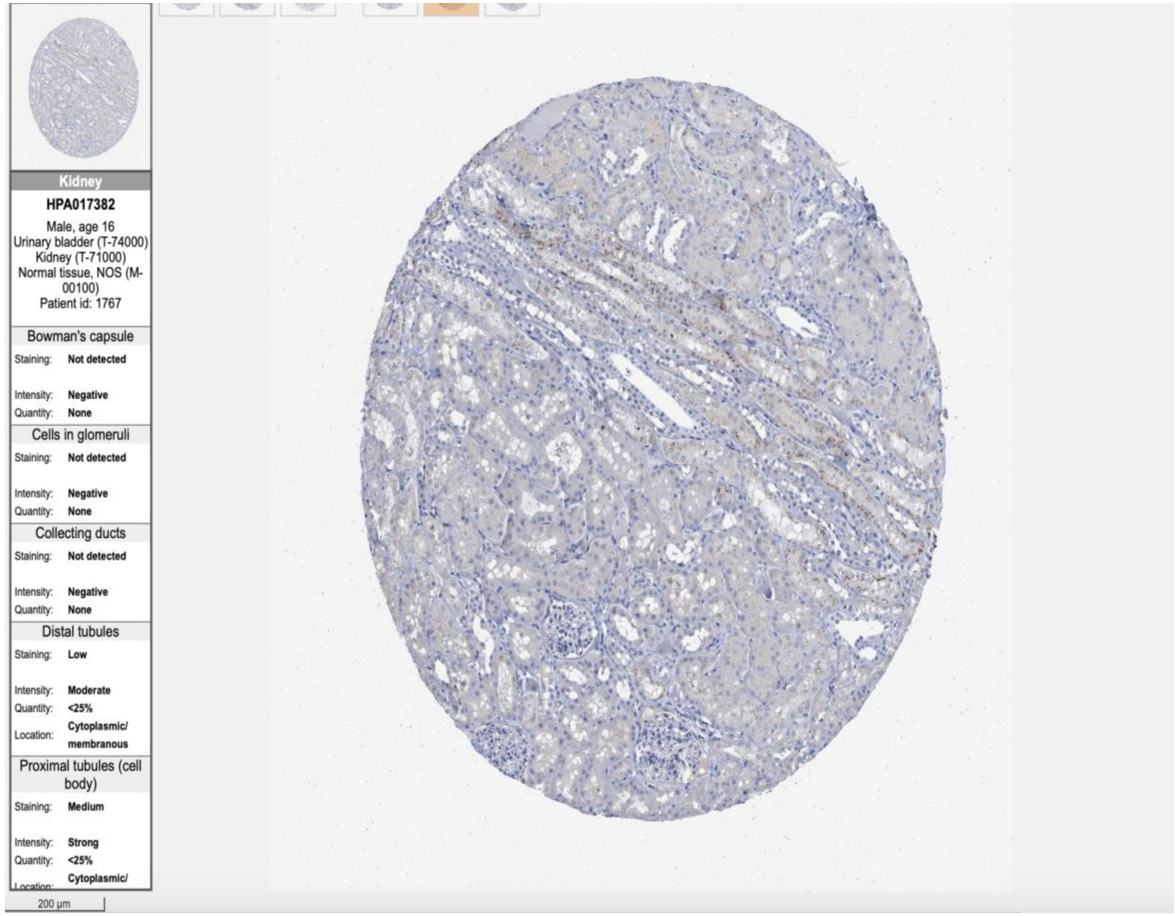
Kidney
HPA017382
Male, age 16
Urinary bladder (T-74000)
Kidney (T-71000)
Normal tissue, NOS (M-00100)
Patient id: 1767
Bowman's capsule
Staining: Not detected
Intensity: Negative
Quantity: None
Cells in glomeruli
Staining: Not detected
Intensity: Negative
Quantity: None
Collecting ducts
Staining: Not detected
Intensity: Negative
Quantity: None
Distal tubules
Staining: Low
Intensity: Moderate
Quantity: <25%
Location: Cytoplasmic/ membranous
Proximal tubules (cell body)
Staining: Medium
Intensity: Strong
Quantity: <25%
Cytoplasmic/
200 µm

SH3GL2

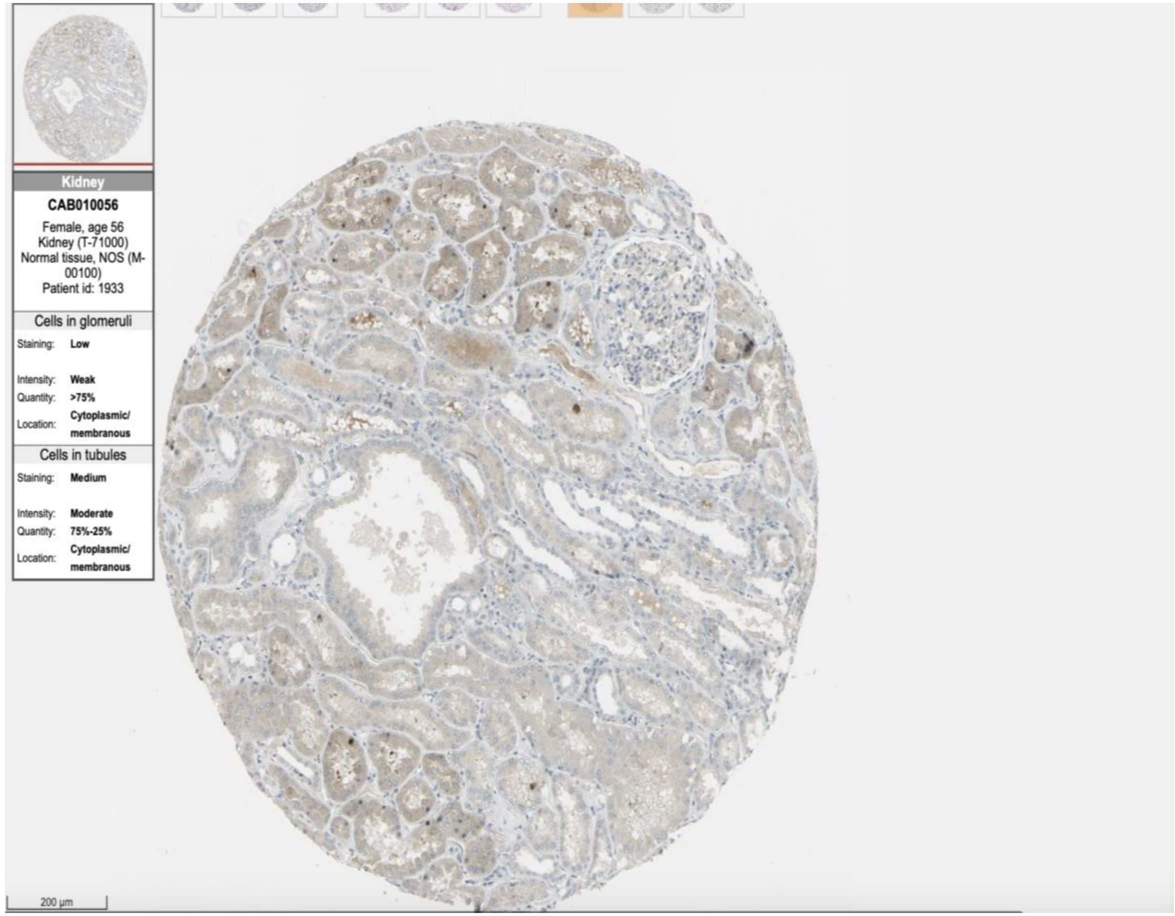
Kidney
CAB010056
Female, age 56
Kidney (T-71000)
Normal tissue, NOS (M-00100)
Patient id: 1933
Cells in glomeruli
Staining: Low
Intensity: Weak
Quantity: >75%
Location: Cytoplasmic/membranous
Cells in tubules
Staining: Medium
Intensity: Moderate
Quantity: 75%-25%
Location: Cytoplasmic/membranous
200 µm

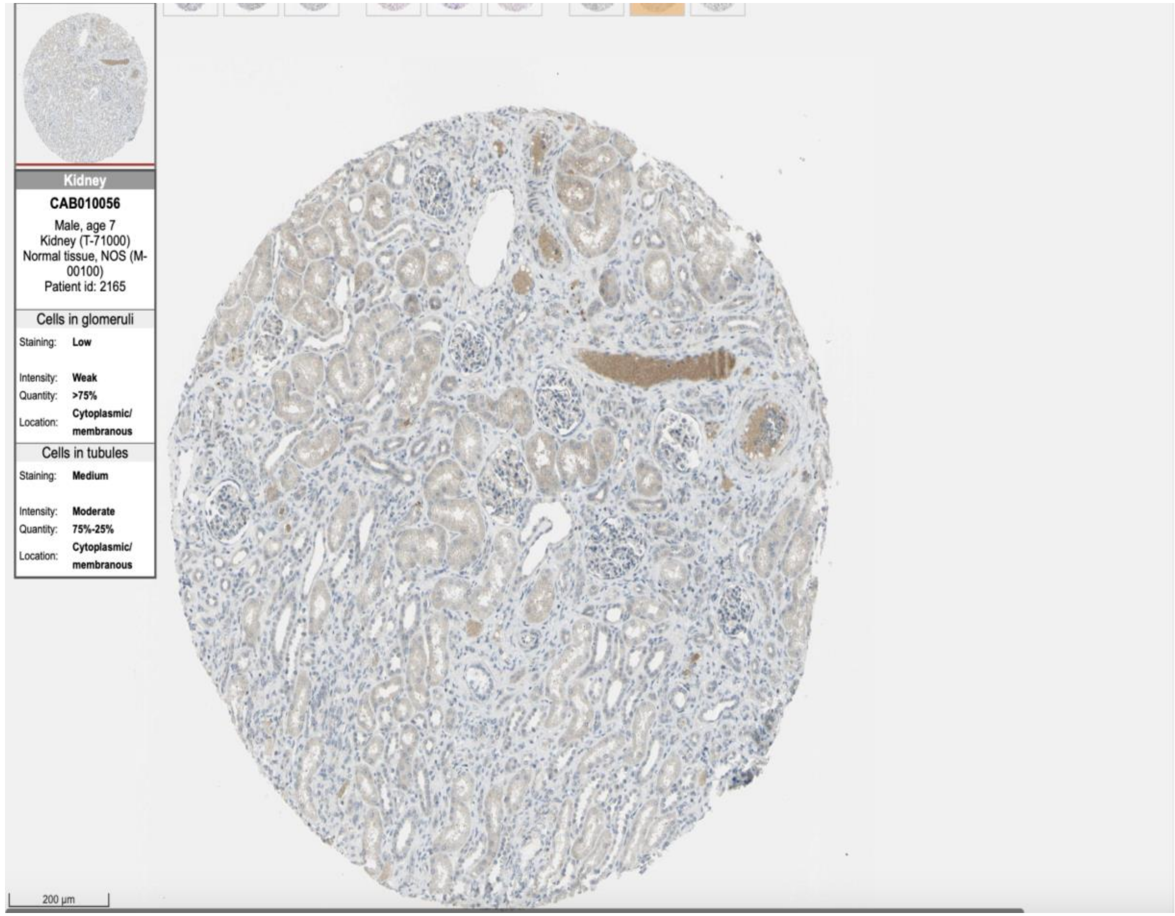
Kidney
CAB010056
Male, age 7
Kidney (T-71000)
Normal tissue, NOS (M-00100)
Patient id: 2165
Cells in glomeruli
Staining: Low
Intensity: Weak
Quantity: >75%
Location: Cytoplasmic/membranous
Cells in tubules
Staining: Medium
Intensity: Moderate
Quantity: 75%-25%
Location: Cytoplasmic/membranous
200 µm

NPHS1

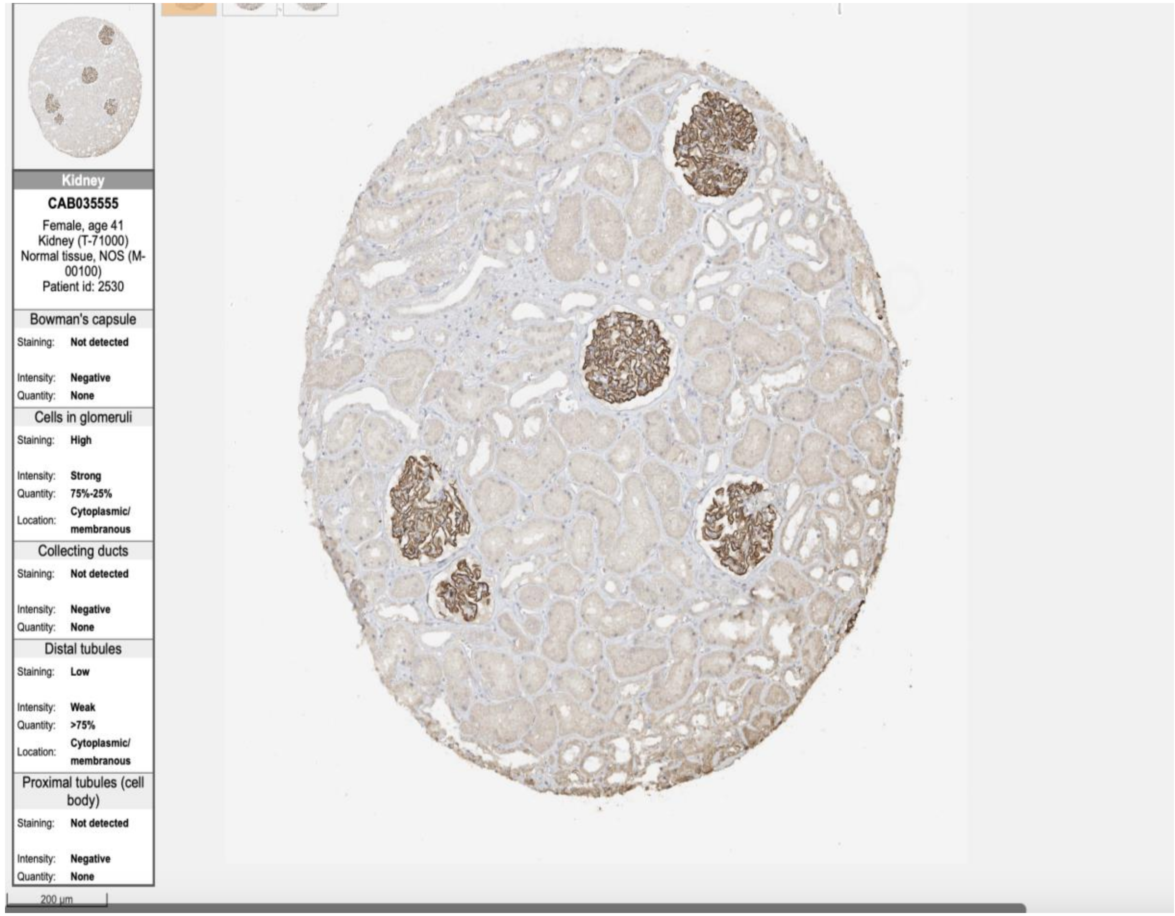

Kidney
CAB035555
Female, age 41
Kidney (T-71000)
Normal tissue, NOS (M-00100)
Patient id: 2530
Bowman's capsule
Staining: Not detected
Intensity: Negative
Quantity: None
Cells in glomeruli
Staining: High
Intensity: Strong
Quantity: 75%-25%
Location: Cytoplasmic/ membranous
Collecting ducts
Staining: Not detected
Intensity: Negative
Quantity: None
Distal tubules
Staining: Low
Intensity: Weak
Quantity: >75%
Location: Cytoplasmic/ membranous
Proximal tubules (cell body)
Staining: Not detected
Intensity: Negative
Quantity: None
200 µm


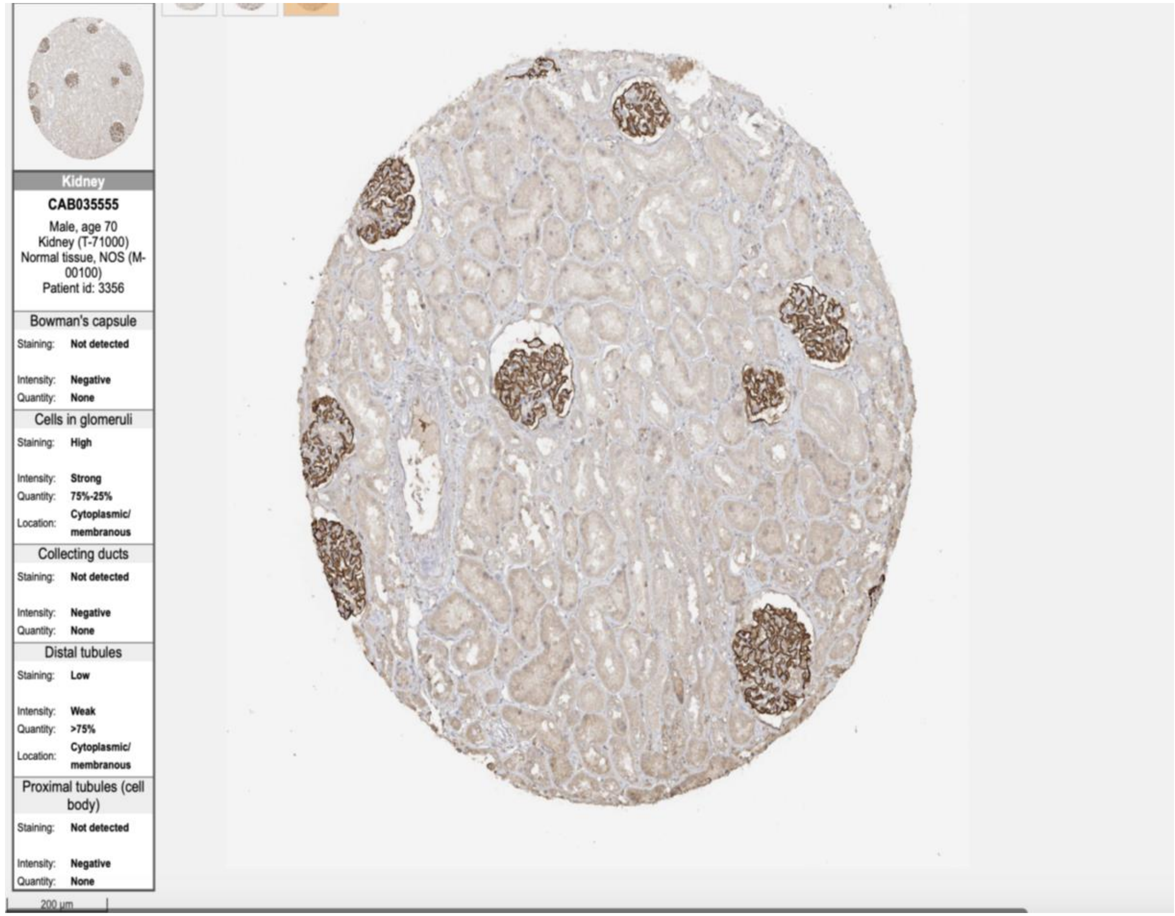

Kidney
CAB035555
Male, age 70
Kidney (T-71000)
Normal tissue, NOS (M-00100)
Patient id: 3356
Bowman's capsule
Staining: Not detected
Intensity: Negative
Quantity: None
Cells in glomeruli
Staining: High
Intensity: Strong
Quantity: 75%-25%
Location: Cytoplasmic/ membranous
Collecting ducts
Staining: Not detected
Intensity: Negative
Quantity: None
Distal tubules
Staining: Low
Intensity: Weak
Quantity: >75%
Location: Cytoplasmic/ membranous
Proximal tubules (cell body)
Staining: Not detected
Intensity: Negative
Quantity: None
200 µm

SYNPO

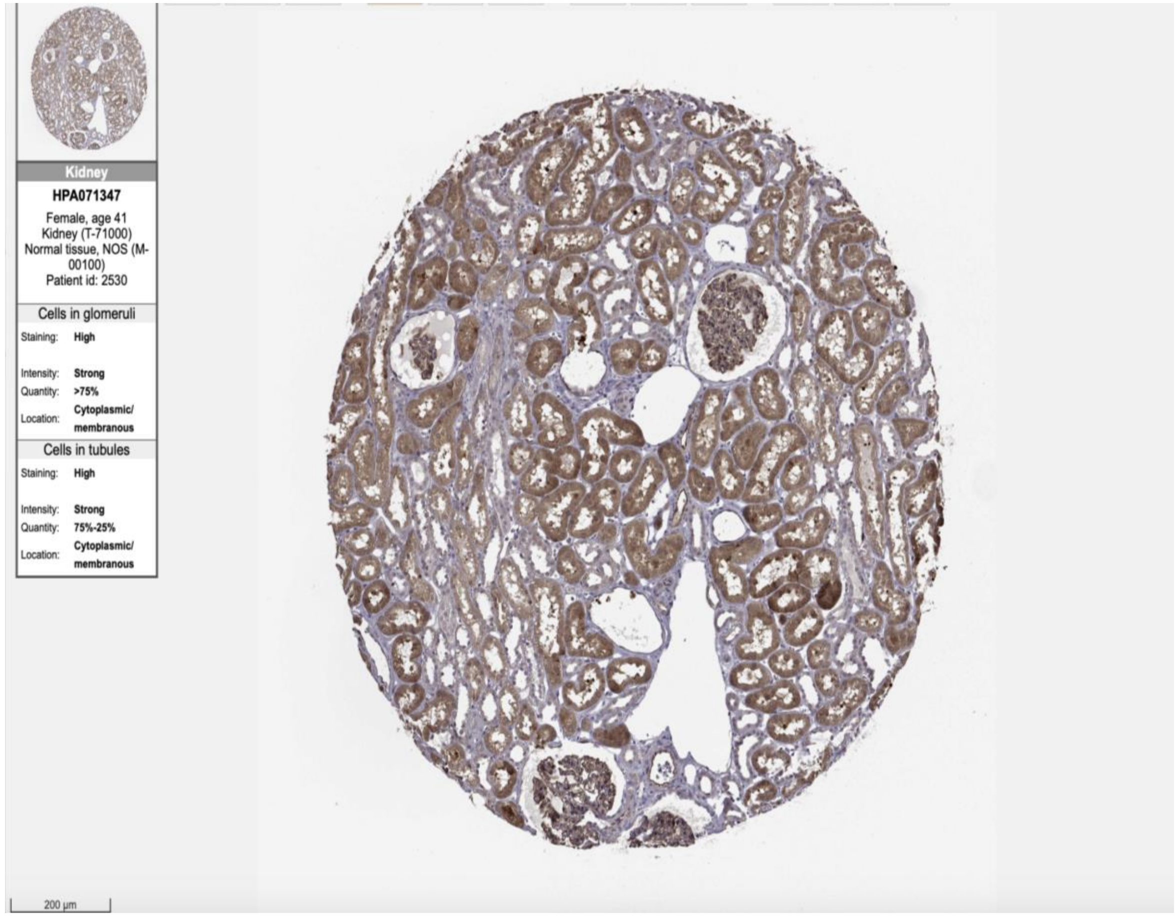
Kidney
HPA071347
Female, age 41
Kidney (T-71000)
Normal tissue, NOS (M-00100)
Patient id: 2530
Cells in glomeruli
Staining: High
Intensity: Strong
Quantity: >75%
Location: Cytoplasmic/ membranous
Cells in tubules
Staining: High
Intensity: Strong
Quantity: 75%-25%
Location: Cytoplasmic/ membranous
200 µm

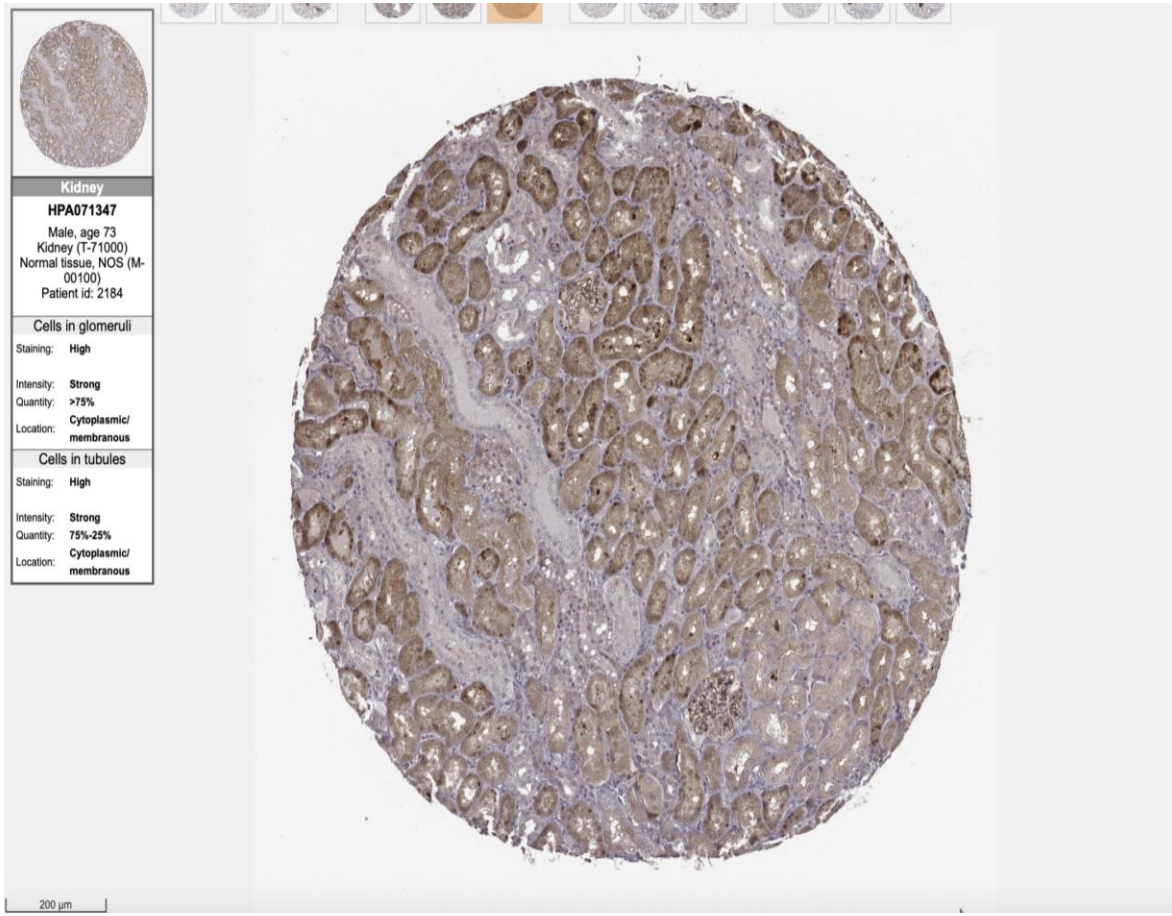
Kidney
HPA071347
Male, age 73
Kidney (T-71000)
Normal tissue, NOS (M-00100)
Patient id: 2184
Cells in glomeruli
Staining: High
Intensity: Strong
Quantity: >75%
Location: Cytoplasmic/ membranous
Cells in tubules
Staining: High
Intensity: Strong
Quantity: 75%-25%
Location: Cytoplasmic/ membranous
200 µm

### Podocyte/Brain

transcription regulator activity (TF´s)

ATF7IP2

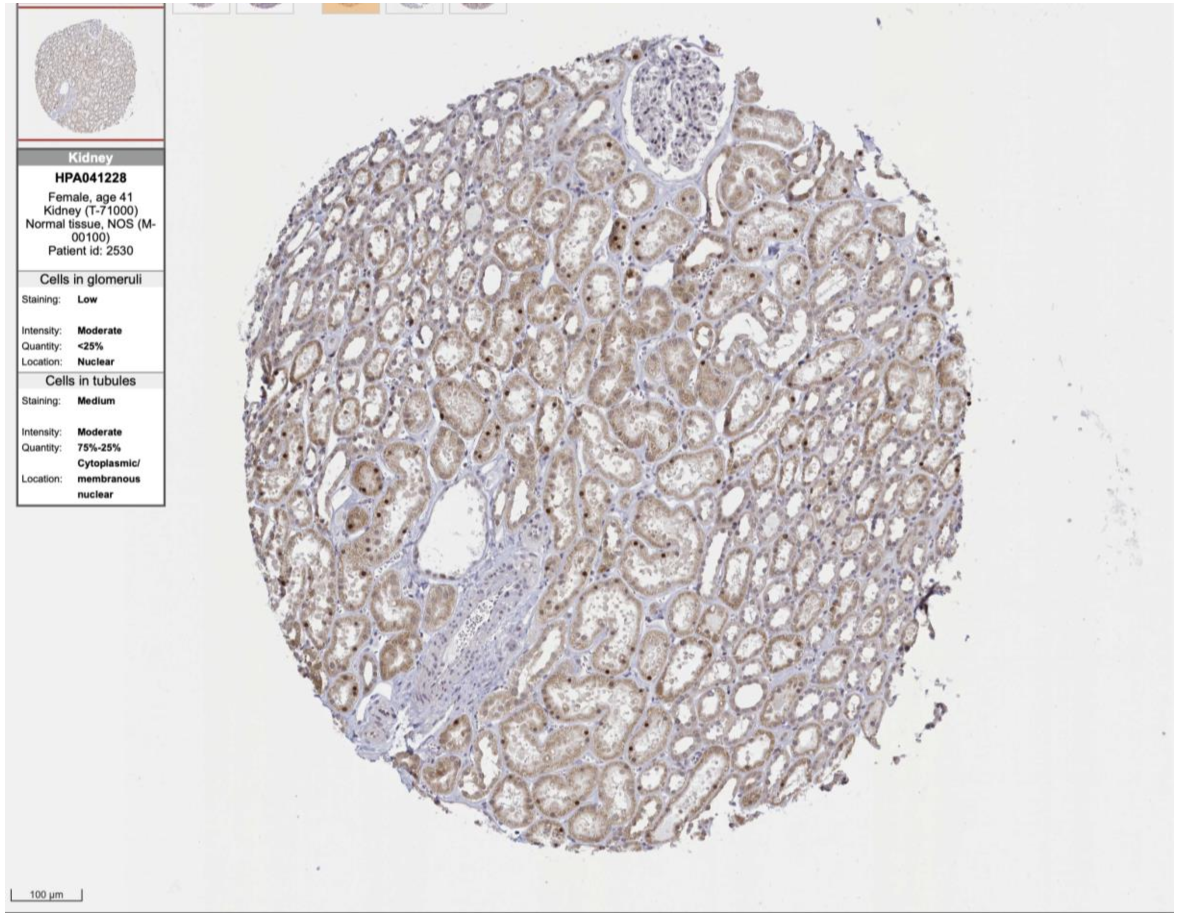
Kidney
HPA041228
Female, age 41
Kidney (T-71000)
Normal tissue, NOS (M-00100)
Patient id: 2530
Cells in glomeruli
Staining: Low
Intensity: Moderate
Quantity: <25%
Location: Nuclear
Cells in tubules
Staining: Medium
Intensity: Moderate
Quantity: 75%-25%
Location: Cytoplasmic/ membranous nuclear
100 µm

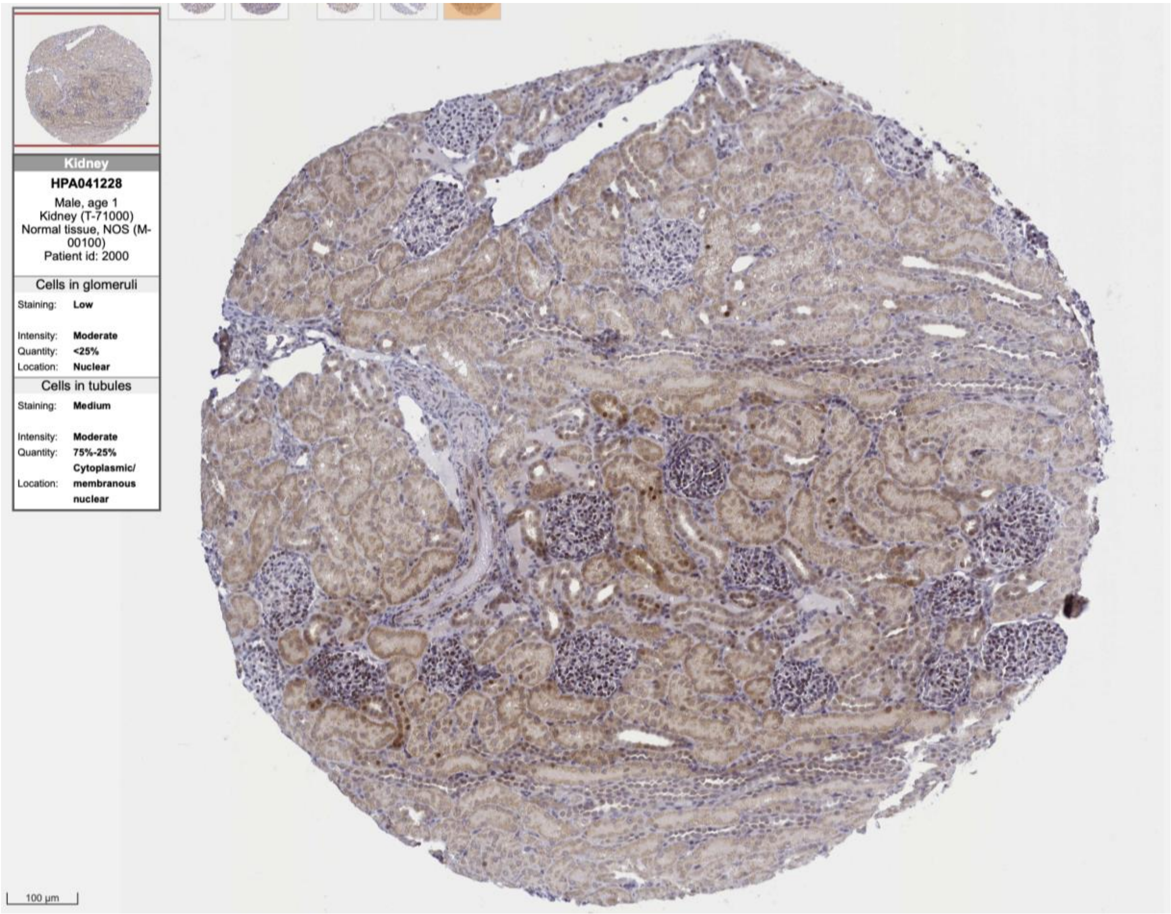
Kidney
HPA041228
Male, age 1
Kidney (T-71000)
Normal tissue, NOS (M-00100)
Patient id: 2000
Cells in glomeruli
Staining: Low
Intensity: Moderate
Quantity: <25%
Location: Nuclear
Cells in tubules
Staining: Medium
Intensity: Moderate
Quantity: 75%-25%
Location: Cytoplasmic/ membranous nuclear
100 µm

BACH2

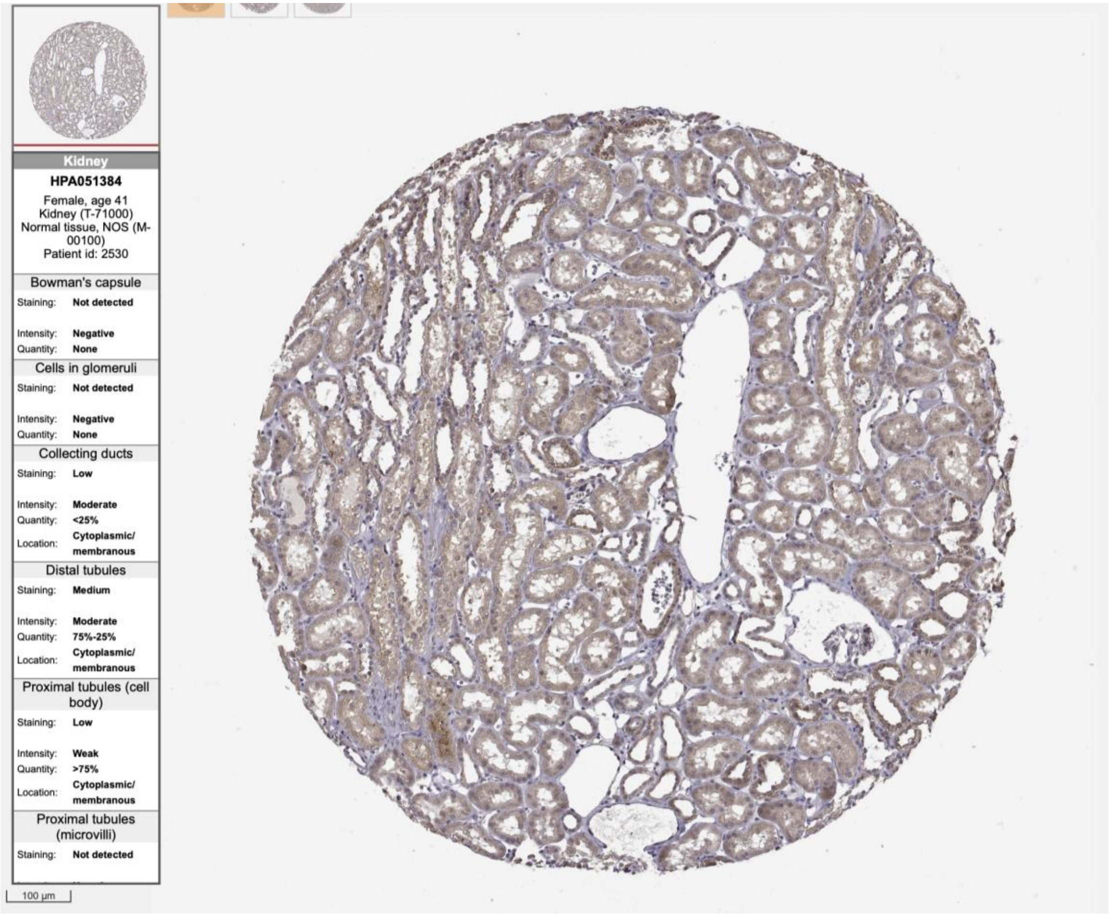
Kidney
HPA051384
Female, age 41
Kidney (T-71000)
Normal tissue, NOS (M-00100)
Patient id: 2530
Bowman's capsule
Staining: Not detected
Intensity: Negative
Quantity: None
Cells in glomeruli
Staining: Not detected
Intensity: Negative
Quantity: None
Collecting ducts
Staining: Low
Intensity: Moderate
Quantity: <25%
Location: Cytoplasmic/ membranous
Distal tubules
Staining: Medium
Intensity: Moderate
Quantity: 75%-25%
Location: Cytoplasmic/ membranous
Proximal tubules (cell body)
Staining: Low
Intensity: Weak
Quantity: >75%
Location: Cytoplasmic/ membranous
Proximal tubules (microvilli)
Staining: Not detected
100 µm

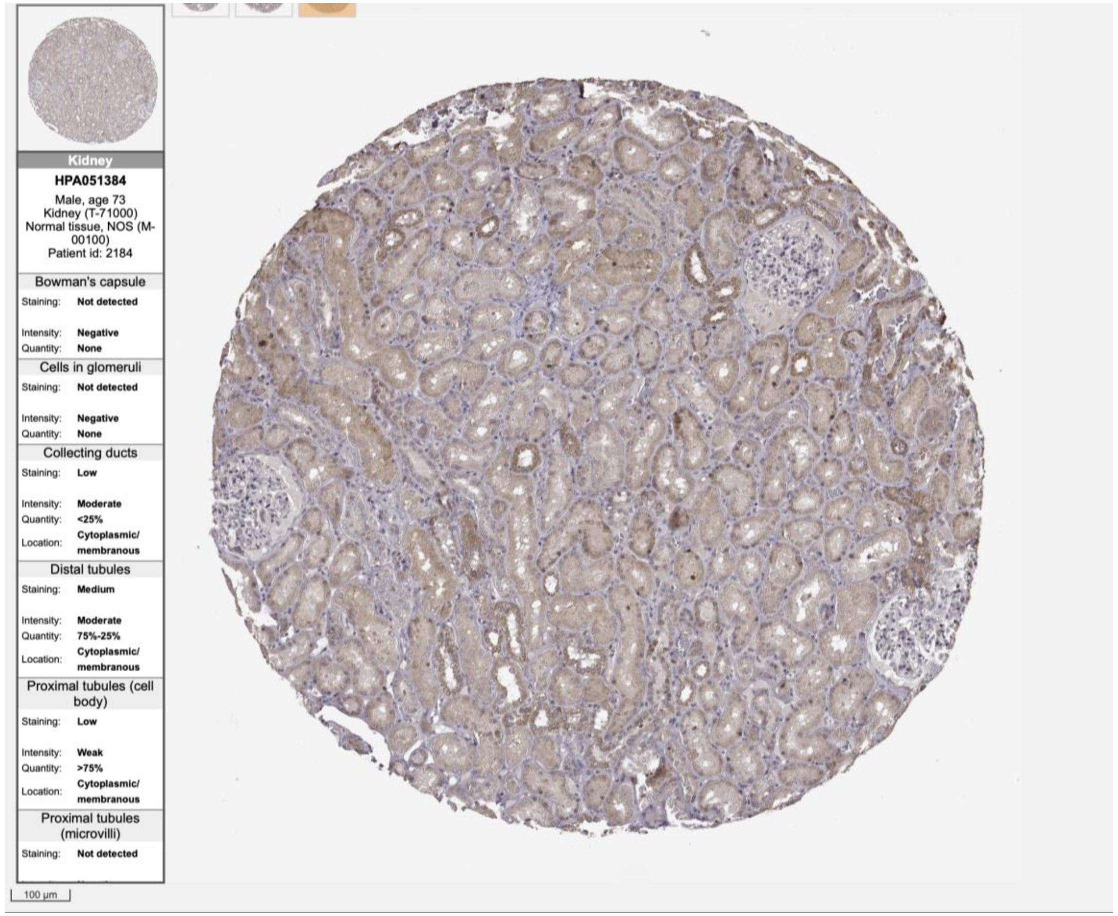
Kidney
HPA051384
Male, age 73
Kidney (T-71000)
Normal tissue, NOS (M-00100)
Patient id: 2184
Bowman's capsule
Staining: Not detected
Intensity: Negative
Quantity: None
Cells in glomeruli
Staining: Not detected
Intensity: Negative
Quantity: None
Collecting ducts
Staining: Low
Intensity: Moderate
Quantity: <25%
Location: Cytoplasmic/ membranous
Distal tubules
Staining: Medium
Intensity: Moderate
Quantity: 75%-25%
Location: Cytoplasmic/ membranous
Proximal tubules (cell body)
Staining: Low
Intensity: Weak
Quantity: >75%
Location: Cytoplasmic/ membranous
Proximal tubules (microvilli)
Staining: Not detected
100 µm

EAF2

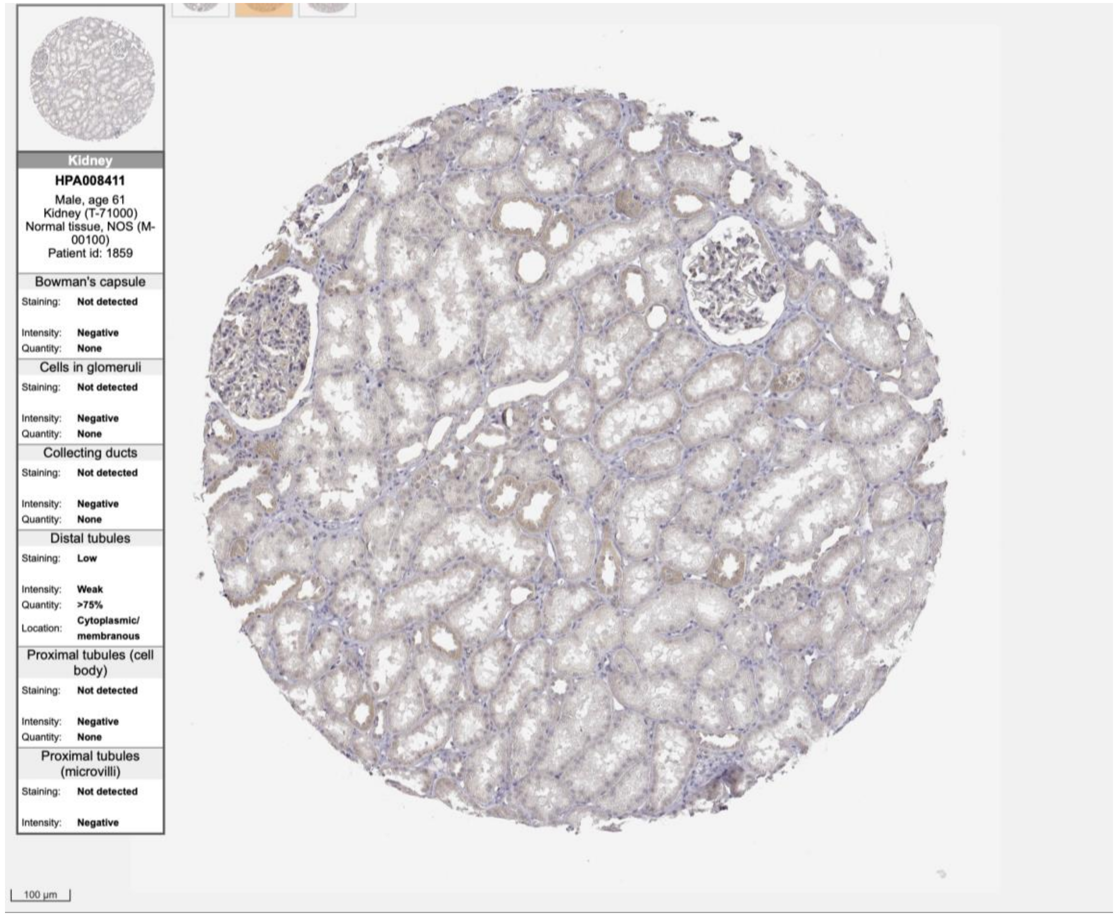
Kidney
HPA008411
Male, age 61
Kidney (T-71000)
Normal tissue, NOS (M-00100)
Patient id: 1859
Bowman's capsule
Staining: Not detected
Intensity: Negative
Quantity: None
Cells in glomeruli
Staining: Not detected
Intensity: Negative
Quantity: None
Collecting ducts
Staining: Not detected
Intensity: Negative
Quantity: None
Distal tubules
Staining: Low
Intensity: Weak
Quantity: >75%
Location: Cytoplasmic/ membranous
Proximal tubules (cell body)
Staining: Not detected
Intensity: Negative
Quantity: None
Proximal tubules (microvilli)
Staining: Not detected
Intensity: Negative
100 µm

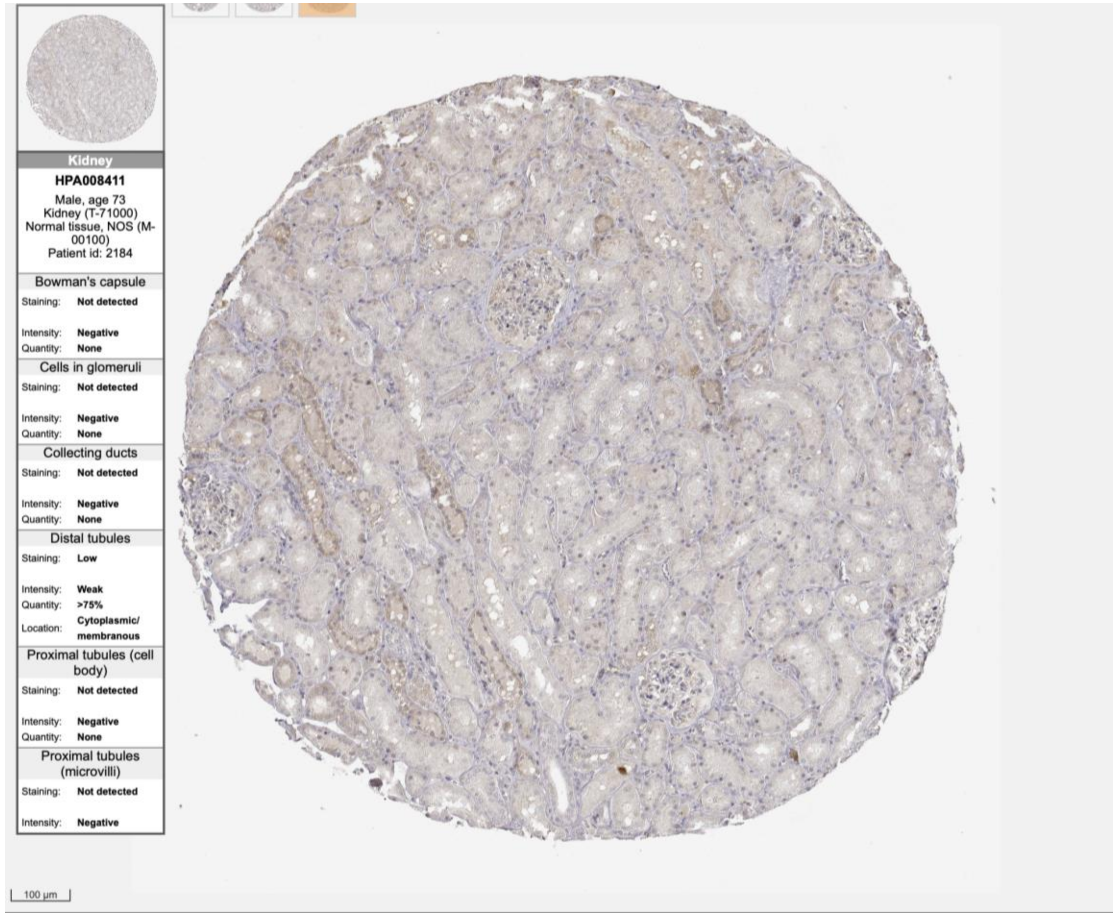
Kidney
HPA008411
Male, age 73
Kidney (T-71000)
Normal tissue, NOS (M-00100)
Patient id: 2184
Bowman's capsule
Staining: Not detected
Intensity: Negative
Quantity: None
Cells in glomeruli
Staining: Not detected
Intensity: Negative
Quantity: None
Collecting ducts
Staining: Not detected
Intensity: Negative
Quantity: None
Distal tubules
Staining: Low
Intensity: Weak
Quantity: >75%
Location: Cytoplasmic/ membranous
Proximal tubules (cell body)
Staining: Not detected
Intensity: Negative
Quantity: None
Proximal tubules (microvilli)
Staining: Not detected
Intensity: Negative
100 µm

**FOXD3**

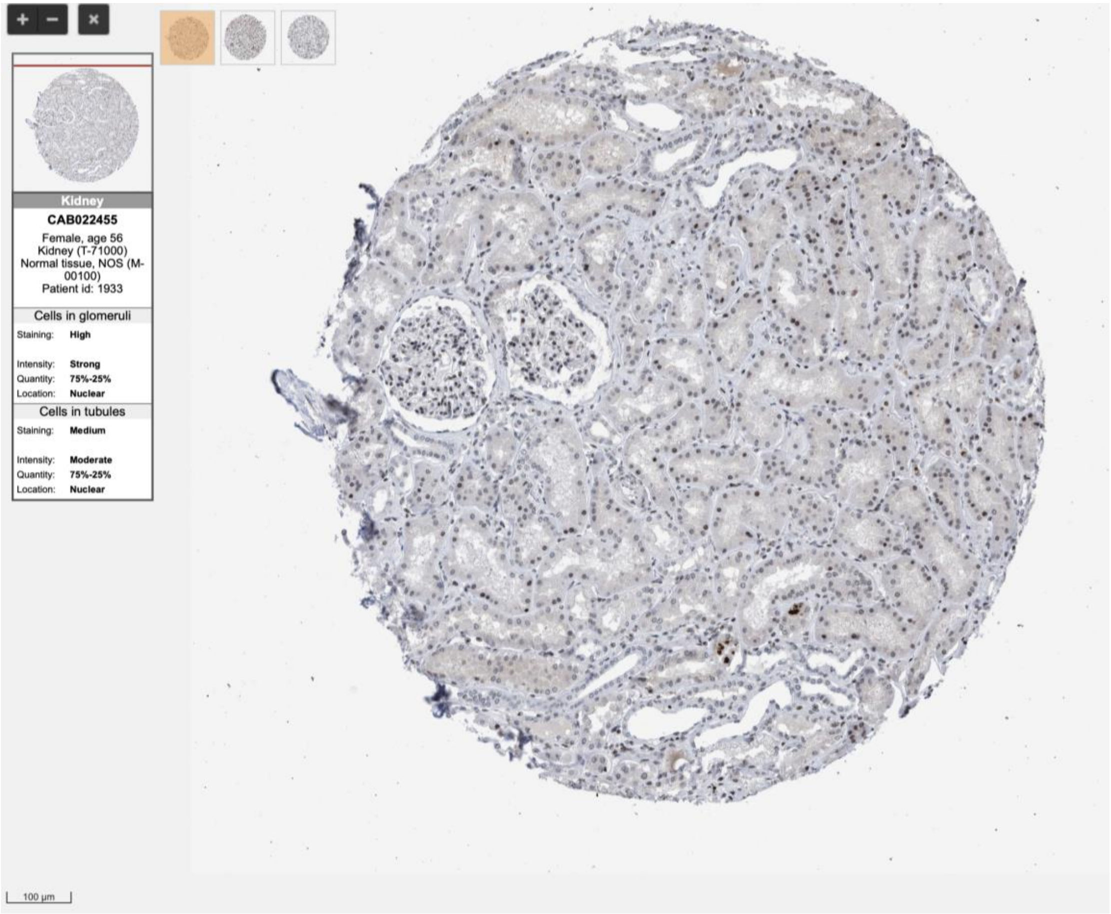
Kidney
CAB022455
Female, age 56
Kidney (T-71000)
Normal tissue, NOS (M-00100)
Patient id: 1933
Cells in glomeruli
Staining: High
Intensity: Strong
Quantity: 75%-25%
Location: Nuclear
Cells in tubules
Staining: Medium
Intensity: Moderate
Quantity: 75%-25%
Location: Nuclear
100 µm

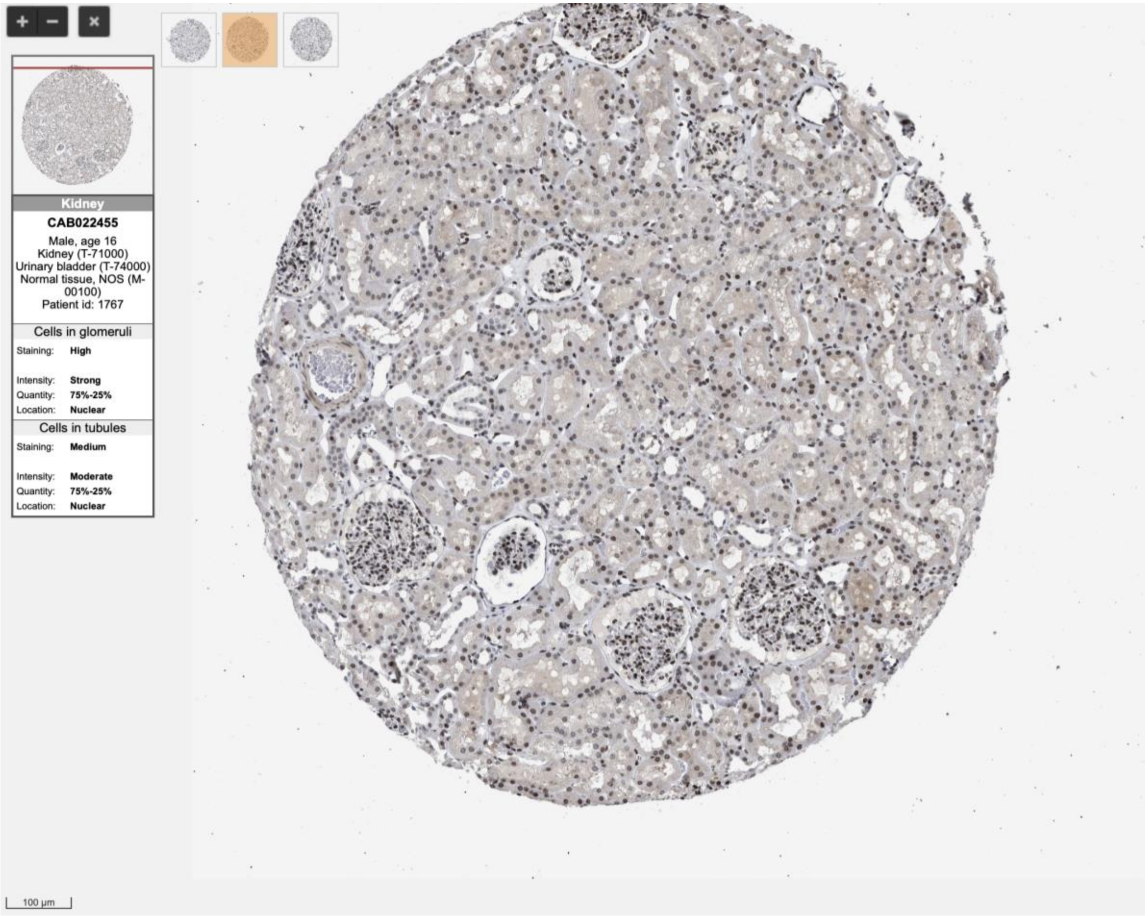
Kidney
CAB022455
Male, age 16
Kidney (T-71000)
Urinary bladder (T-74000)
Normal tissue, NOS (M-00100)
Patient id: 1767
Cells in glomeruli
Staining: High
Intensity: Strong
Quantity: 75%-25%
Location: Nuclear
Cells in tubules
Staining: Medium
Intensity: Moderate
Quantity: 75%-25%
Location: Nuclear
100 µm

**FOXS1**

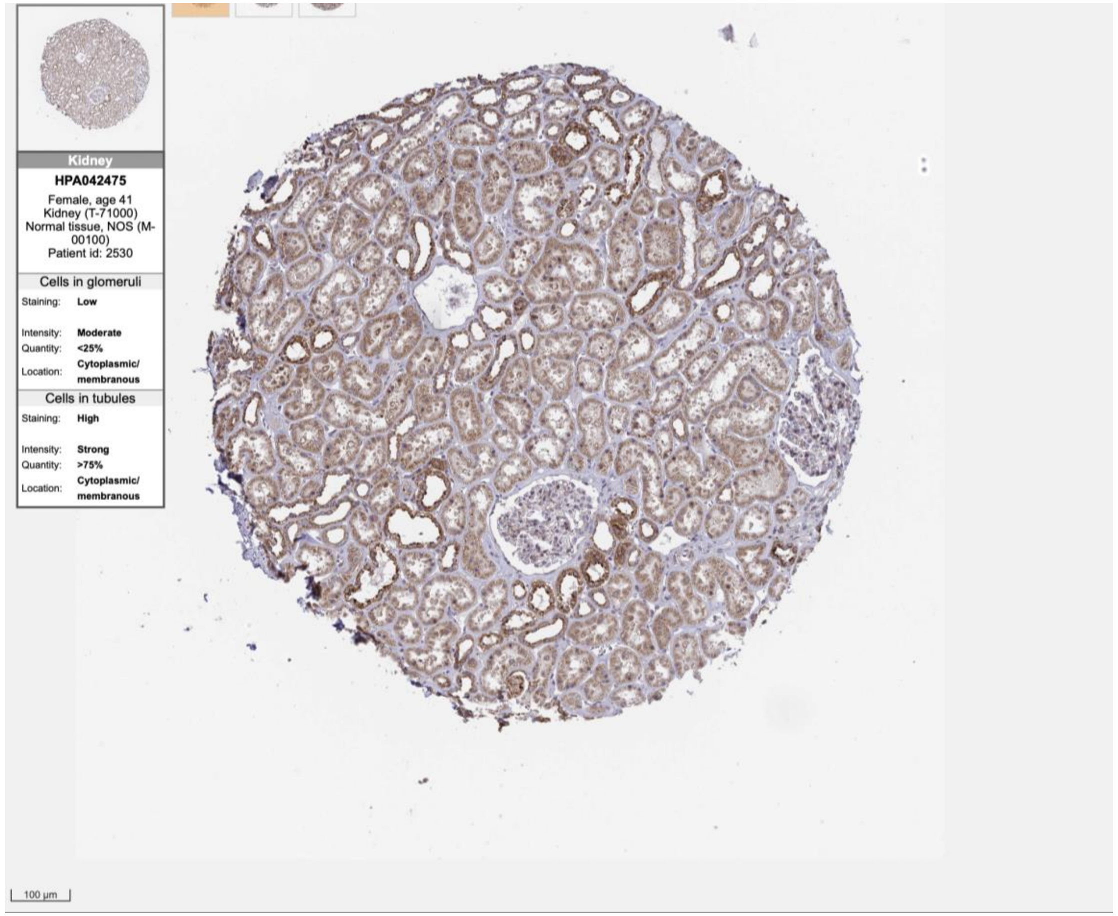
Kidney
HPA042475
Female, age 41
Kidney (T-71000)
Normal tissue, NOS (M-00100)
Patient id: 2530
Cells in glomeruli
Staining: Low
Intensity: Moderate
Quantity: <25%
Location: Cytoplasmic/ membranous
Cells in tubules
Staining: High
Intensity: Strong
Quantity: >75%
Location: Cytoplasmic/ membranous
100 µm

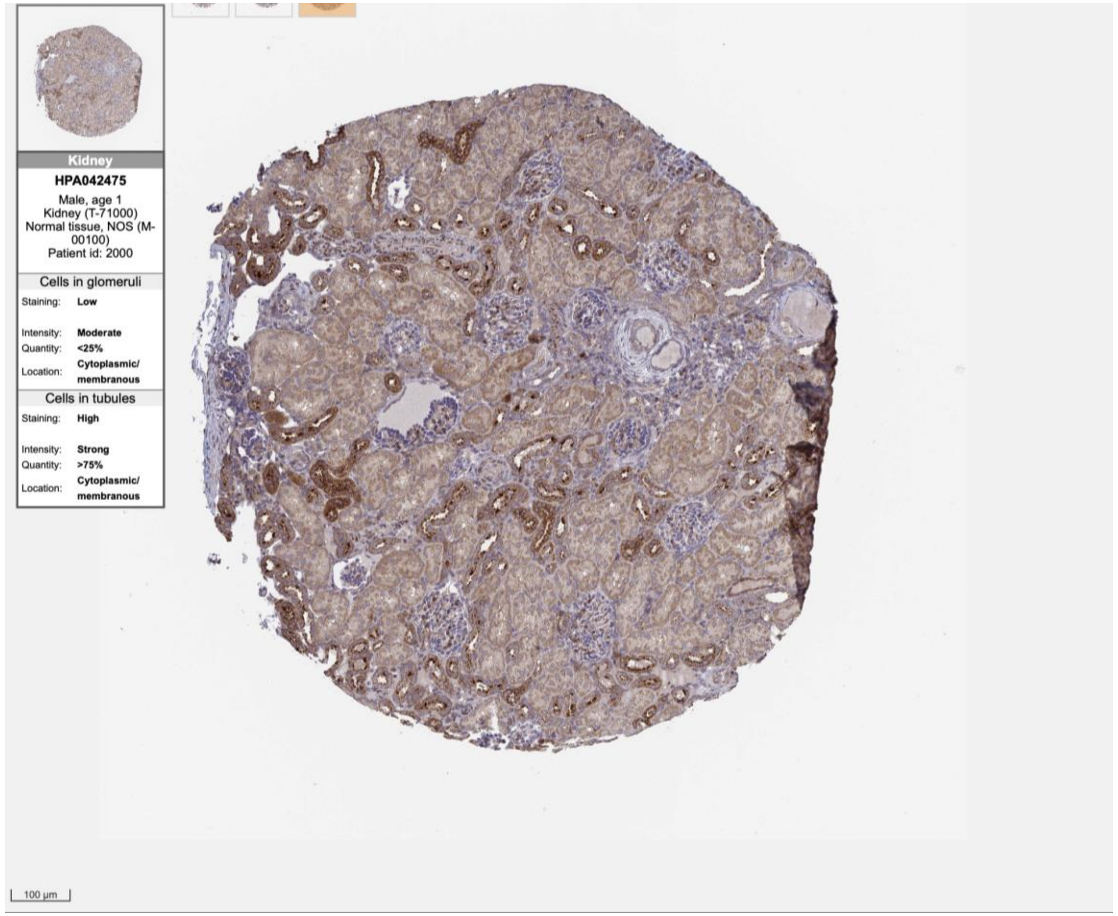
Kidney
HPA042475
Male, age 1
Kidney (T-71000)
Normal tissue, NOS (M-00100)
Patient id: 2000
Cells in glomeruli
Staining: Low
Intensity: Moderate
Quantity: <25%
Location: Cytoplasmic/ membranous
Cells in tubules
Staining: High
Intensity: Strong
Quantity: >75%
Location: Cytoplasmic/ membranous
100 µm

**GATA4**

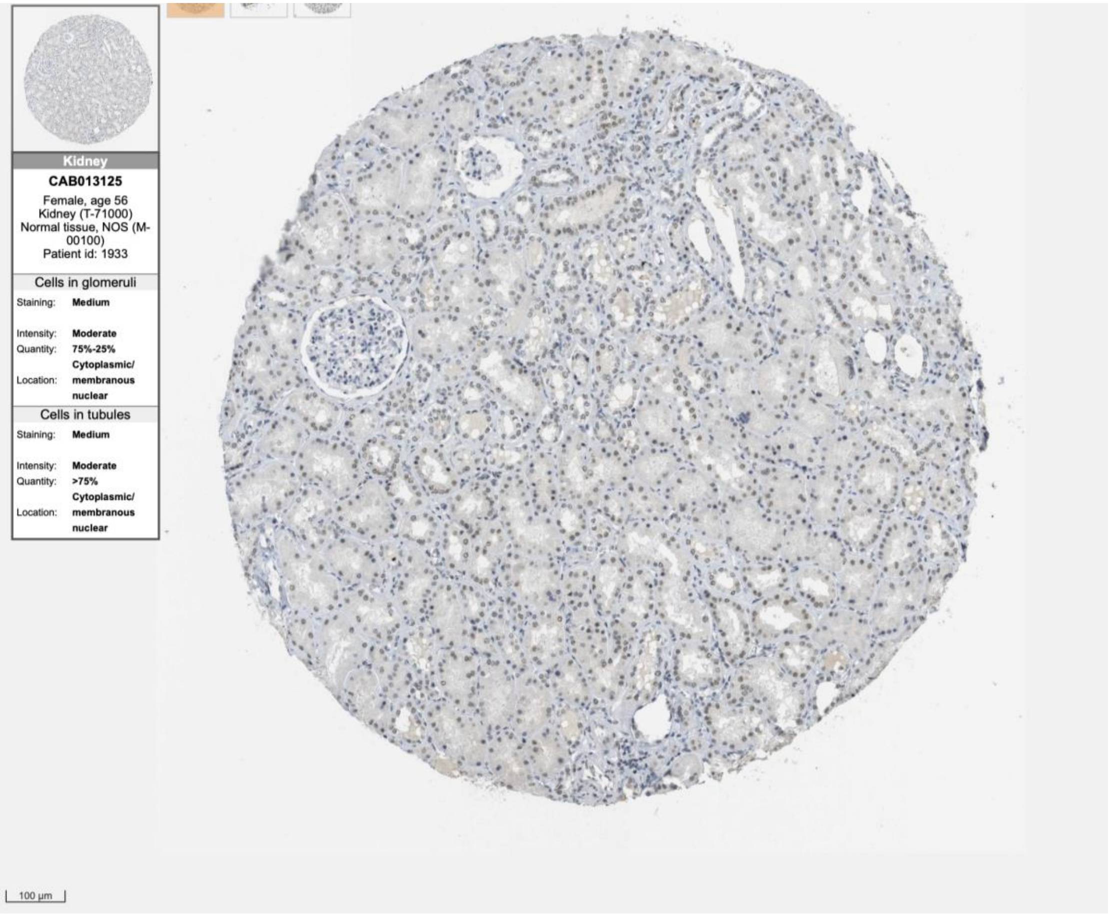

Kidney
CAB013125
Female, age 56
Kidney (T-71000)
Normal tissue, NOS (M-00100)
Patient id: 1933
Cells in glomeruli
Staining: Medium
Intensity: Moderate
Quantity: 75%-25%
Location: Cytoplasmic/ membranous nuclear
Cells in tubules
Staining: Medium
Intensity: Moderate
Quantity: >75%
Location: Cytoplasmic/ membranous nuclear
100 µm


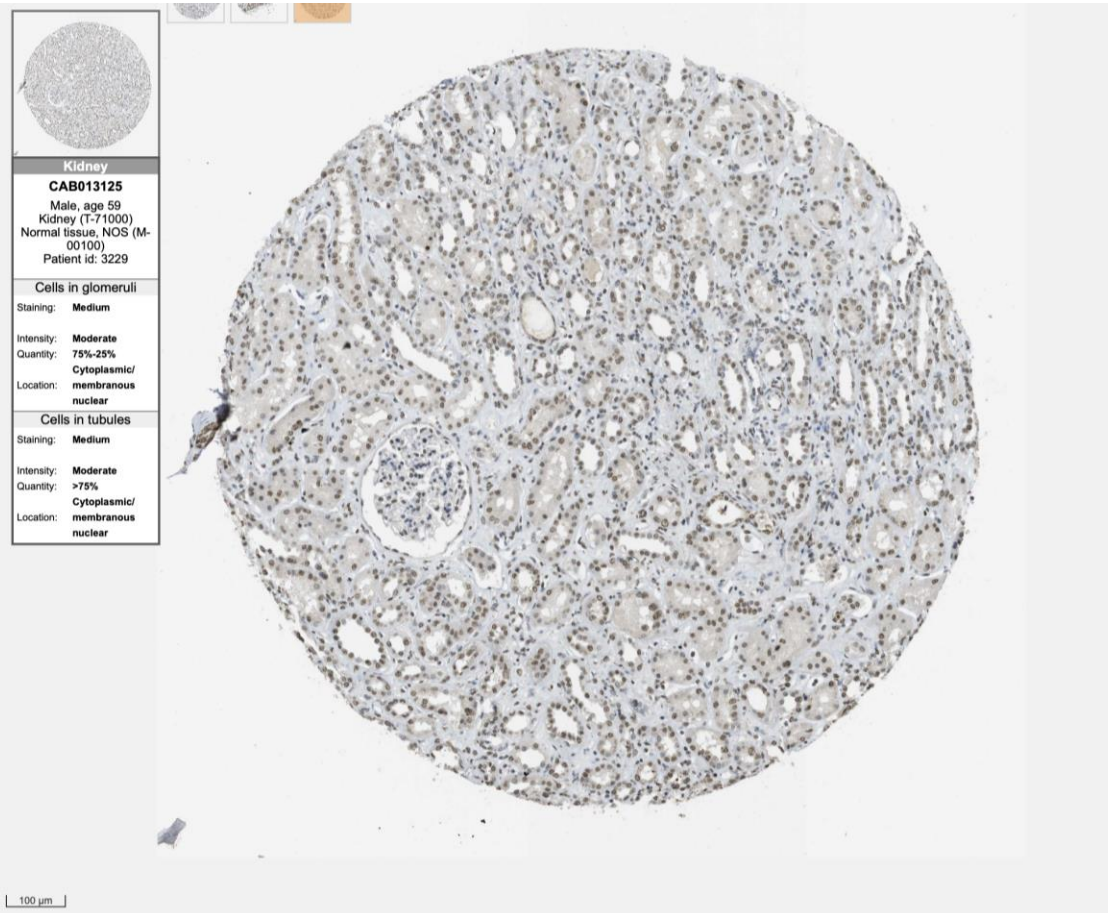

Kidney
CAB013125
Male, age 59
Kidney (T-71000)
Normal tissue, NOS (M-00100)
Patient id: 3229
Cells in glomeruli
Staining: Medium
Intensity: Moderate
Quantity: 75%-25%
Location: Cytoplasmic/ membranous nuclear
Cells in tubules
Staining: Medium
Intensity: Moderate
Quantity: >75%
Location: Cytoplasmic/ membranous nuclear
100 µm

MEIS1

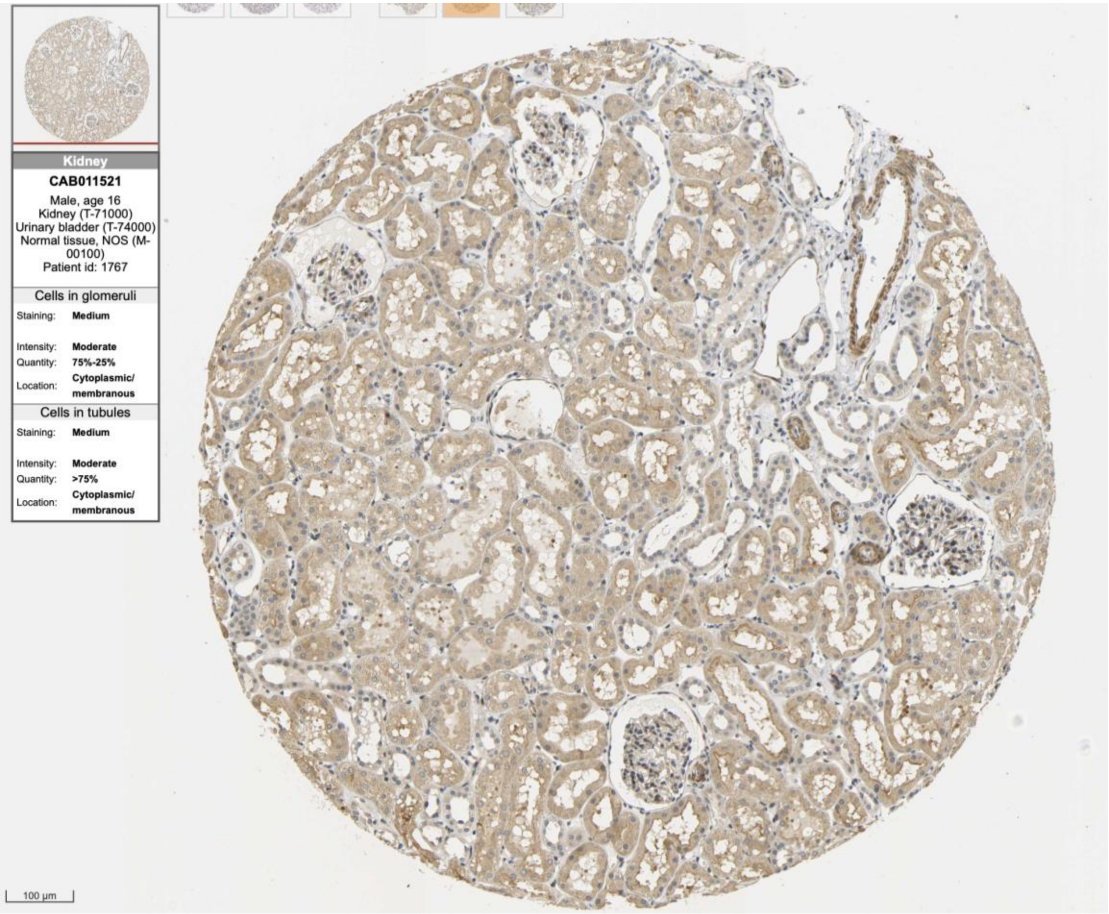
Kidney
CAB011521
Male, age 16
Kidney (T-71000)
Urinary bladder (T-74000)
Normal tissue, NOS (M-00100)
Patient id: 1767
Cells in glomeruli
Staining: Medium
Intensity: Moderate
Quantity: 75%-25%
Location: Cytoplasmic/membranous
Cells in tubules
Staining: Medium
Intensity: Moderate
Quantity: >75%
Location: Cytoplasmic/membranous
100 µm

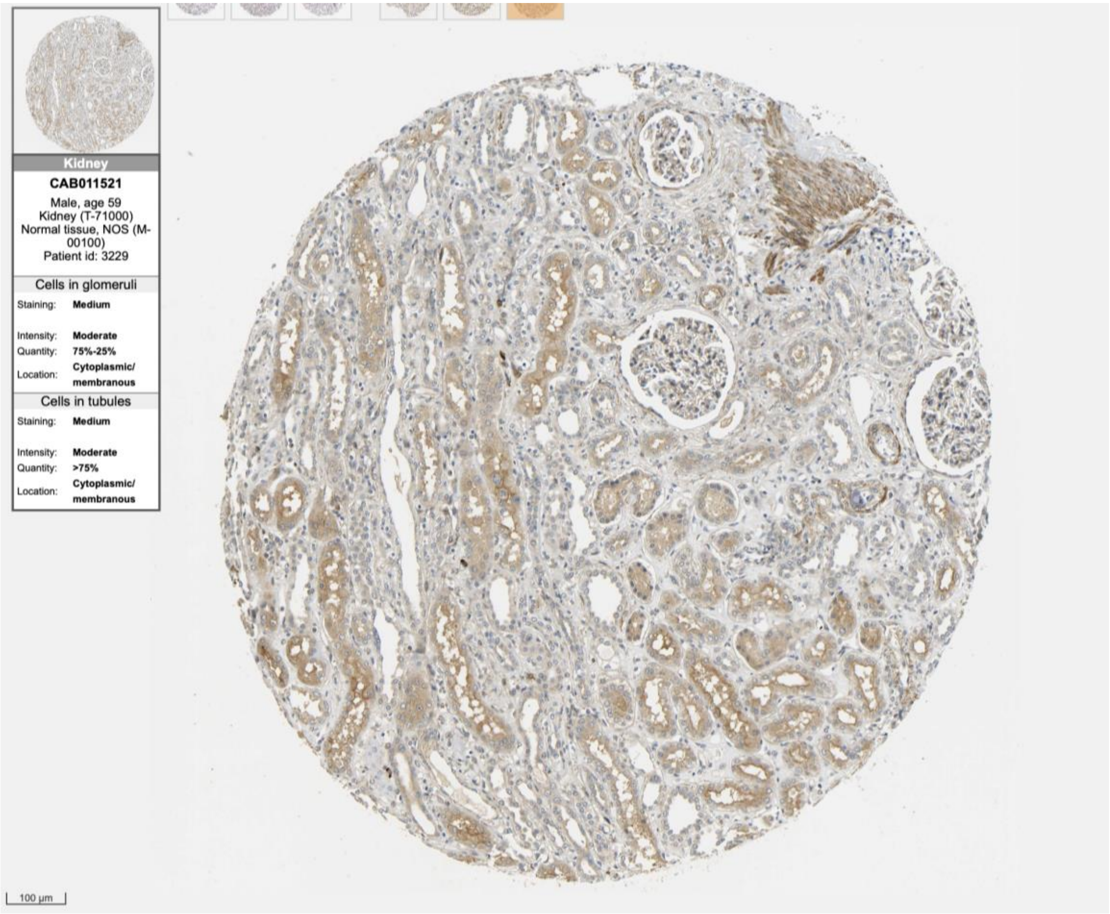
Kidney
CAB011521
Male, age 59
Kidney (T-71000)
Normal tissue, NOS (M-00100)
Patient id: 3229
Cells in glomeruli
Staining: Medium
Intensity: Moderate
Quantity: 75%-25%
Location: Cytoplasmic/membranous
Cells in tubules
Staining: Medium
Intensity: Moderate
Quantity: >75%
Location: Cytoplasmic/membranous
100 µm

SFMBT2

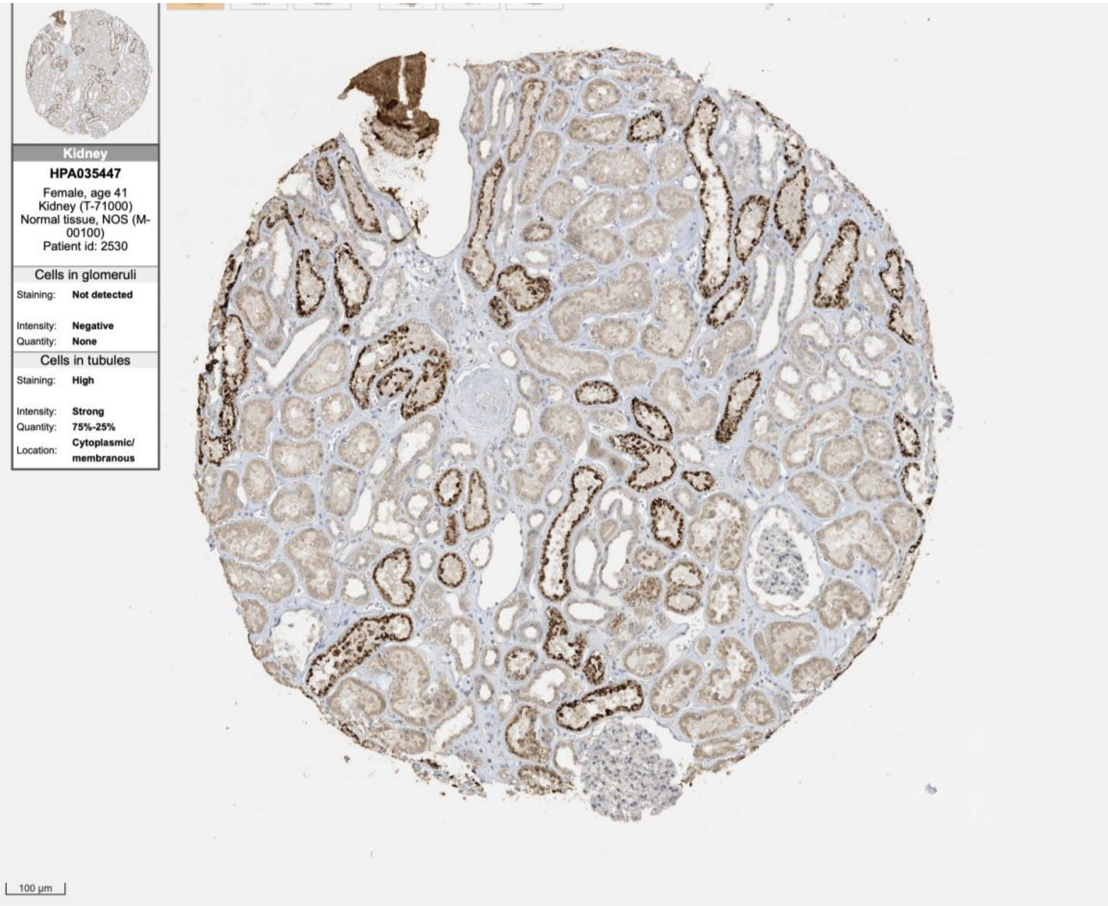

Kidney
HPA035447
Female, age 41
Kidney (T-71000)
Normal tissue, NOS (M-00100)
Patient id: 2530
Cells in glomeruli
Staining: Not detected
Intensity: Negative
Quantity: None
Cells in tubules
Staining: High
Intensity: Strong
Quantity: 75%-25%
Location: Cytoplasmic/ membranous
100 µm


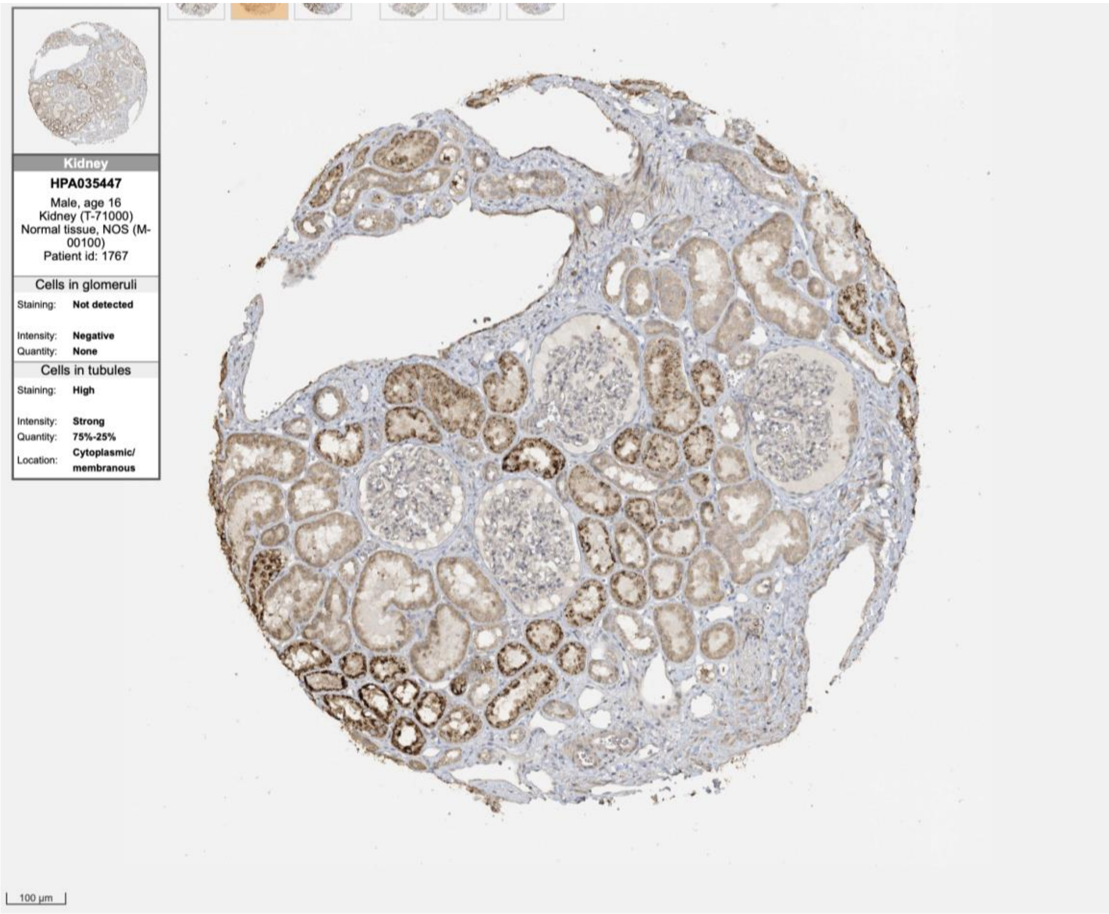

Kidney
HPA035447
Male, age 16
Kidney (T-71000)
Normal tissue, NOS (M-00100)
Patient id: 1767
Cells in glomeruli
Staining: Not detected
Intensity: Negative
Quantity: None
Cells in tubules
Staining: High
Intensity: Strong
Quantity: 75%-25%
Location: Cytoplasmic/ membranous
100 µm

**PSMC3IP**

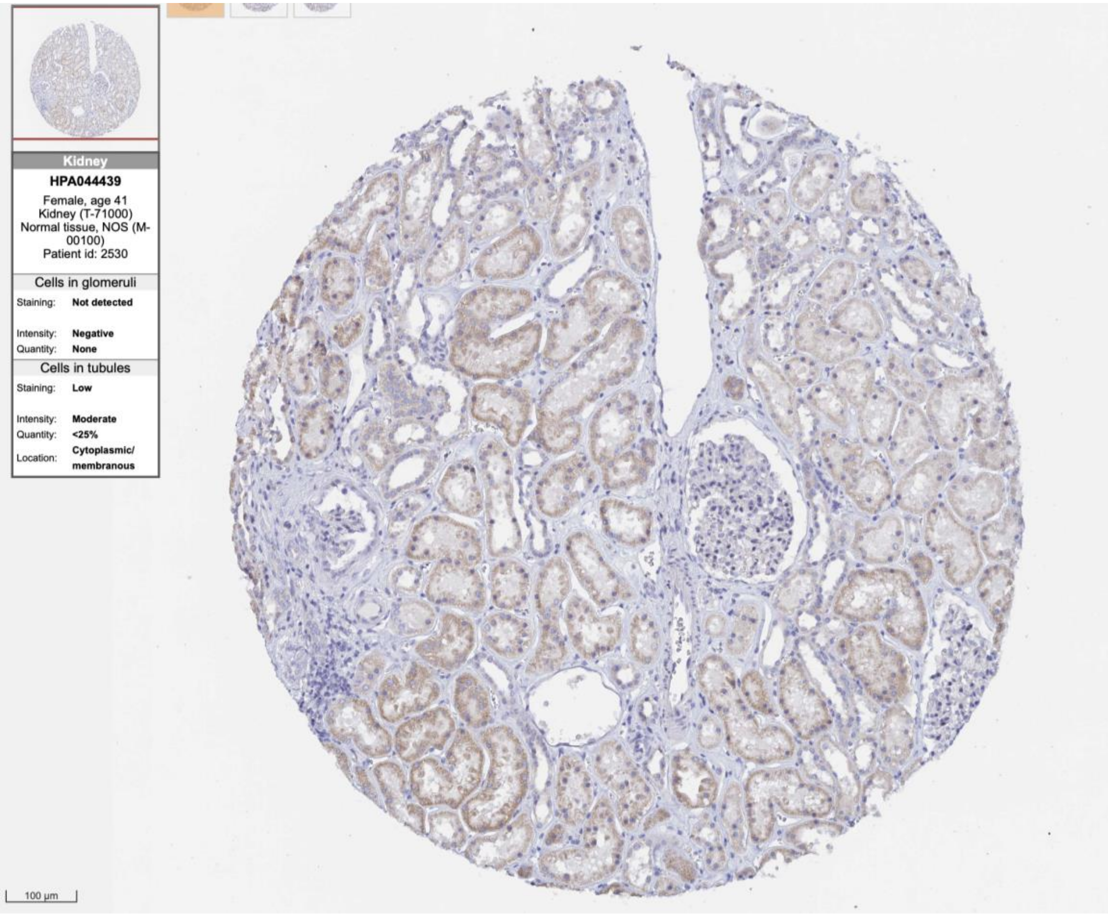
Kidney
HPA044439
Female, age 41
Kidney (T-71000)
Normal tissue, NOS (M-00100)
Patient id: 2530
Cells in glomeruli
Staining: Not detected
Intensity: Negative
Quantity: None
Cells in tubules
Staining: Low
Intensity: Moderate
Quantity: <25%
Location: Cytoplasmic/ membranous
100 µm

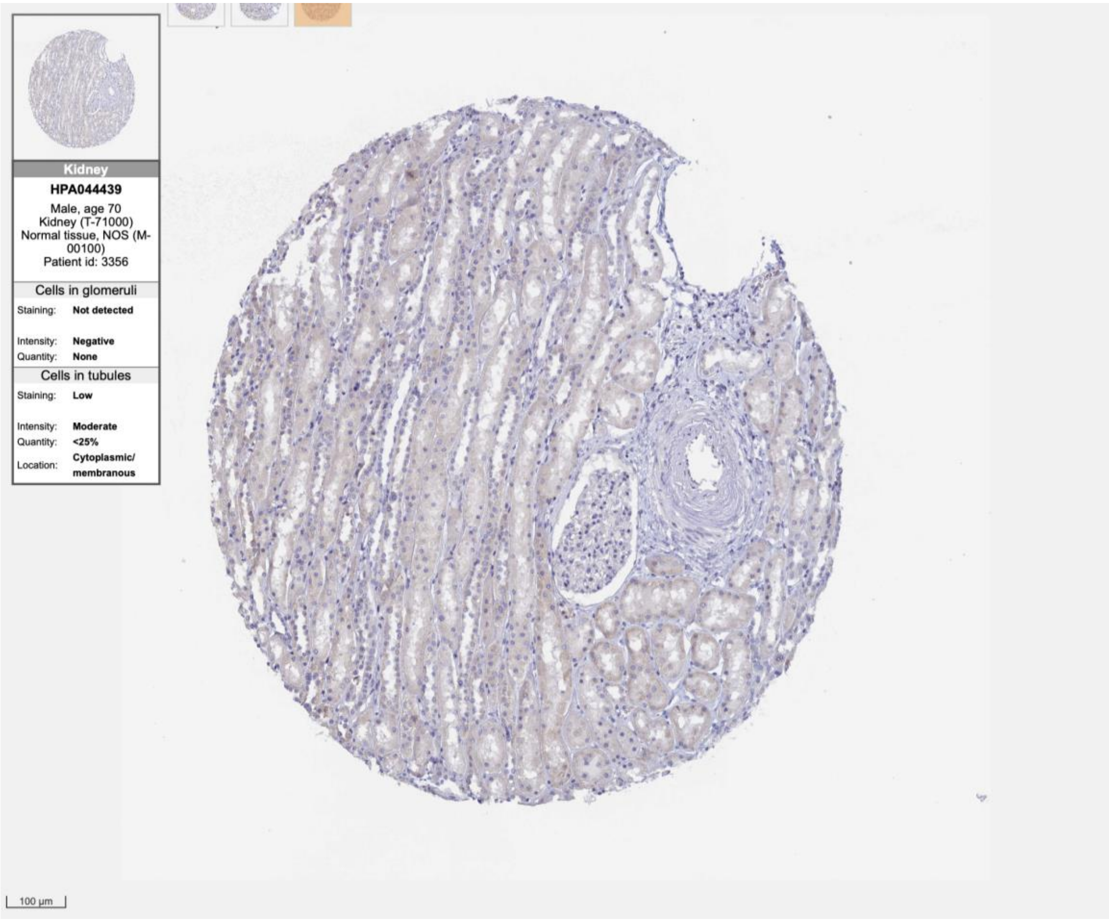
Kidney
HPA044439
Male, age 70
Kidney (T-71000)
Normal tissue, NOS (M-00100)
Patient id: 3356
Cells in glomeruli
Staining: Not detected
Intensity: Negative
Quantity: None
Cells in tubules
Staining: Low
Intensity: Moderate
Quantity: <25%
Location: Cytoplasmic/ membranous
100 µm

TFAP2B

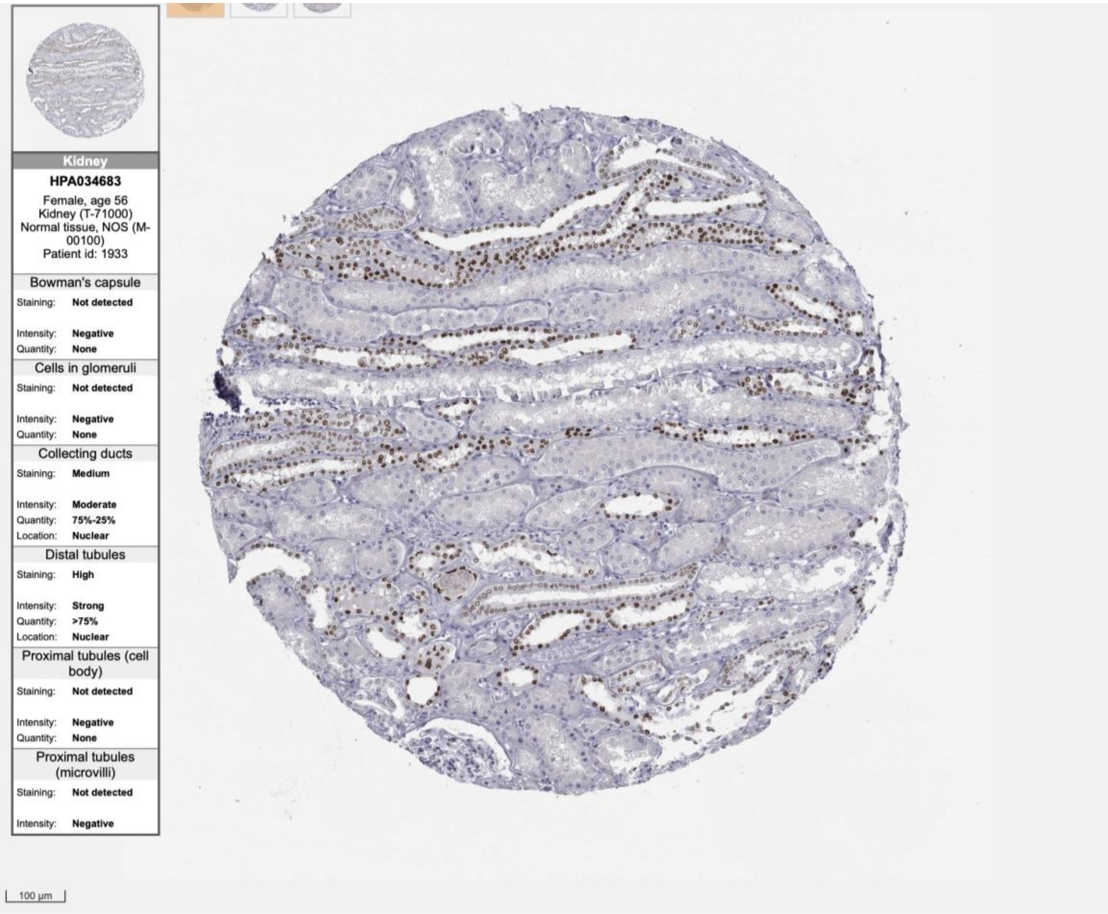
Kidney
HPA034683
Female, age 56
Kidney (T-71000)
Normal tissue, NOS (M-00100)
Patient id: 1933
Bowman's capsule
Staining: Not detected
Intensity: Negative
Quantity: None
Cells in glomeruli
Staining: Not detected
Intensity: Negative
Quantity: None
Collecting ducts
Staining: Medium
Intensity: Moderate
Quantity: 75%-25%
Location: Nuclear
Distal tubules
Staining: High
Intensity: Strong
Quantity: >75%
Location: Nuclear
Proximal tubules (cell body)
Staining: Not detected
Intensity: Negative
Quantity: None
Proximal tubules (microvilli)
Staining: Not detected
Intensity: Negative
100 µm

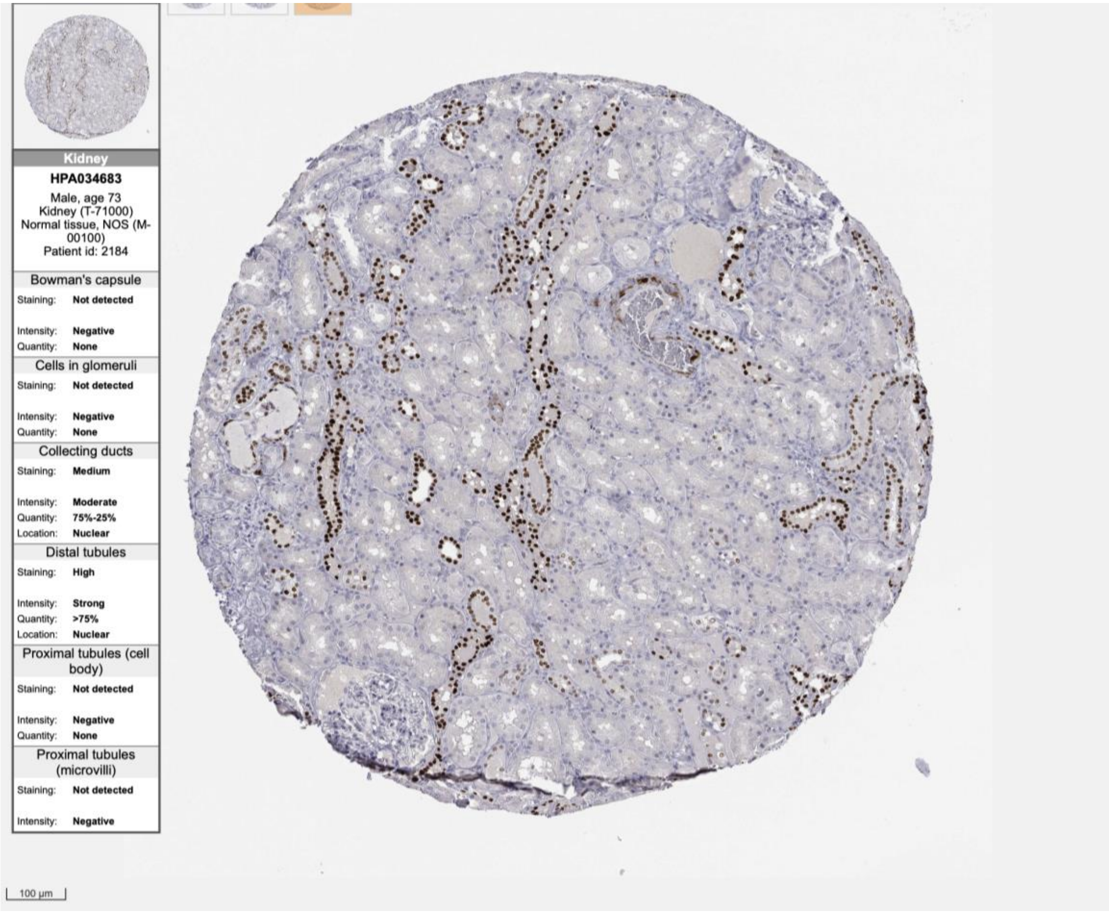
Kidney
HPA034683
Male, age 73
Kidney (T-71000)
Normal tissue, NOS (M-00100)
Patient id: 2184
Bowman's capsule
Staining: Not detected
Intensity: Negative
Quantity: None
Cells in glomeruli
Staining: Not detected
Intensity: Negative
Quantity: None
Collecting ducts
Staining: Medium
Intensity: Moderate
Quantity: 75%-25%
Location: Nuclear
Distal tubules
Staining: High
Intensity: Strong
Quantity: >75%
Location: Nuclear
Proximal tubules (cell body)
Staining: Not detected
Intensity: Negative
Quantity: None
Proximal tubules (microvilli)
Staining: Not detected
Intensity: Negative
100 µm

ZNF343

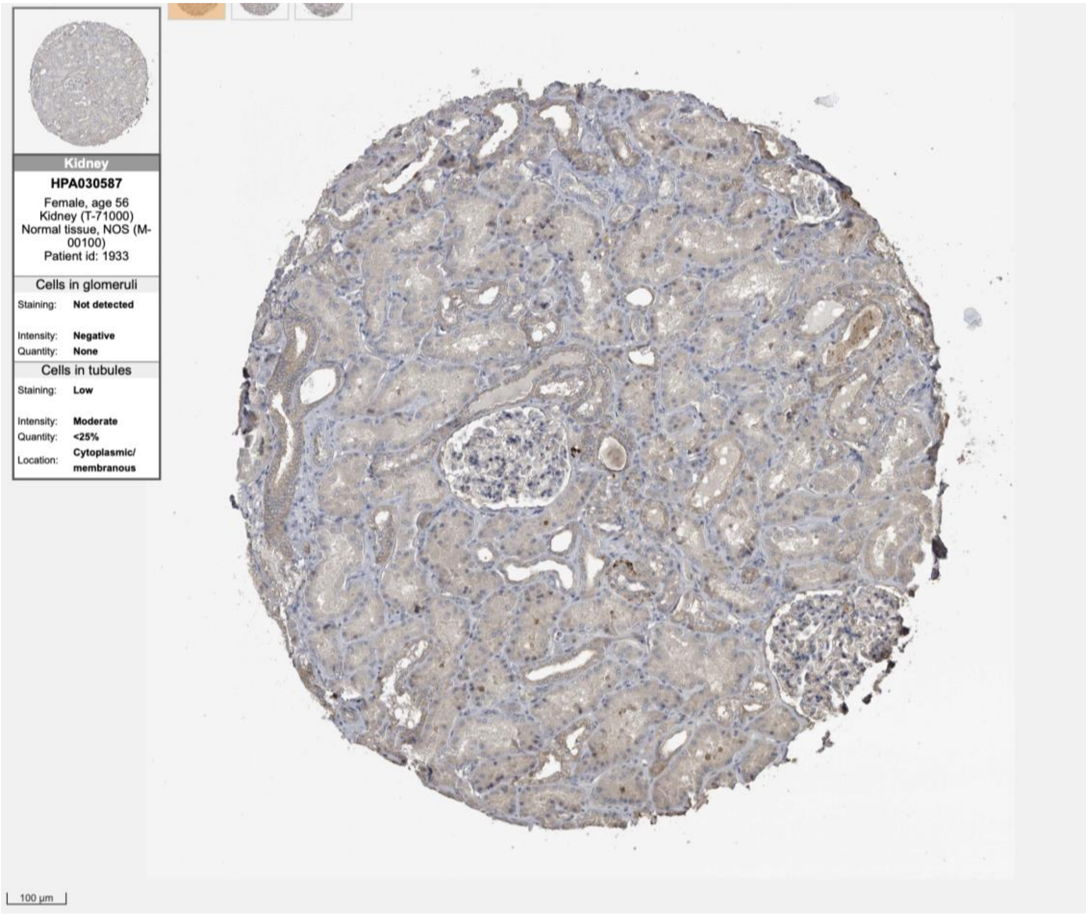

Kidney
HPA030587
Female, age 56
Kidney (T-71000)
Normal tissue, NOS (M-00100)
Patient id: 1933
Cells in glomeruli
Staining: Not detected
Intensity: Negative
Quantity: None
Cells in tubules
Staining: Low
Intensity: Moderate
Quantity: <25%
Location: Cytoplasmic/membranous
100 µm


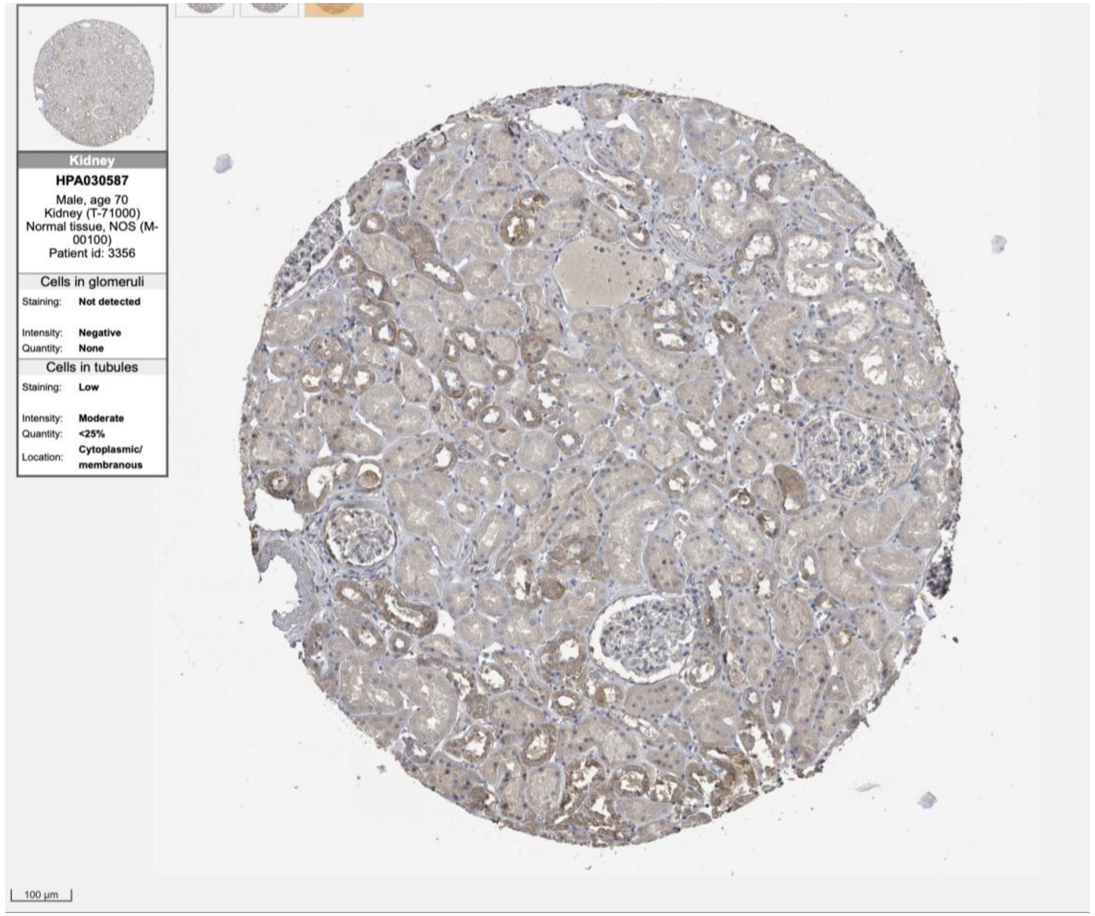

Kidney
HPA030587
Male, age 70
Kidney (T-71000)
Normal tissue, NOS (M-00100)
Patient id: 3356
Cells in glomeruli
Staining: Not detected
Intensity: Negative
Quantity: None
Cells in tubules
Staining: Low
Intensity: Moderate
Quantity: <25%
Location: Cytoplasmic/membranous
100 µm

ZNF367

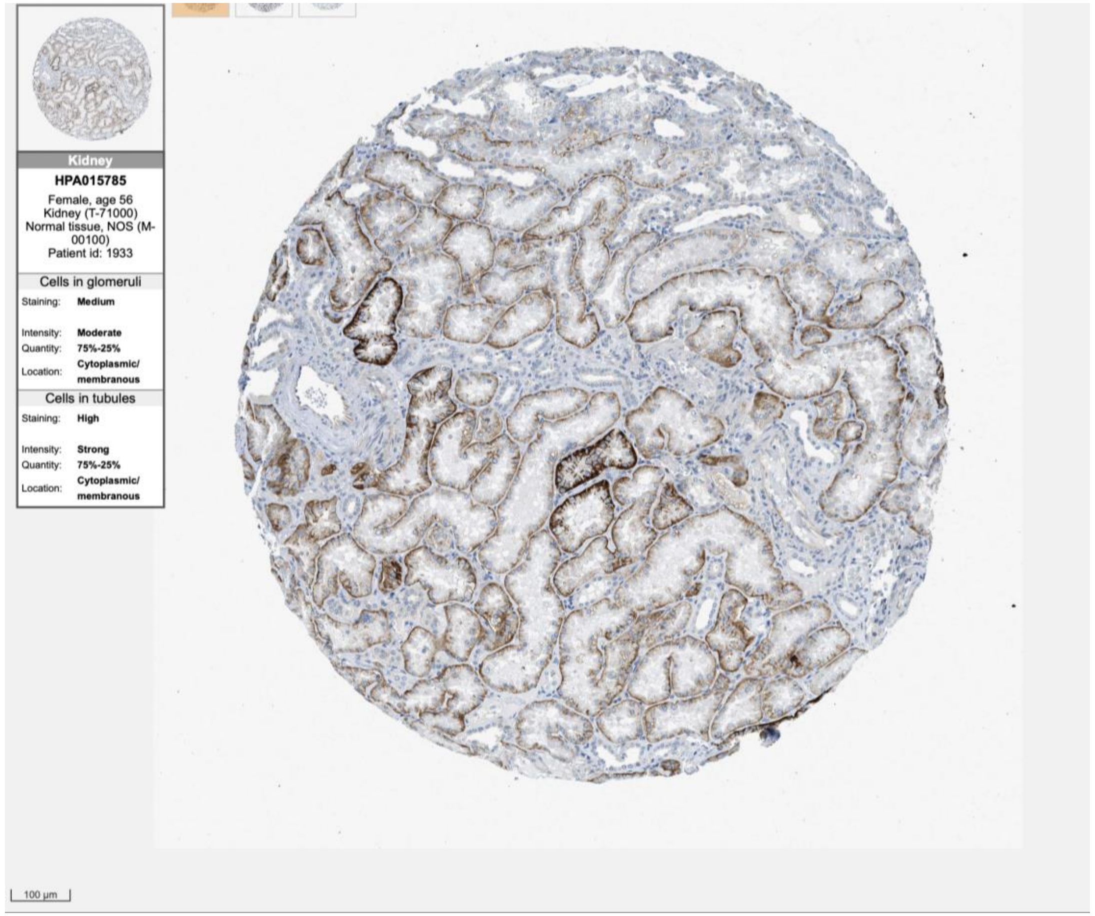

Kidney
HPA015785
Female, age 56
Kidney (T-71000)
Normal tissue, NOS (M-00100)
Patient id: 1933
Cells in glomeruli
Staining: Medium
Intensity: Moderate
Quantity: 75%-25%
Location: Cytoplasmic/membranous
Cells in tubules
Staining: High
Intensity: Strong
Quantity: 75%-25%
Location: Cytoplasmic/membranous
100 µm


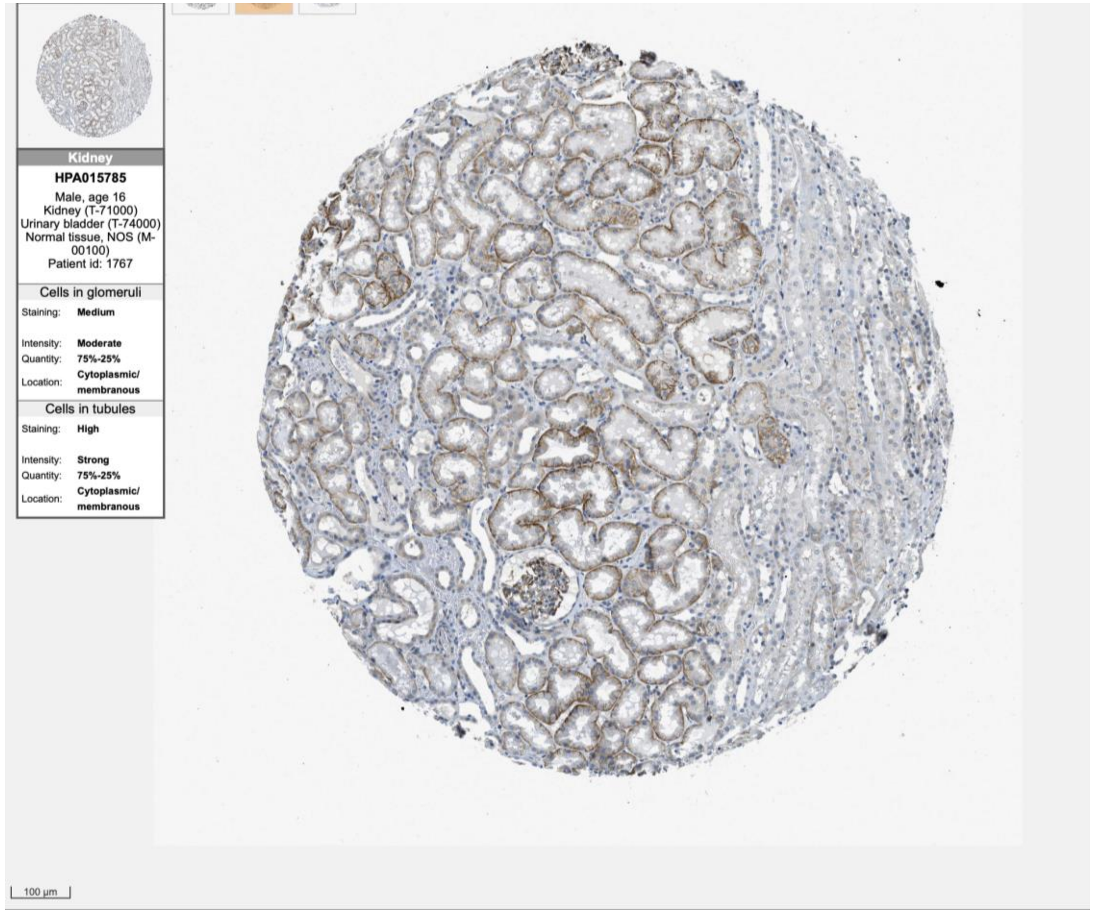

Kidney
HPA015785
Male, age 16
Kidney (T-71000)
Urinary bladder (T-74000)
Normal tissue, NOS (M-00100)
Patient id: 1767
Cells in glomeruli
Staining: Medium
Intensity: Moderate
Quantity: 75%-25%
Location: Cytoplasmic/membranous
Cells in tubules
Staining: High
Intensity: Strong
Quantity: 75%-25%
Location: Cytoplasmic/membranous
100 µm

ZNF529

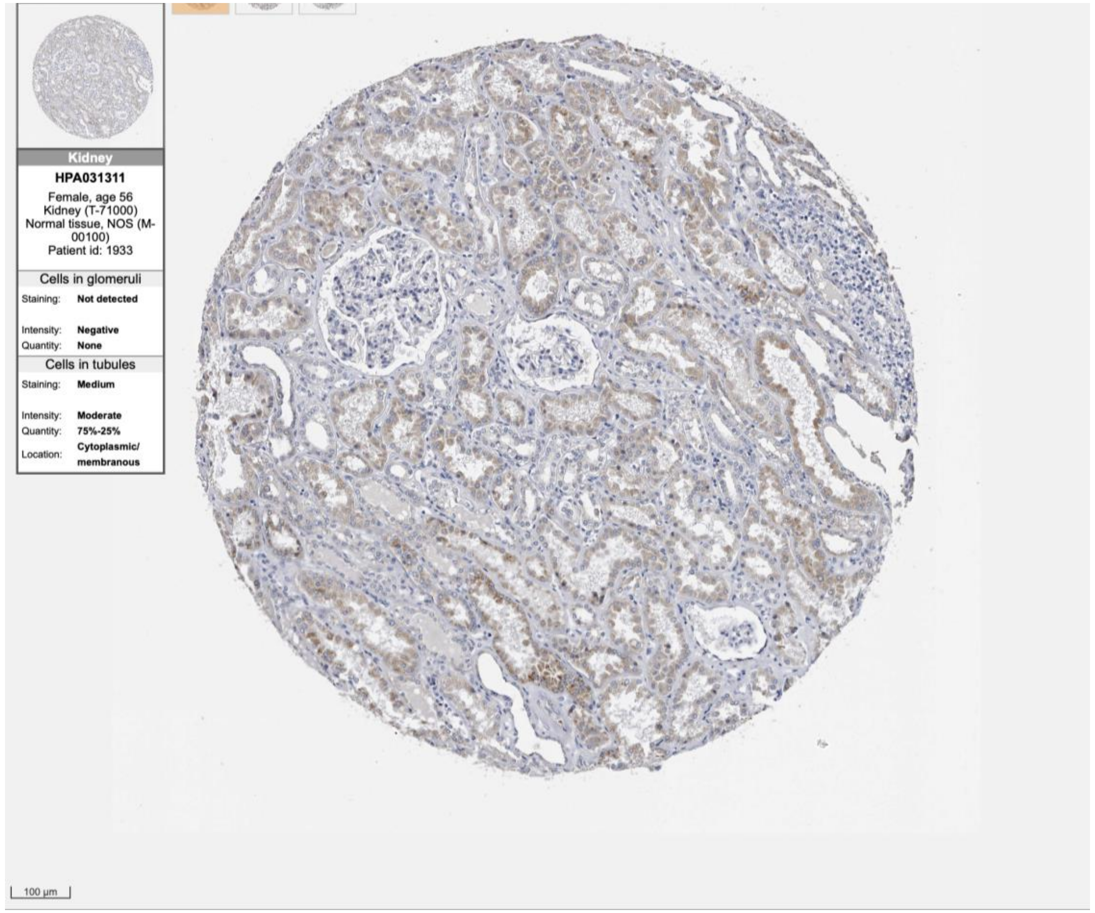
Kidney
HPA031311
Female, age 56
Kidney (T-71000)
Normal tissue, NOS (M-00100)
Patient id: 1933
Cells in glomeruli
Staining: Not detected
Intensity: Negative
Quantity: None
Cells in tubules
Staining: Medium
Intensity: Moderate
Quantity: 75%-25%
Location: Cytoplasmic/ membranous
100 µm

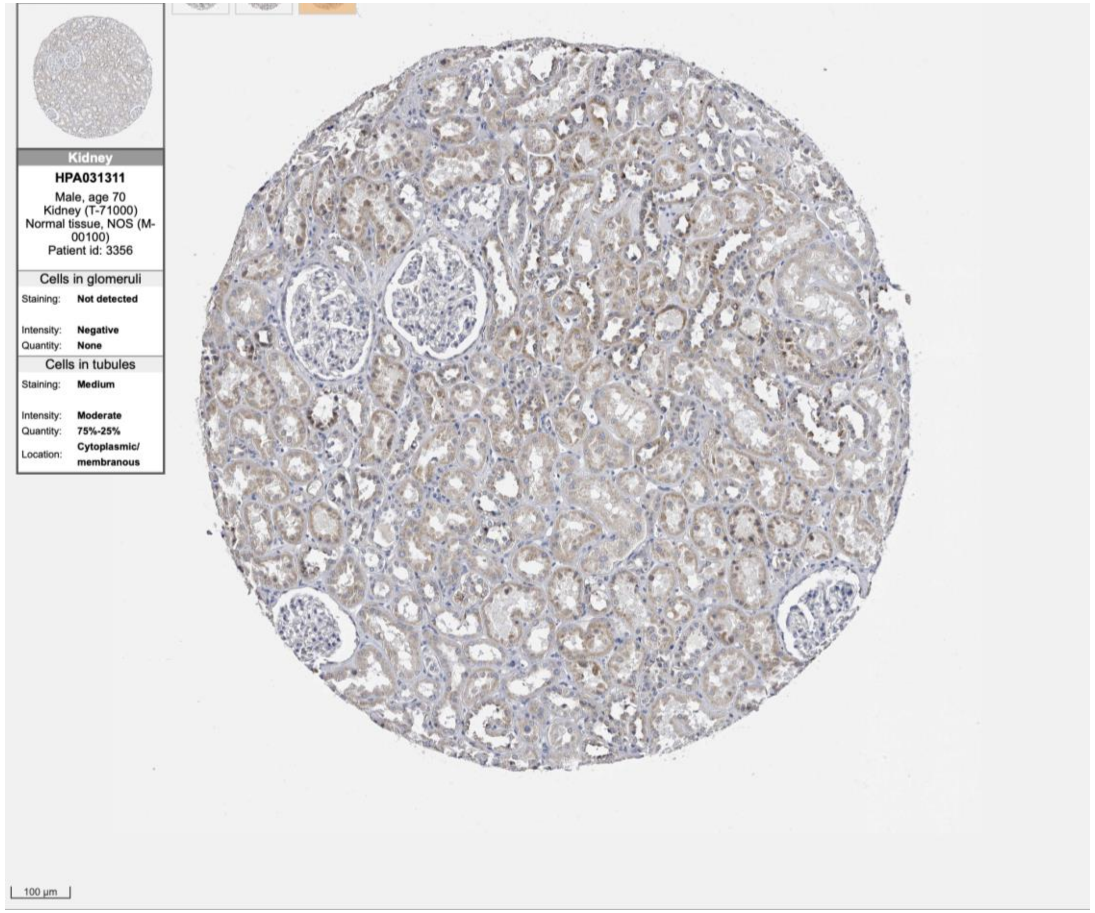
Kidney
HPA031311
Male, age 70
Kidney (T-71000)
Normal tissue, NOS (M-00100)
Patient id: 3356
Cells in glomeruli
Staining: Not detected
Intensity: Negative
Quantity: None
Cells in tubules
Staining: Medium
Intensity: Moderate
Quantity: 75%-25%
Location: Cytoplasmic/ membranous
100 µm

**ZNF665**

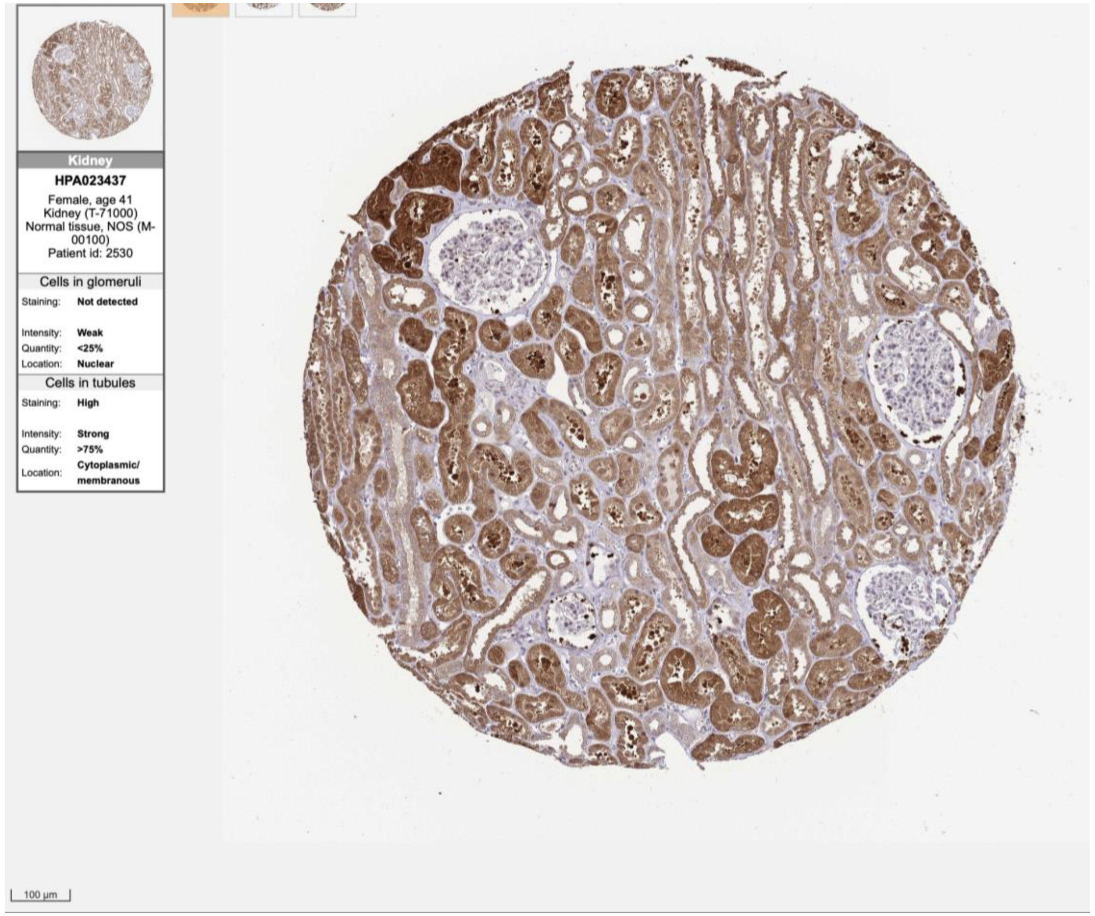

Kidney
HPA023437
Female, age 41
Kidney (T-71000)
Normal tissue, NOS (M-00100)
Patient id: 2530
Cells in glomeruli
Staining: Not detected
Intensity: Weak
Quantity: <25%
Location: Nuclear
Cells in tubules
Staining: High
Intensity: Strong
Quantity: >75%
Location: Cytoplasmic/ membranous
100 µm


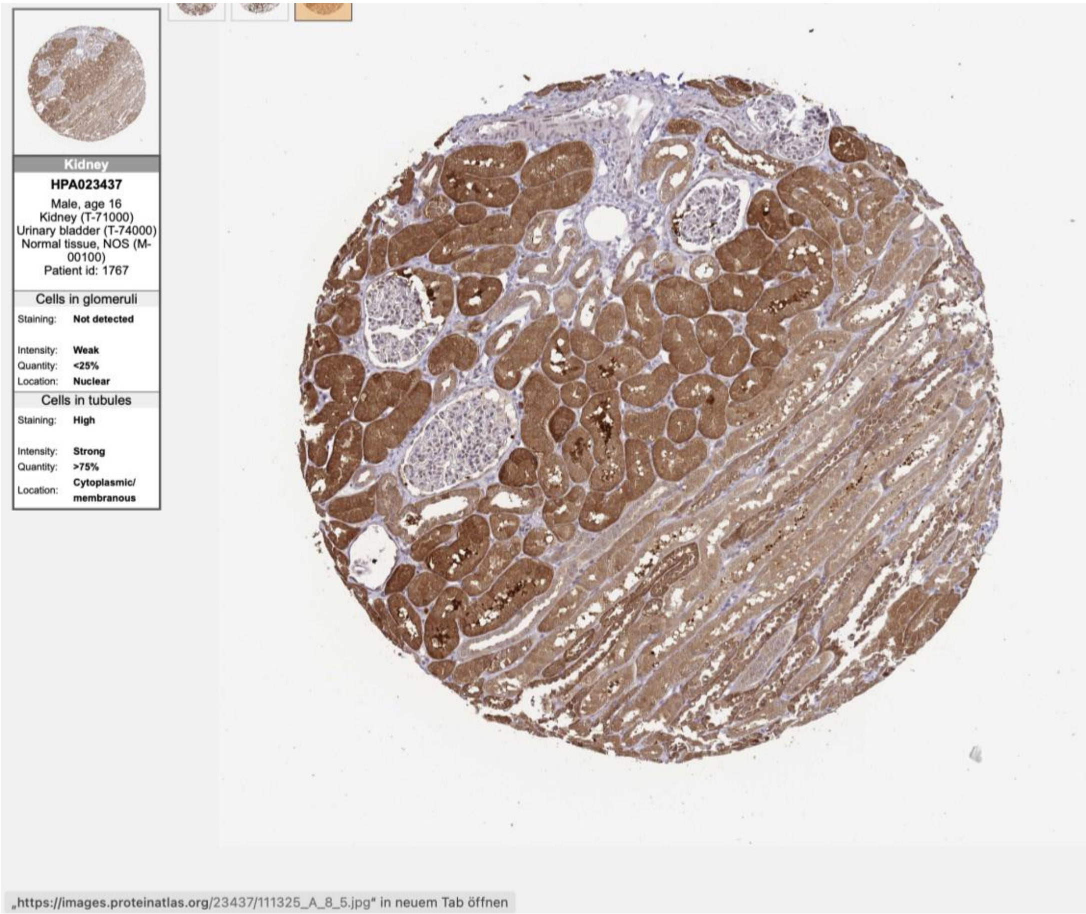

Kidney
HPA023437
Male, age 16
Kidney (T-71000)
Urinary bladder (T-74000)
Normal tissue, NOS (M-00100)
Patient id: 1767
Cells in glomeruli
Staining: Not detected
Intensity: Weak
Quantity: <25%
Location: Nuclear
Cells in tubules
Staining: High
Intensity: Strong
Quantity: >75%
Location: Cytoplasmic/ membranous
„https://images.proteinatlas.org/23437/111325_A_8_5.jpg“ in neuem Tab öffnen

**ZNF669**

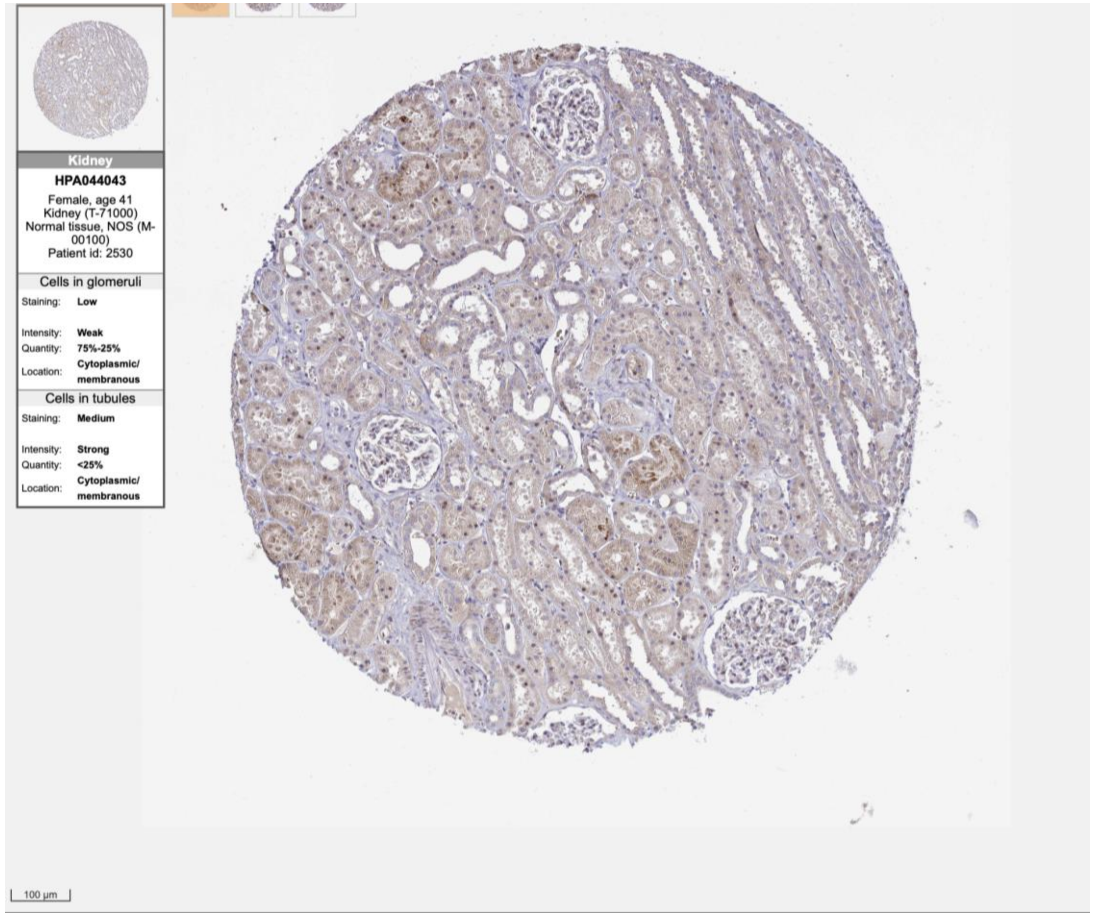
Kidney
HPA044043
Female, age 41
Kidney (T-71000)
Normal tissue, NOS (M-00100)
Patient id: 2530
Cells in glomeruli
Staining: Low
Intensity: Weak
Quantity: 75%-25%
Location: Cytoplasmic/ membranous
Cells in tubules
Staining: Medium
Intensity: Strong
Quantity: <25%
Location: Cytoplasmic/ membranous
100 µm

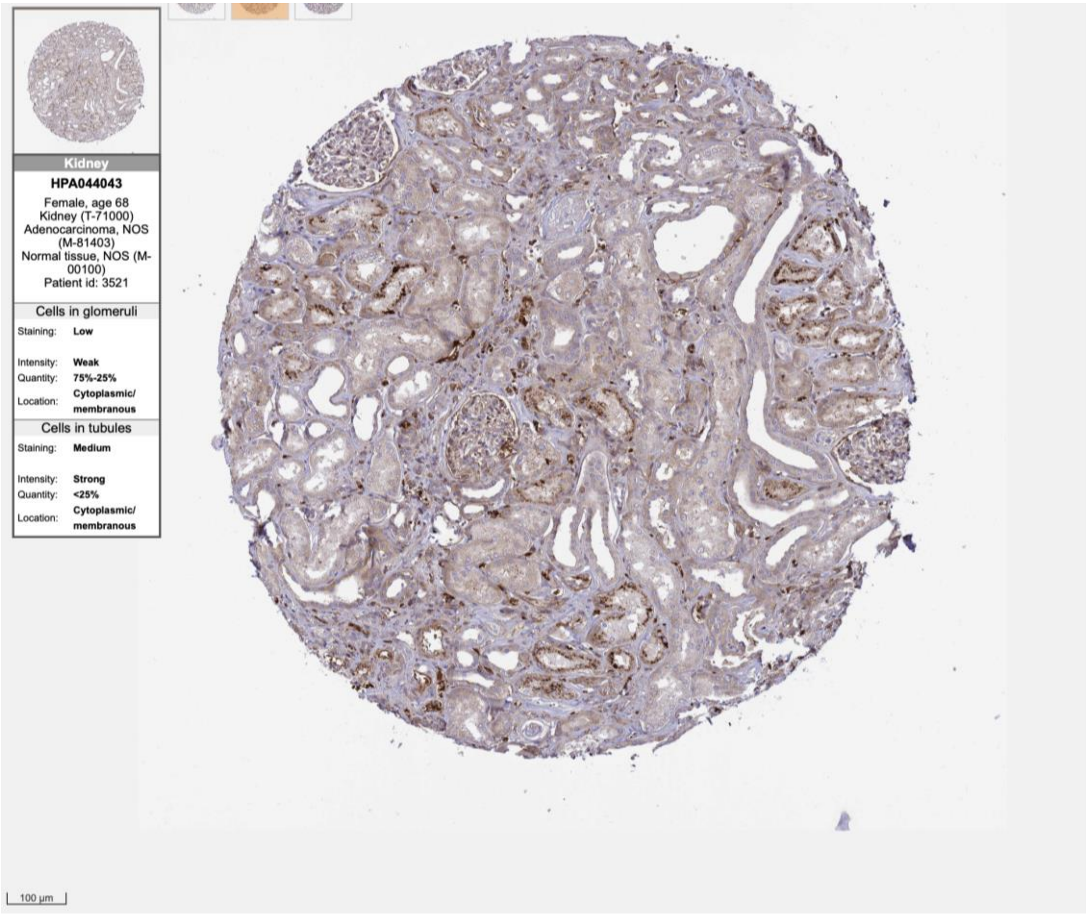
Kidney
HPA044043
Female, age 68
Kidney (T-71000)
Adenocarcinoma, NOS (M-81403)
Normal tissue, NOS (M-00100)
Patient id: 3521
Cells in glomeruli
Staining: Low
Intensity: Weak
Quantity: 75%-25%
Location: Cytoplasmic/ membranous
Cells in tubules
Staining: Medium
Intensity: Strong
Quantity: <25%
Location: Cytoplasmic/ membranous
100 µm

ZNF682

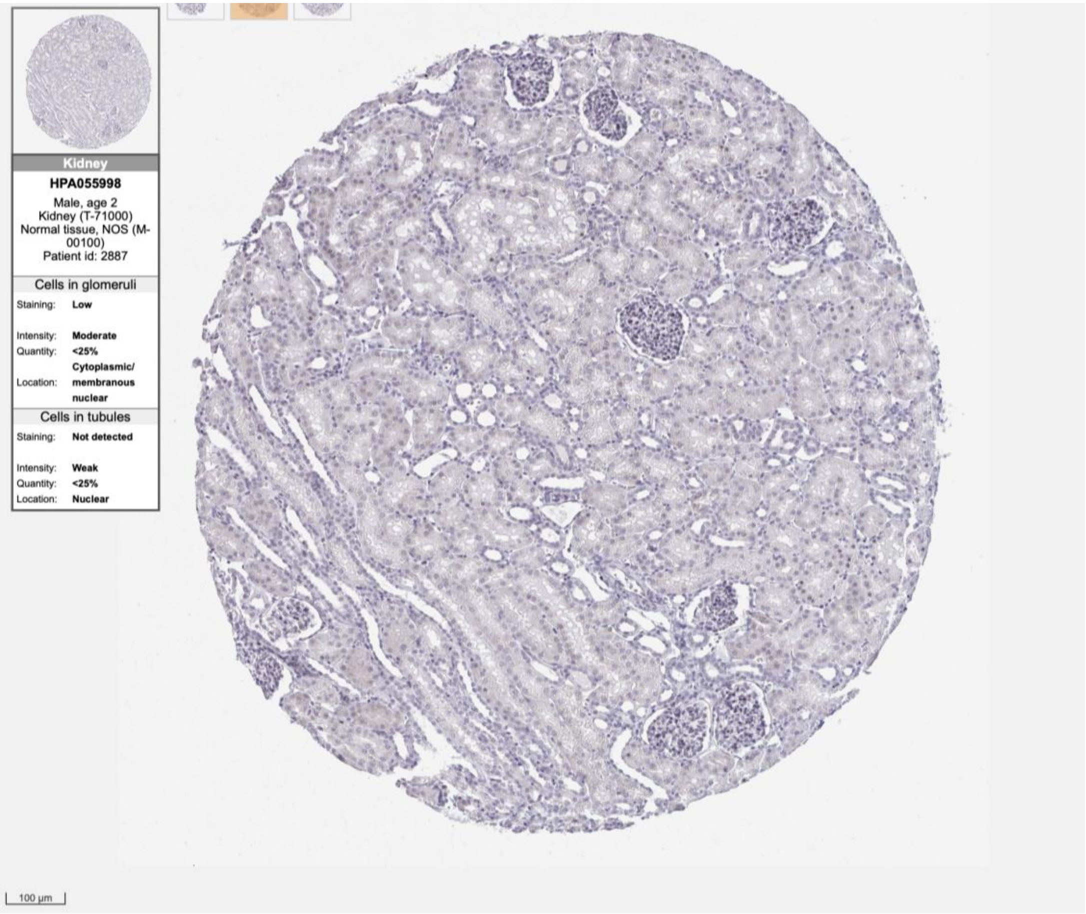
Kidney
HPA055998
Male, age 2
Kidney (T-71000)
Normal tissue, NOS (M-00100)
Patient id: 2887
Cells in glomeruli
Staining: Low
Intensity: Moderate
Quantity: <25%
Location: Cytoplasmic/ membranous nuclear
Cells in tubules
Staining: Not detected
Intensity: Weak
Quantity: <25%
Location: Nuclear
100 µm

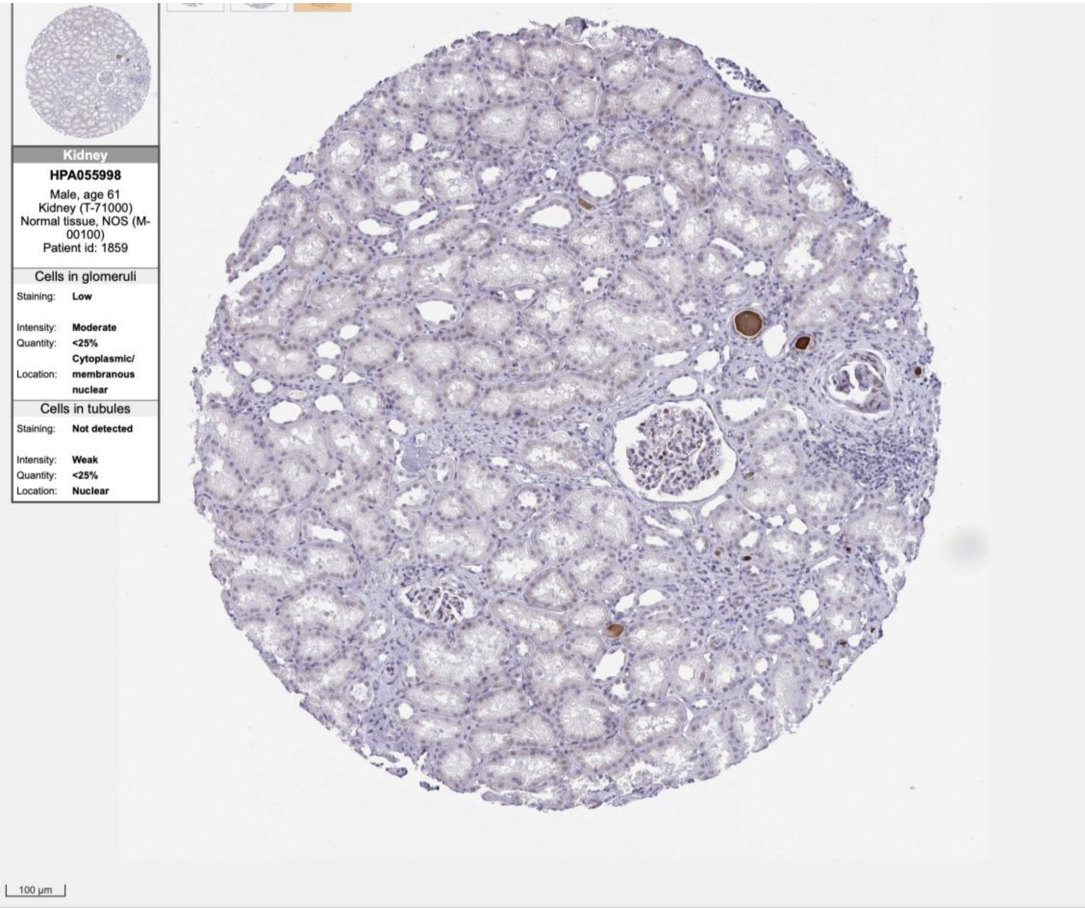
Kidney
HPA055998
Male, age 61
Kidney (T-71000)
Normal tissue, NOS (M-00100)
Patient id: 1859
Cells in glomeruli
Staining: Low
Intensity: Moderate
Quantity: <25%
Location: Cytoplasmic/ membranous nuclear
Cells in tubules
Staining: Not detected
Intensity: Weak
Quantity: <25%
Location: Nuclear
100 µm

ZNF772

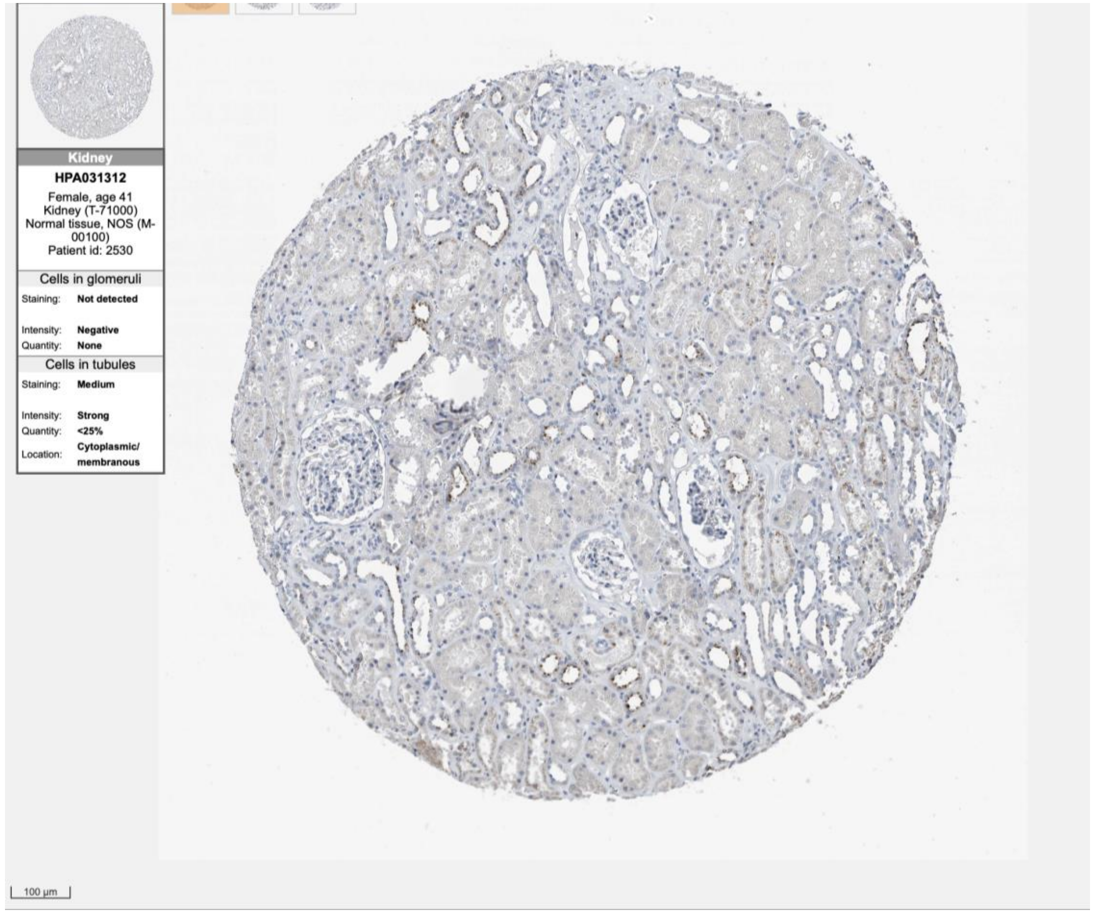
Kidney
HPA031312
Female, age 41
Kidney (T-71000)
Normal tissue, NOS (M-00100)
Patient id: 2530
Cells in glomeruli
Staining: Not detected
Intensity: Negative
Quantity: None
Cells in tubules
Staining: Medium
Intensity: Strong
Quantity: <25%
Location: Cytoplasmic/membranous
100 µm

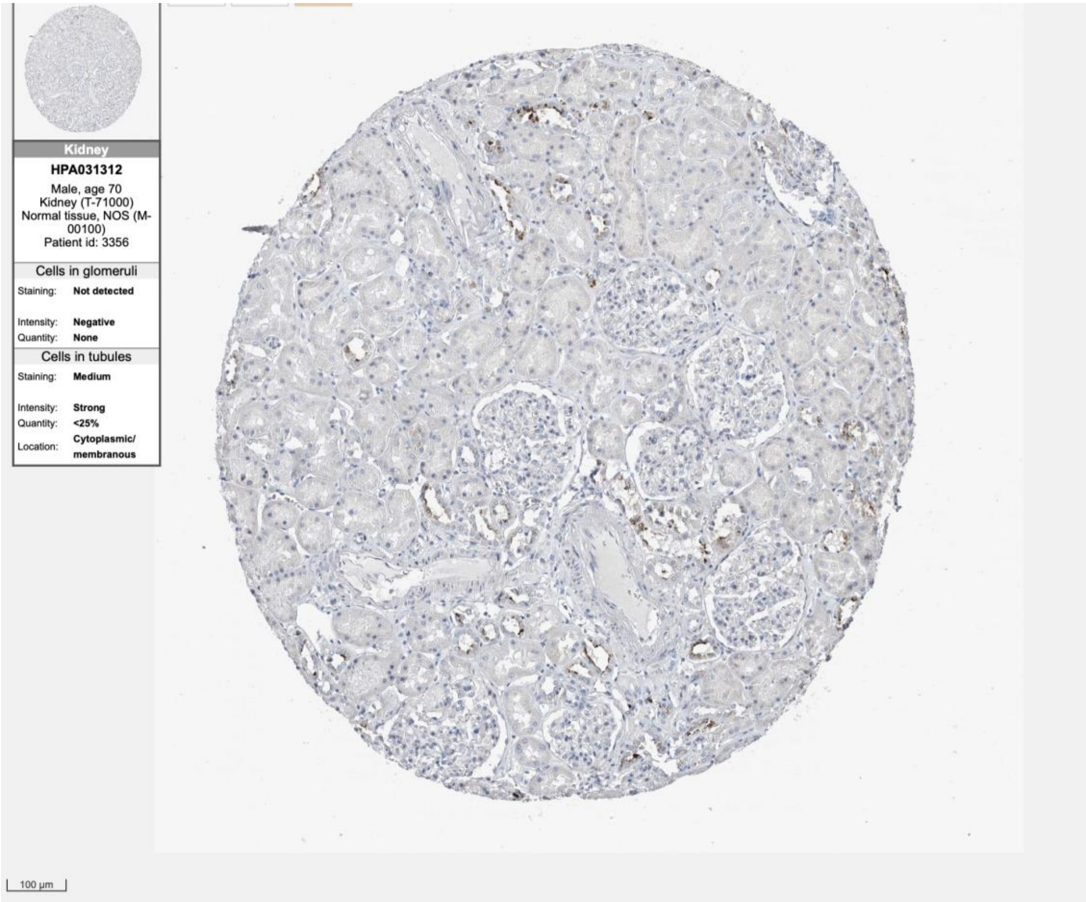
Kidney
HPA031312
Male, age 70
Kidney (T-71000)
Normal tissue, NOS (M-00100)
Patient id: 3356
Cells in glomeruli
Staining: Not detected
Intensity: Negative
Quantity: None
Cells in tubules
Staining: Medium
Intensity: Strong
Quantity: <25%
Location: Cytoplasmic/membranous
100 µm

SOX2

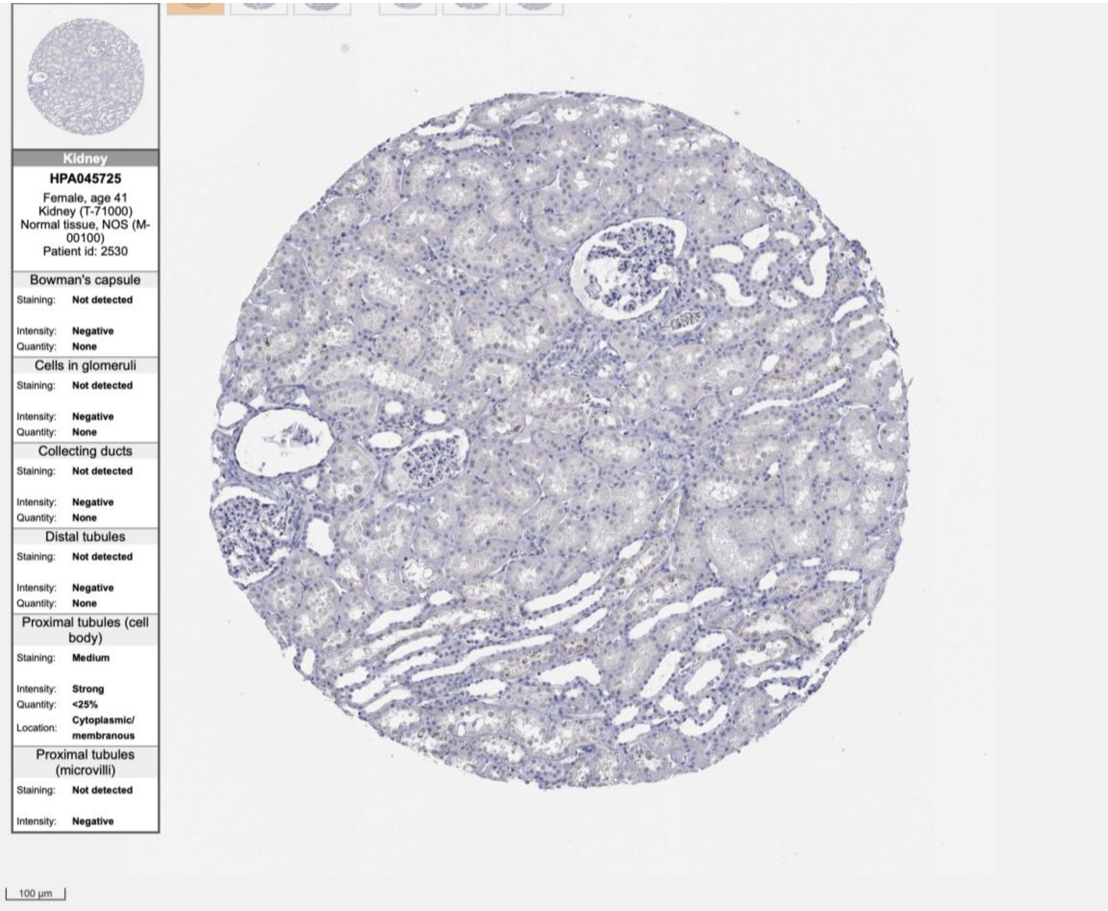

Kidney
HPA045725
Female, age 41
Kidney (T-71000)
Normal tissue, NOS (M-00100)
Patient id: 2530
Bowman's capsule
Staining: Not detected
Intensity: Negative
Quantity: None
Cells in glomeruli
Staining: Not detected
Intensity: Negative
Quantity: None
Collecting ducts
Staining: Not detected
Intensity: Negative
Quantity: None
Distal tubules
Staining: Not detected
Intensity: Negative
Quantity: None
Proximal tubules (cell body)
Staining: Medium
Intensity: Strong
Quantity: <25%
Location: Cytoplasmic/ membranous
Proximal tubules (microvilli)
Staining: Not detected
Intensity: Negative
100 μm


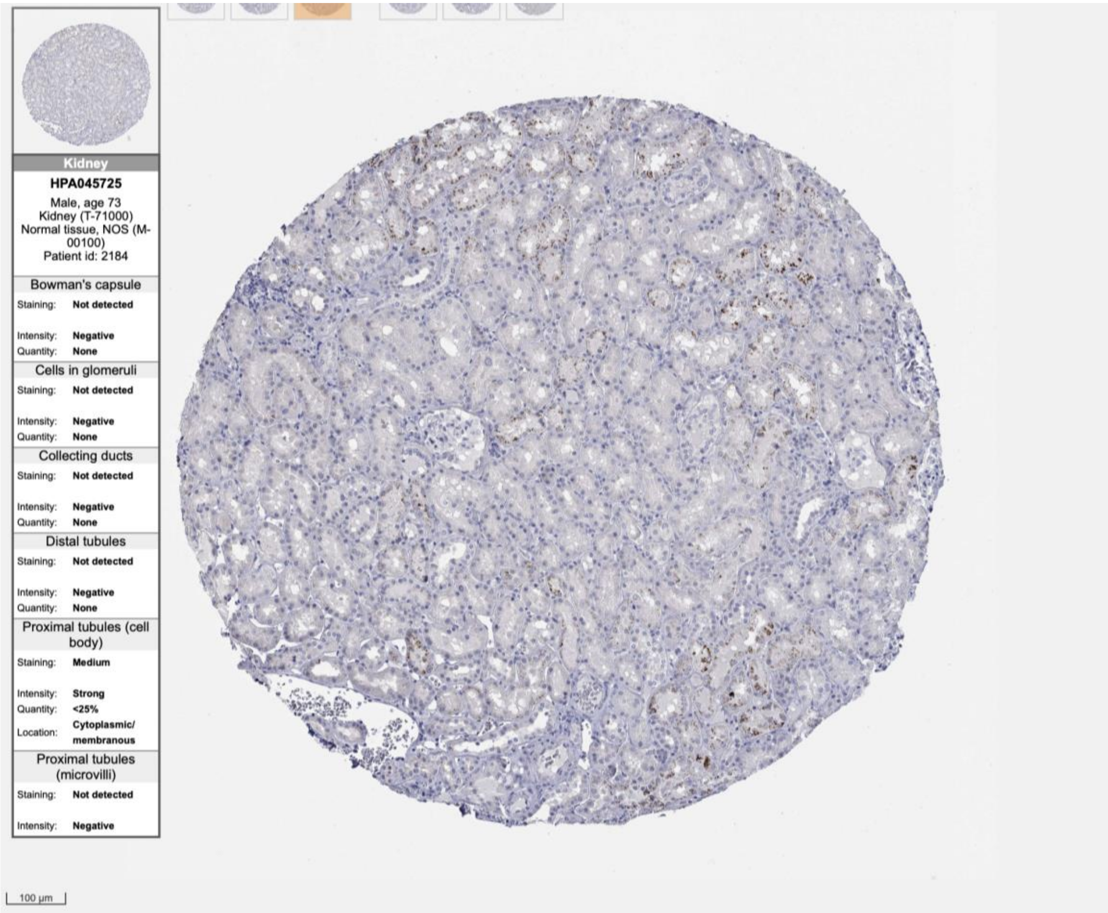

Kidney
HPA045725
Male, age 73
Kidney (T-71000)
Normal tissue, NOS (M-00100)
Patient id: 2184
Bowman's capsule
Staining: Not detected
Intensity: Negative
Quantity: None
Cells in glomeruli
Staining: Not detected
Intensity: Negative
Quantity: None
Collecting ducts
Staining: Not detected
Intensity: Negative
Quantity: None
Distal tubules
Staining: Not detected
Intensity: Negative
Quantity: None
Proximal tubules (cell body)
Staining: Medium
Intensity: Strong
Quantity: <25%
Location: Cytoplasmic/ membranous
Proximal tubules (microvilli)
Staining: Not detected
Intensity: Negative
100 μm

ZNF367

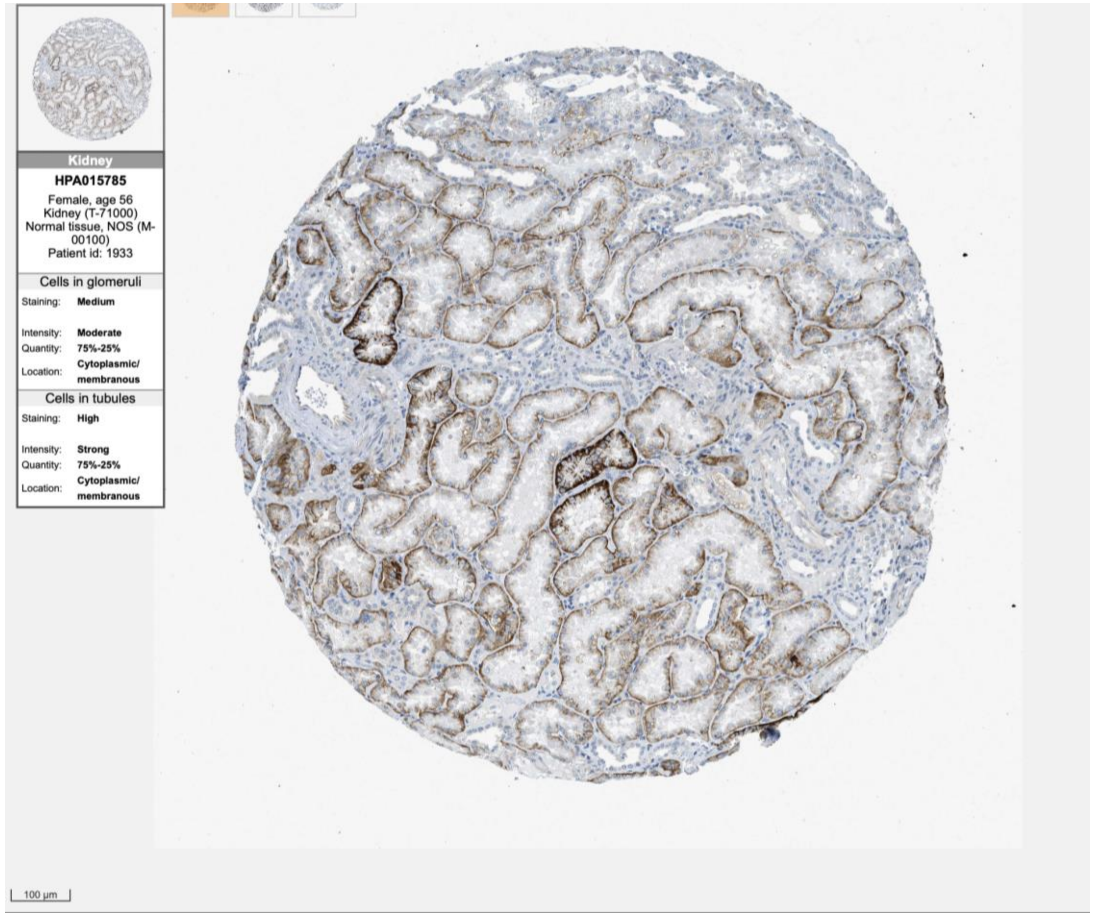
Kidney
HPA015785
Female, age 56
Kidney (T-71000)
Normal tissue, NOS (M-00100)
Patient id: 1933
Cells in glomeruli
Staining: Medium
Intensity: Moderate
Quantity: 75%-25%
Location: Cytoplasmic/ membranous
Cells in tubules
Staining: High
Intensity: Strong
Quantity: 75%-25%
Location: Cytoplasmic/ membranous
100 µm

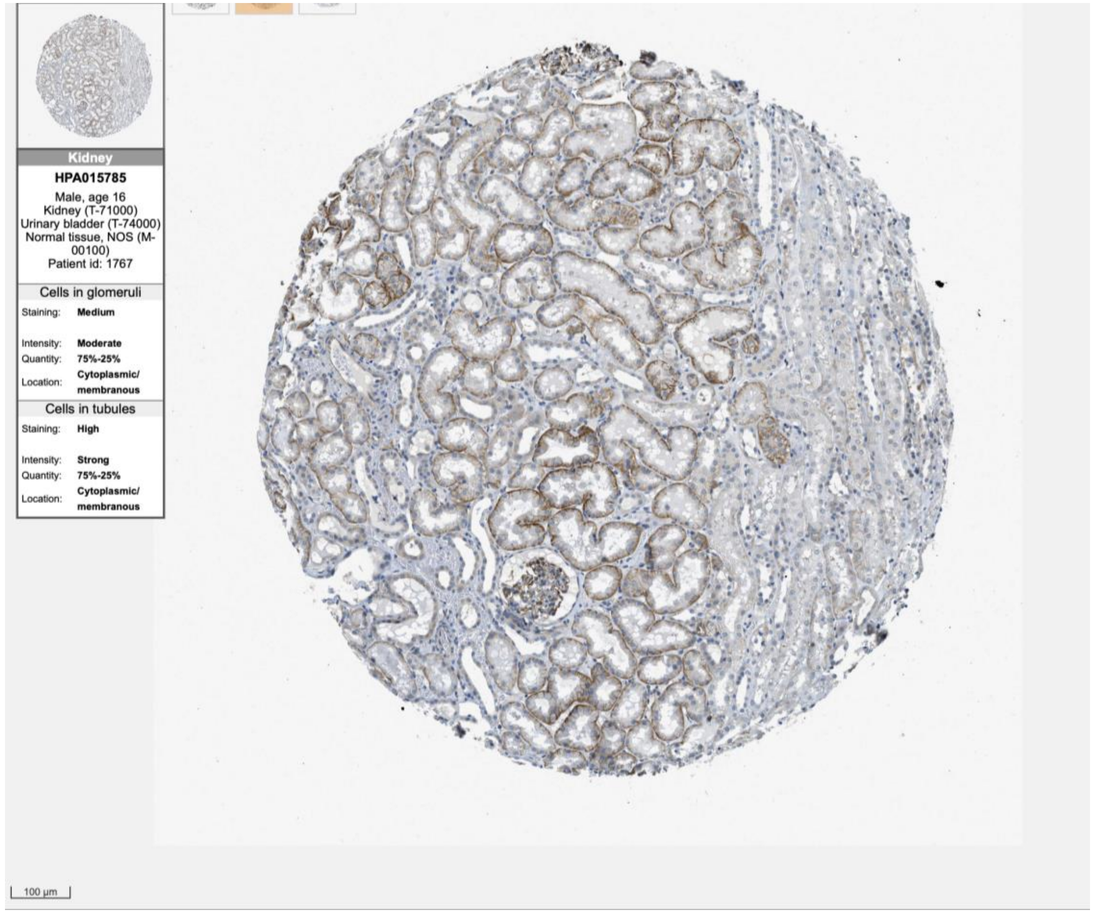
Kidney
HPA015785
Male, age 16
Kidney (T-71000)
Urinary bladder (T-74000)
Normal tissue, NOS (M-00100)
Patient id: 1767
Cells in glomeruli
Staining: Medium
Intensity: Moderate
Quantity: 75%-25%
Location: Cytoplasmic/ membranous
Cells in tubules
Staining: High
Intensity: Strong
Quantity: 75%-25%
Location: Cytoplasmic/ membranous
100 µm
