## Supplementary Figure 8 for "Evidence for conserved gene expression and biological processes operative in human podocytes and adult brain"

A

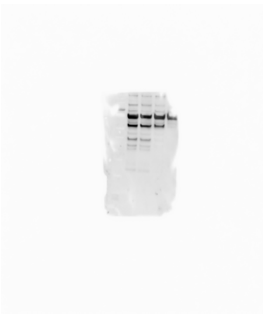

B

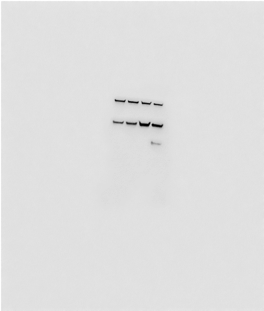

C

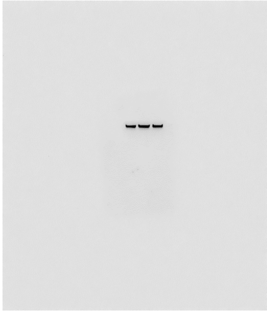

D

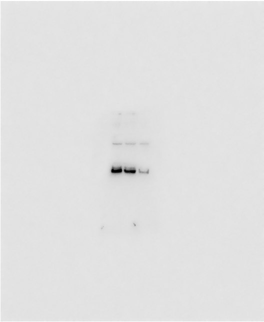

E

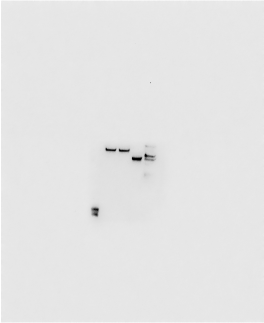

F

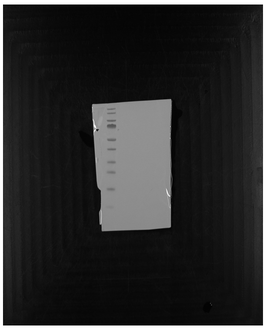

**Supplementary Figure S8: Uncropped western Blot images UF21/UM27.**

The detected proteins are presented from left to right. A: MAP2, B: α-actinin-4 + β-TUBULIN, C: TAU, D+E: GAPDH. Supplementary figure F represents the ladder for the western blot images. The loading scheme starting next to the ladder from the left side as follows: UF21 podocyte and UM27 podocytes. The uncropped western blot images belong to Figure 6A.

A

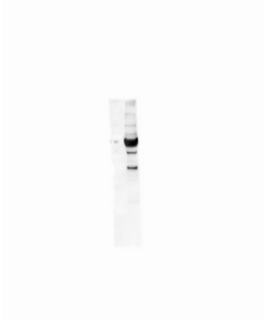

B



C

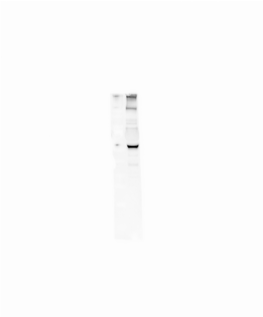

D

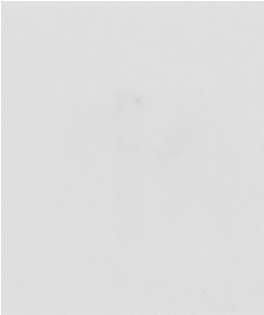

E

F

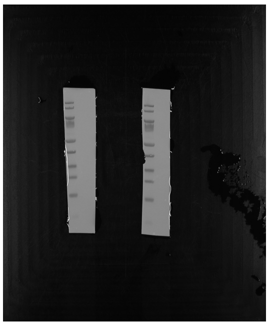

**Supplementary Figure S8: Uncropped western Blot images Human immortalized podocytes (AB 8/13).**

The detected proteins are presented from left to right. A: MAP2, B: α-actinin-4, C: β-TUBULIN, D: TAU, E: GAPDH. Supplementary figure F represents the ladder for the western blot images. The loading scheme starting next to the ladder from the left side as follows: Human immortalized podocytes (AB 8/13). The uncropped western blot images belong to Figure 6B.

A

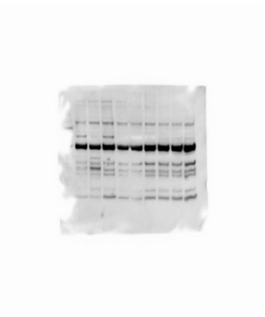

B

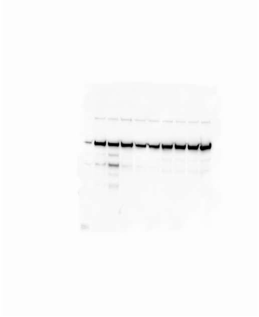

C

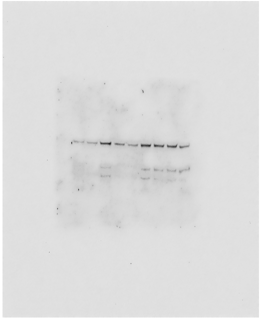

D

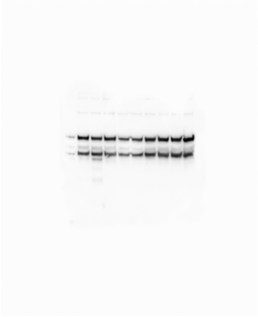

E

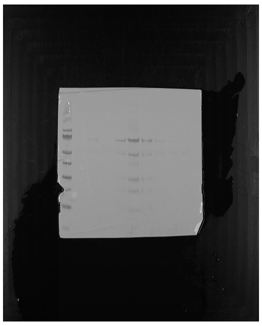

**Supplementary Figure S8: Uncropped western Blot images UM48/UM51 podocytes.**

The detected proteins are presented from left to right. A: MAP2, B: α-actinin-4, C: TAU, D: β-TUBULIN +GAPDH. Supplementary figure E represents the ladder for the western blot images. The loading scheme starting next to the ladder from the left side as follows: UM51 podocytes, UM451 podocytes 24 h ANGII, three rows samples out of context, UM48 podocytes, UM48 podocytes 24 h ANGII, two rows samples out of context. The uncropped western blot images belong to Figure 6C.
